# A divide-and-conquer approach for genomic prediction in rubber tree using machine learning

**DOI:** 10.1101/2022.03.30.486381

**Authors:** Alexandre Hild Aono, Felipe Roberto Francisco, Livia Moura Souza, Paulo de Souza Gonçalves, Erivaldo J. Scaloppi, Vincent Le Guen, Roberto Fritsche-Neto, Gregor Gorjanc, Marcos Gonçalves Quiles, Anete Pereira de Souza

## Abstract

Rubber tree (*Hevea brasiliensis*) is the main feedstock for commercial rubber; however, its long vegetative cycle has hindered the development of more productive varieties via breeding programs. With the availability of *H. brasiliensis* genomic data, several linkage maps with associated quantitative trait loci (QTLs) have been constructed and suggested as a tool for marker-assisted selection (MAS). Nonetheless, novel genomic strategies are still needed, and genomic selection (GS) may facilitate rubber tree breeding programs aimed at reducing the required cycles for performance assessment. Even though such a methodology has already been shown to be a promising tool for rubber tree breeding, increased model predictive capabilities and practical application are still needed. Here, we developed a novel machine learning-based approach for predicting rubber tree stem circumference based on molecular markers. Through a divide-and-conquer strategy, we propose a neural network prediction system with two stages: (1) subpopulation prediction and (2) phenotype estimation. This approach yielded higher accuracies than traditional statistical models in a single-environment scenario. By delivering large accuracy improvements, our methodology represents a powerful tool for use in *Hevea* GS strategies. Therefore, the incorporation of machine learning techniques into rubber tree GS represents an opportunity to build more robust models and optimize *Hevea* breeding programs.

## 1. Introduction

Rubber tree (*Hevea brasiliensis*) has an elevated importance in the global economy, being almost the only feedstock for commercial rubber (Cros et al., 2019; Warren-Thomas et al., 2015). Considering the long perennial vegetative cycle of *Hevea*, breeding programs aim to improve its yield production in order to reach the rapidly increasing rubber demand (Ahrends et al., 2015; Cros et al., 2019; Warren-Thomas et al., 2015). Therefore, genomic approaches are needed in rubber tree breeding, especially considering its recent domestication history (Rosa et al., 2018). *H. brasiliensis* is a diploid species (2*n* = 36) with an elevated occurrence of duplicated regions in its genome (~ 70%) (Lau et al., 2016; Liu et al., 2020; Tang et al., 2016), and this complex genomic organization has hindered the development of genomic strategies for breeding. However, with the improvement of next-generation sequencing (NGS) technologies and the consequent reduction in genotyping costs, data generation has become more efficient, providing more genomic resources in less time and with lower associated costs (Roorkiwal et al., 2018). This greater availability of data improved precision in selection with higher genetic gains in various crops (González-Camacho et al., 2018; Roorkiwal et al., 2018) and, in rubber tree, could complement traditional approaches based on only phenotypic and pedigree information (Hayes et al., 2013; Roorkiwal et al., 2018).

Various rubber tree genomic resources have become available in recent decades, such as a large set of different molecular markers (Lespinasse et al., 2000b; Nakkanong et al., 2008; de Souza et al., 2016; Venkatachalam et al., 2006), draft genomes (Lau et al., 2016; Tang et al., 2016), and, more recently, a chromosome-level assembled genome (Liu et al., 2020). These data have already allowed the construction of saturated linkage maps with associated quantitative trait loci (QTLs), which were proposed as a tool for marker-assisted selection (MAS) (An et al., 2019). Although QTLs for several traits have been identified in rubber tree (An et al., 2019; Le Guen et al., 2011, 2007; Lespinasse et al., 2000a; Rosa et al., 2018; Souza et al., 2013; Tran et al., 2016), the amount of phenotypic variance explained by these identified QTLs is usually small (Souza et al., 2013) because of the highly complex genetic architectures associated with growth and rubber production traits. The configuration of these phenotypes is controlled by many genes with small effects (Washburn et al., 2019), and weak QTLs may not be identified using existing methodologies (Cros et al., 2019; Muranty et al., 2015), which prevents the identification of interindividual differences (Bellot et al., 2018). Together with the environmental and genetic background restrictions of QTLs (Crossa et al., 2017), these features limit the application of *Hevea* QTLs for MAS (de Souza et al., 2016). Consequently, novel genomic strategies that can assist in rubber tree breeding programs are needed, especially considering the time required to evaluate these phenotypes, the elevated costs, and the low female fertility in *H. brasiliensis* (An et al., 2019; Cros et al., 2019; Souza et al., 2019).

Aimed at solving such difficulties in many crops, genomic selection (GS) has arisen as a promising methodology for considerably reducing the required breeding cycle (Hayes et al., 2001). GS has shown better performance than MAS (Bernardo & Yu, 2007; Heffner et al., 2010), mainly because of its associated genetic gains (Albrecht et al., 2011) and reduced costs over a long time period (Wang et al., 2018). This strategy enables the selection of plants based on their estimated performance obtained with a large dataset of molecular markers (Ma et al., 2018; Roorkiwal et al., 2018), reducing breeding time by avoiding the need to evaluate a considerable number of phenotypes over different years (Crossa et al., 2017). Using known phenotypic and genotypic information from a training population (Crossa et al., 2019), it is possible to create a predictive model that can be used to predict the breeding values of a testing population using only genotypic data (Roorkiwal et al., 2018). This modeling is generally based on a mixed-effect regression method (Montesinos-López et al., 2018) and has already been demonstrated to be promising for several crops (Crossa et al., 2016; Spindel et al., 2015; Wolfe et al., 2017; Xavier et al., 2016; Zhao et al., 2012). In rubber tree, Souza et al. (2019) and Cros et al. (2019) assessed the potential of GS for predicting stem circumference (SC) and rubber production (RP), respectively, simulating breeding schemes through cross-validation (CV) techniques.

There are several CV approaches for simulating a real application of GS in a plant breeding program. These methods take into account the population structure in the dataset and the appropriateness of applying the developed predictive model to a set of plants. There are basically three approaches, which are used to (1) predict traits in an untested environment using previously tested lines (CV0) (Roorkiwal et al., 2018), (2) predict new lines’ traits that were not evaluated in any environment (CV1) (Montesinos-López et al., 2019b), and (3) predict traits that were evaluated in some environments but not in others (CV2) (Jarquín et al., 2017). These three scenarios were already evaluated in rubber tree. Cros et al. (2019) assessed the potential of GS in a within-family context using CV0 and CV1 methods, and Souza et al. (2019) tested three different populations with CV1 and CV2. These initiatives represent the first attempts to use GS on rubber tree data, but with low associated predictive capabilities for some of the created CV schemes, mostly when prediction is performed with genotypes that have not already been tested.

Different approaches have been used in GS to create predictive models, including parametric and nonparametric methods (Crossa et al., 2017; De Los Campos et al., 2009; Endelman, 2011; Hayes et al., 2001; Jannink et al., 2010; VanRaden, 2007, 2008). Significant differences in predictive capabilities have not been demonstrated when changing the predictive approach (Ma et al., 2018; Roorkiwal et al., 2016; Varshney, 2016); thus, linking genotypes and phenotypes remains a great challenge (Bellot et al., 2018; Harfouche et al., 2019), especially for plant species with high genomic complexity. In this context, more robust techniques for estimating these models with higher prediction capabilities are needed to expand the practical implementation of GS in rubber tree. Nonlinear techniques have already shown improved performance in representing complex traits with nonadditive effects (Crossa et al., 2014; González-Camacho et al., 2012, 2018; Pérez-Rodríguez et al., 2012), and, in this context, machine learning (ML) strategies have emerged as a promising set of tools for complementing these statistical nonlinear methods.

The objective of this work was to develop a genomic prediction approach for rubber tree data. Considering that ML methods have not been proven to have better performance than statistical methodologies for GS (Bellot et al., 2018; Montesinos-López et al., 2019a), we evaluated their efficiency in rubber tree, also suggesting a novel approach for constructing a predictive system with neural networks based on two-stage prediction: (1) subpopulation prediction and (2) phenotype estimation. Such a divisive approach was created considering a common paradigm in Computer Science: divide and conquer. For datasets with a clear subpopulation structure, such as rubber tree, the proposed approach represents a promising alternative for the development of predictive models.

## 2. Material and methods

### 2.1. Plant material and phenotypic characterization

The data used in this work were obtained with different experiments in two previous studies. Therefore, our analyses were conducted by separating the methodologies and considering two datasets: experimental group 1 (EG1) and experimental group 2 (EG2). EG1 includes 408 samples of three F1 segregant populations obtained with crosses between (Pop1) GT1 and PB235 (30 genotypes) (Souza et al., 2019), (Pop2) GT1 and RRIM701 (127 genotypes) (Conson et al., 2018; Souza et al., 2019), and (Pop3) PR255 and PB217 (251 genotypes) (Rosa et al., 2018; Souza et al., 2013, 2019). EG2 is based on an F1 cross between RRIM600 and PB260 (330 samples) (Cros et al., 2019).

The parents of the crosses used are important clones for rubber tree breeding programs. PR255, PB235, PB260, and RRIM600 have high yield, and PB217 has considerable potential for long-term yield performance due to its slow growth process (Cros et al., 2019; Souza et al., 2019). PR255 and RRIM701 have good growth, and RRIM701 also presents an increased SC after initial tapping (Romain & Thierry, 2011). The latex production is stable in PR255 and medium in RRIM600. Stable or medium latex production represents a good adaptation to several environments, as observed in GT1, a clone tolerant to wind and cold. Additionally, PB260 presents high female fertility (Baudouin et al., 1997), and PB235 is susceptible to tapping panel dryness (Sivakumaran et al., 1988).

In EG1 and EG2, we analyzed the SC trait. In EG1, Pop3 was planted in 2006 in a randomized block design in Itiquira, Mato Grosso State, Brazil, 17°24′ 03′′ S and 54°44′ 53′′ W (Rosa et al., 2018; Souza et al., 2013, 2019). Each individual was represented by four grafted trees in each plot and four replications. Pop1 and Pop2 were planted in 2012 at the Center of Rubber Tree and Agroforestry Systems/Agronomic Institute (IAC - Brazil), 20°25′ 00′′ S and 49°59′ 00′′ W, following an augmented block design, with four blocks containing two clones per plot spaced 4 m apart for each trial, which was repeated four times (Conson et al., 2018; Souza et al., 2019).

Even though EG2 corresponds to only one cross, this population was planted following an almost complete block design at two different sites (Cros et al., 2019), which for convenience we named site 1 (S1) and site 2 (S2). In S1, 189 clones were planted in 2012 in Société des Caoutchoucs de Grand-Béréby (SOGB - Ivory Coast), 4°40′ 54′′ N and 7°06′ 05′′ W. In S2, 143 clones were planted in 2013 in Société Africaine de Plantations d’Hévéas (SAPH - Ivory Coast), 5°19′ 47.79′′ N and 4°36′ 39.74′′ W. This cross consisted of six blocks with randomized trees spaced 2.5 m apart and a mean number of ramets per clone of 11 for S1 (ranging between 7 and 17) and 13 for S2 (ranging between 5 and 20).

SC measurements of Pop3 in EG1 were obtained in four years (from 2007 to 2010) and those of Pop1 and Pop2 were obtained from 2013 to 2016, considering that growth traits are usually measured only during the first 6 years (Rao & Kole, 2016; Souza et al., 2019). According to the water distribution of the experiments installed, EG1 phenotypes were measured to supply information considering low-water (LW) and well-watered (WW) conditions; thus, Pop3 was evaluated in October 2007-2010 (LW) and in April 2008-2010 (WW), and Pop1 and Pop2 were evaluated in June 2013, December 2013, May 2014, November 2014, and June 2015-2016. SCs were measured for individual trees at 50 cm above ground level. For both phenotypes, the average per plot was calculated. SC in EG2 was measured at 1 m above ground level before tapping for 3 months every two days except on Sundays (with the beginning at 32 months after planting in S1 and 38 months after planting in S2).

### 2.2. Phenotypic data analysis

All phenotypic analyses were performed using R statistical software (Team et al., 2013). EG1 and EG2 traits were analyzed with the following steps: (1) data distribution evaluation; (2) standardized normalization with the R package bestNormalize (Peterson, 2017); (3) mixed-effect model creation and residual appropriateness verification through quantile-quantile (Q-Q) plots using the breedR package (Muñoz & Sanchez, 2019); (4) estimation of best linear unbiased predictions (BLUPs) based on the models created; (5) hierarchical clustering on BLUP values using a complete hierarchical clustering approach based on Euclidean distances and dendrogram visualization with the ggtree R package (Yu et al., 2017); and (6) identification of phenotypic groups using the clustering approach of (5), with cluster numbers ranging between 2 and 5, and several clustering indexes implemented in the NbClust R package (Charrad et al., 2014).

In EG1, we employed the following statistical mixed-effect model:

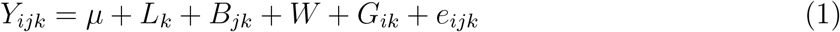

where *Y*_*ijk*_ corresponds to the phenotype of the *i*th genotype in the *j*th block and *k*th location. The phenotypic mean is represented by *μ*, and the fixed effects represent the contribution of the *k*th location (*L*_*k*_), the *j*th block at the *k*th location (*B*_*jk*_), and the watering condition of the measurement (*W*). The genotype *G* and the residual error *e* (nongenetic effects) represent the random effects.

EG2 SC phenotypes were modeled for each site (S1 and S2) according to the following statistical model:

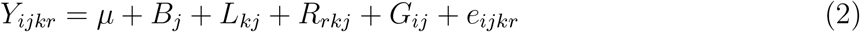

where *Y*_*ijkr*_ corresponds to the phenotype of the *i*th genotype positioned in the *r*th rank of the *k*th line in the *j*th block. The phenotypic mean is represented by *μ*, and the fixed effects represent the contribution of the *j*th block (*B*_*j*_), the *k*th line of the *j*th block (*L*_*kj*_), and the *r*th rank of the *k*th line in the *j*th block (*R*_*rkj*_). The genotype *G* and the residual error *e* (nongenetic effects) represent the random effects. Broad-sense heritability (*H*^2^) was estimated as 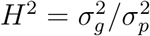, with 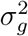 and 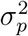 representing the genetic and phenotypic variances, respectively.

### 2.3. Genotyping process

DNA extraction from EG1 was described by Conson et al. (2018); Souza et al. (2013), and the genotyping process was performed using a genotyping-by-sequencing (GBS) protocol (Elshire et al., 2011) with *EcoT22I* restriction enzyme followed by Illumina sequencing using the HiSeq platform for Pop3 and the GAIIx platform for Pop1 and Pop2 (Souza et al., 2019). Raw sequencing reads were processed using the TASSEL 5.0 pipeline (Glaubitz et al., 2014), with a minimum count of 6 reads for creating a tag. The tag mapping process was performed using Bowtie2 v.2.1 (Li & Durbin, 2009) with the *very sensitive* algorithm and *H. brasiliensis* reference genome (Liu et al., 2020). Single nucleotide polymorphisms (SNPs) were called with the TASSEL algorithm, and only biallelic SNPs were retained using VCFtools (Danecek et al., 2011). These markers were filtered using the R package snpReady (Granato et al., 2018b) with a maximum of 20% missing data for a SNP and 50% in an individual and a minimum allele frequency (MAF) of 5%. Missing data were imputed using the k-nearest neighbors (Cover & Hart, 1967) algorithm implemented in the snpReady package.

EG2 samples were genotyped with simple sequence repeat (SSR) markers, following the protocol for DNA extraction and genotyping described by Le Guen et al. (2009). A total of 332 SSRs were used for S1 (Tran et al., 2016) and 296 for S2 (Cros et al., 2019). Missing data were imputed using BEAGLE 3.3.2 (Browning & Browning, 2007) with 25 iterations of the phasing algorithm and 20 haplotype pairs to sample for each individual in an iteration. The genotypic profile of individuals in EG1 and EG2 was evaluated using principal component analyses (PCAs) in R statistical software (Team et al., 2013) with the ggplot2 package (Wickham, 2016).

### 2.4. Statistical models for genomic prediction

We employed two different strategies for creating traditional genomic prediction models: Bayesian ridge regression (BRR) (Gianola, 2013) and a single-environment, main genotypic effect model with a Gaussian kernel (SM-GK) (Cuevas et al., 2016). BRR and SM-GK models were implemented in the BGLR (Pérez & de Los Campos, 2014) and BGGE (Granato et al., 2018a) R packages, respectively. Considering the genotype matrix with *n* individuals and *p* markers, BRR models were implemented considering the following:

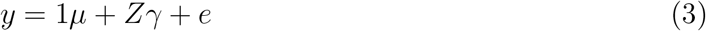

where *y* represents the BLUP values calculated based on the established mixed-effect models for phenotypic data analyses, *μ* the overall mean, *Z* the genotype matrix, *e* the residuals, and *γ* the vector of marker effects. In SM-GK, *Z* is the incidence matrix of genetic effects, and *γ* is the vector of genetic effects with variance estimated through a Gaussian kernel calculated using the snpReady R package.

### 2.5. Genomic prediction via machine learning

For genomic prediction via ML, we selected the following algorithms: (a) AdaBoost (Freund & Schapire, 1997), (b) multilayer perceptron (MLP) neural networks (Popescu et al., 2009), (c) random forests (Breiman, 2001), and (d) support vector machine (SVM) (Shawe-Taylor & Cristianini, 2000). To create these models, we used Python v.3 programming language together with the library scikit-learn v.0.19.0 (Pedregosa et al., 2011). We also tested a combination of feature selection (FS) techniques for increasing the predictive accuracies (Aono et al., 2020), using a combination of three different methods: (i) L1-based FS through an SVM model (Shawe-Taylor & Cristianini, 2000), (ii) univariate FS with Pearson correlations (and ANOVA for discrete variables) (p-value of 0.05), and (iii) gradient tree boosting (Chen & Guestrin, 2016). Such a strategy is based on marker subset selection, separating the markers identified by all of these methods together (intersection of the 3 approaches, named Inter3) or by at least two of them simultaneously (Inter2), and using such subsets for prediction.

To understand the subset selection, we performed functional annotation of the genomic regions underlying these markers selected through FS considering a 10,000 base-pair (bp) window for the up- and downstream regions. Using BLASTn software (Altschul et al., 1990) (minimum e-value of 1e-6), these sequences were aligned against coding DNA sequences (CDSs) from the *Malpighiales* clade (*Linum usitatissimum* v1.0, *Manihot esculenta* v8.1, *Populus deltoides* WV94 v2.1, *Populus trichocarpa* v4.1, *Ricinus communis* v0.1, and *Salix purpurea* v5.1) of the Phytozome v.13 database (Goodstein et al., 2012). On the basis of significant correspondence, Gene Ontology (GO) terms (Botstein et al., 2000) were retrieved.

### 2.6. Multilayer perceptron neural network

As the final approach for genomic prediction in EG1, we proposed the creation of neural networks with novel architectures for each of the biparental populations, using the Keras Python v.3 library for this task (Chollet et al., 2015). We employed MLP networks, which have an architecture based on multiple layers and feedforward signal propagation (Da Silva et al., 2017). The MLP architecture is organized into one input layer (IL), followed by at least one hidden layer (HL) and one output layer (OL). Each one of these layers contains processing elements, named neurons, which are interconnected with associated unidirectional numeric values (weights) (Hecht-Nielsen, 1992). The number of neurons in the IL corresponds to the quantity of explanatory (independent) variables of the problem, which will be propagated across the MLP structure in one direction (from the input to the output) (Da Silva et al., 2017). The HLs receive the output of the previous layer until this feedforward propagation generates the OL, respecting the established connections and weights of the architecture. HLs are included in an MLP to extract unknown patterns from the dataset, making decisions that will contribute to the overall prediction process (Da Silva et al., 2017; O’Shea & Nash, 2015). After the HLs, the architecture contains the OL, which is related to the response (dependent) variable of the problem. For regression tasks with a single output, there is only one neuron in the OL with linear values (Kurková & Sanguineti, 2013).

Each neuron in an MLP has an output value corresponding to impulses that will be propagated into the network. The input signals (*x*_1_, *x*_2_, …, *x*_*n*_) of a neuron are multiplied by synaptic weights (*w*_1_, *w*_2_, …, *w*_*n*_) representing their importance in neuron activation (Da Silva et al., 2017). The results of these multiplications are aggregated through summation and subtracted by an activation threshold/bias (*θ*). Thus, an output signal is produced, whose value is limited with the use of an activation function *g*, e.g., rectified linear activation (ReLU), logistic, arc tangent, and hyperbolic tangent functions. The purpose of such functions is to introduce non-linearity into the network (Wang, 2003). The output *s* of an HL neuron can then be summarized in (Da Silva et al., 2017)

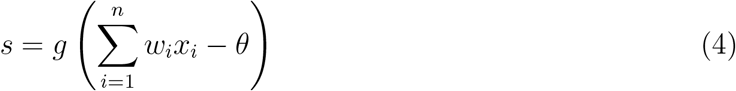

The structure of an artificial neural network is adaptive, changing its conformation during a process called training, which aims to reach stability in the network via minimal error in predictive performance through changes in the connection weights (Sheela & Deepa, 2013). The synaptic weights in an MLP are adjusted by measuring the predictive performance of the architecture via an error function, such as the sum of squared errors (Wang, 2003). Even though the propagation of signals in an MLP is in the forward direction, adjustments of weights are not propagated in these feedforward connections. Based on the comparison of the network output with the desirable response and the obtainment of an error value, the weights are updated in backward propagation (Da Silva et al., 2017) to minimize the found error using this backpropagation strategy together with an optimization algorithm (Hecht-Nielsen, 1992; Rumelhart, 1986), such as stochastic gradient descent (SGD), adaptive moment estimation (Adam) (Kingma & Ba, 2014), and Rmsprop (Bengio, 2015). This process is repeated using the training data in a number of cycles (Hoffer et al., 2017), named epochs, and this backpropagation strategy usually employs a batch of samples at each gradient computation for updating the weights (Hoffer et al., 2019).

For all the predictive tasks, we considered an MLP structure with two HLs and used the mean absolute error (MAE) as the error function for training and defining the architecture of the networks. Additionally, 200 epochs were considered (batch size of 16). The training process of the networks was performed using the backpropagation strategy together with the Adam optimization algorithm (Kingma & Ba, 2014), which aims to minimize the MAE by updating the synaptic weights using a gradient-based strategy that combines heuristics from a momentum term and RMSProp (Bengio, 2015). The update process is based on a change of Δ*w*_*ij*_ for each connection, considering the individual influence of a weight *w*_*ij*_ on the MAE value obtained with the gradient descent *g*_*t*_ in the iteration *t* calculated with *∂MAE/∂w*_*ij*_ and used in the equation

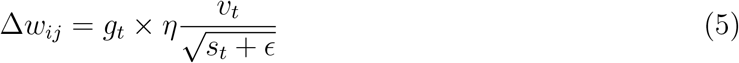

where *η* is the learning rate representing the amount of change in the process of training, *v*_*t*_ is the exponential average of gradients along the weights *w*_*i*_ of layer *i*, and *s*_*t*_ is the exponential average of squares of gradients along *w*_*i*_. The Adam optimizer employs two other hyperparameters for the optimization process (*β*_1_ and *β*_2_), which are used for the calculation of *v*_*t*_ (*v*_*t*_ = *β*_1_ × *v*_*t*−1_ − (1 − *β*_1_) × *g*_*t*_) and 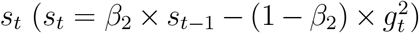. We used *β*_1_ = 0.9 and *β*_2_ = 0.999 (Kingma & Ba, 2014). We tested the following configurations for the MLP hyperparameters: (a) number of neurons in the first HL, varying from 1 to 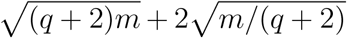 (*m* individuals and *q* output neurons in the OL); (b) number of neurons in the second HL, varying from 1 to 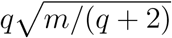); (c) ReLU, sigmoid and hyperbolic tangent activation functions; and (d) learning rates of 0.005, 0.001, and 0.0001.

### 2.7. Proposed approach and validation strategies

Each of the sets of hyperparameters estimated for the MLP networks was used to create a joint and single system for prediction in EG1, which we indicate as part of a divide-and-conquer approach created for genomic prediction (Fig. 1). Considering an individual as part of a dataset subpopulation that has a specific phenotypic distribution, we propose the use of a two-stage prediction process based on the following steps: (1) creating four different neural networks according to different hyperparameter searches and the training data (division step), (2) predicting which subpopulation an unlabeled observation belongs to according to the network induced for this task (prediction 1 and conquer step), and (3) predicting its phenotypic performance based on the network trained specifically for the subpopulation predicted (prediction 2 and final conquer step).

**Fig. 1.**
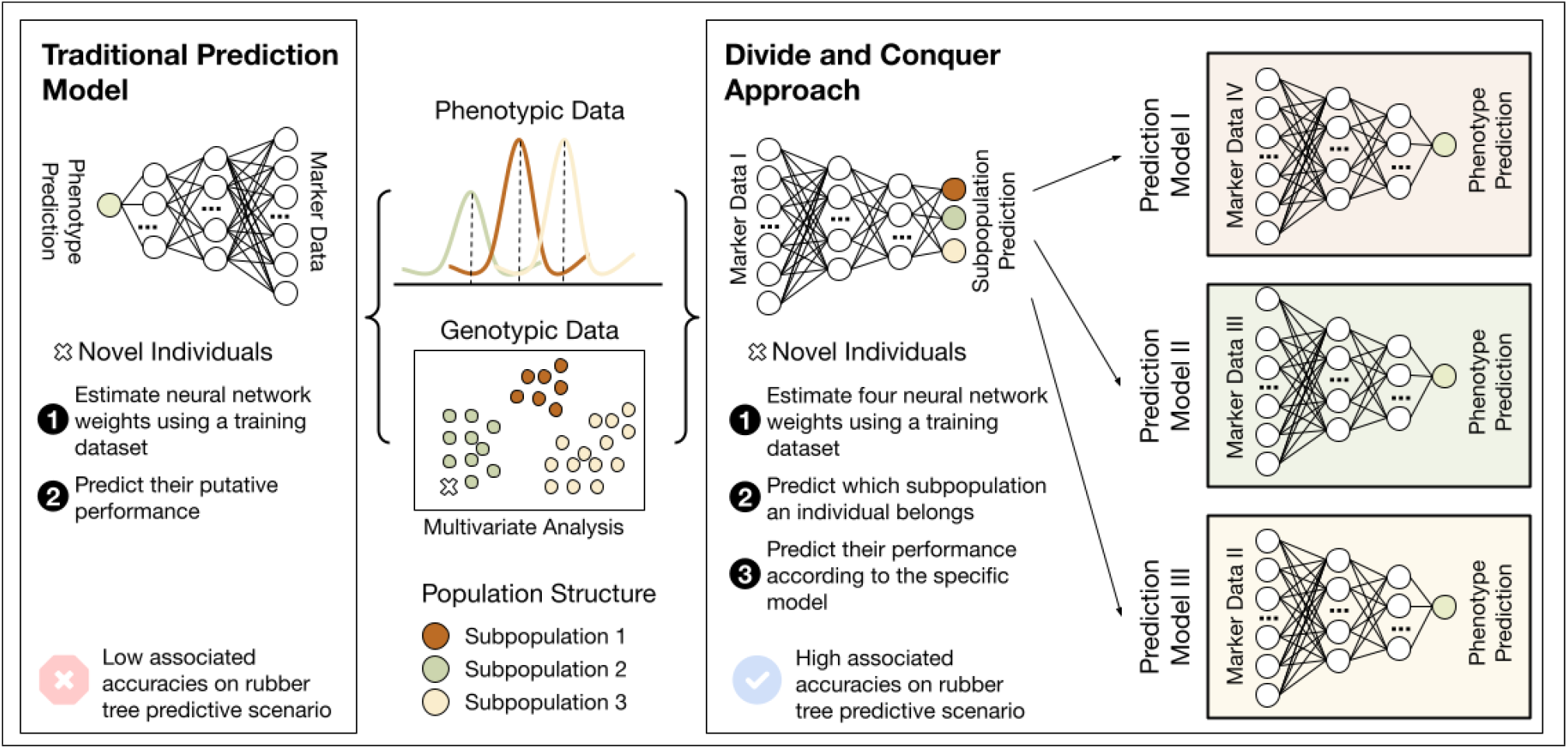
Overview of the approach proposed. Based on a divide-and-conquer strategy with different neural networks combined into a single model (part 1), individuals with unknown phenotypic performance (a) are classified into a subpopulation using a specific neural network (part 2) and (b) have their phenotypic values estimated through an induced network specific to the subpopulation they belong to (part 3).

CV1 was the strategy employed for the selection of data for evaluating the models’ performance due to its reduced bias when splitting the dataset and the low prediction accuracies described (Souza et al., 2019). We first separated a test dataset using 10% of the genotypes with a stratified holdout strategy implemented in the scikit-learn Python v.3 module (Pedregosa et al., 2011). The stratification was performed only in EG1 and was based on the subpopulation structure present in the dataset. For all the models evaluated in this work (statistical and ML based), the same dataset split was considered in every round of CV.

The remaining 90% of the genotypes were used as the development set for defining the networks’ architecture and for evaluating the overall models’ performance through a stratified k-fold approach (k=4) with 50 repetitions (subpopulation stratification). The predictive accuracy in every CV split was evaluated by comparing the predicted and real BLUPs by measuring (1) the Pearson correlation coefficient (*R*) and (2) the mean absolute percentage error (MAPE). For each trait, we compared the predictive accuracy differences using ANOVA and multiple comparisons by Tukey’s test with the agricolae R package (de Mendiburu & de Mendiburu, 2019).

For EG1, four different MLP architectures were estimated: (a) subpopulation prediction, (b) BLUP prediction for Pop1, (c) BLUP prediction for Pop2, and (d) BLUP prediction for Pop3. After defining the network hyperparameters with the development set, all of these structures were joined into a single predictive system that was used for the final prediction. In addition to evaluating the predictive performance through the CV scenarios created, we also checked the performance of the model for a leave-one-out (LOO) CV configuration.

## 3. Results

### 3.1. Phenotypic and genotypic data analysis

The raw phenotypic data were evaluated considering the experimental groups proposed. EG1 (Supplementary Fig. 1) had reduced values compared to those of EG2 (Supplementary Fig. 2) due to the different heights and years of stem measurements. However, for the normalized SC values (Supplementary Figs. 3-5), such an evident discrepancy was not observed. By modeling the phenotypic measures with the mixed-effect models established and contrasting the raw values with the normalized ones through Q-Q plots, we observed that the residuals obtained with the normalized measurements in EG1 (Supplementary Fig. 6) and EG2 (Supplementary Figs. 7-8) were more appropriate. Heritabilities (*H*^2^) were estimated as 0.55 for EG1, 0.83 for EG2-S1 and 0.93 for EG2-S2, which is in accordance with the findings of Souza et al. (2019) and Cros et al. (2019).

Interestingly, BLUPs from EG1 (Supplementary Fig. 9) and EG2-S1 (Supplementary Fig. 10) presented reduced variability when compared to that of BLUPs estimated for EG2-S2 (Supplementary Fig. 10). This observation is corroborated by the hierarchical clustering analyses performed for these experimental groups. EG1 (Supplementary Fig. 11) and EG2-S1 (Supplementary Fig. 12) could be divided into three phenotypic groups according to the best data partitioning scheme established through NbClust clustering indexes (Charrad et al., 2014), and EG2-S2 could be arranged into 5 such groups (Supplementary Fig. 13). Therefore, it was expected that for the genomic prediction step, EG2-S2 would represent a more difficult task due to its higher data variability.

SNP calling in EG1 was performed according to the TASSEL pipeline. Of the 363,641 tags produced, approximately 84.78% could be aligned against the *H. brasiliensis* reference genome, which generated 107,466 SNPs. These markers were filtered separately for each population using the parameters established, and then these separated datasets were combined through intersection comparisons, yielding a final dataset of 7,414 high-quality SNP markers. For EG2 predictions, 332 and 296 SSR markers were used for EG2-S1 and EG2-S2, respectively.

Using these datasets, we performed PCAs for EG1 (Supplementary Fig. 14) and EG2 (Supplementary Fig. 15). In the figures, the colors of the genotypes correspond to their BLUP values, and their shapes correspond to population structure in EG1 and site in EG2. As expected, for the SC trait, there were no clear associations between markers and BLUPs, underlining the challenge of creating genomic prediction models. Additionally, the subpopulation structure in EG1 was evident.

### 3.2. Genomic prediction

From the BLUP and marker datasets, we fit genomic prediction models using the traditional statistical approaches (BRR and SM-GK) and the ML algorithms (AdaBoost, MLP, RF, and SVM) selected. For EG1 (Supplementary Fig. 16), EG2-S1 (Supplementary Fig. 17) and EG2-S2 (Supplementary Fig. 18), no substantial changes were observed when changing the prediction approach. After applying Tukey’s multiple comparisons test, we found equivalent performance values for SVM, SM-GK and BRR for all the experimental groups. The worst performance was observed for MLP, however, considering the default architectures employed in scikit-learn (Pedregosa et al., 2011).

Additionally, we also tested the inclusion of FS techniques for increasing model performance in ML algorithms. Using the Inter2 approach, we selected 539 (~7.27%), 69 (~20.78%) and 82 (~27.70%) markers for EG1, EG2-S1 and EG2-S2, respectively. For Inter3, 113 (~1.52%), 8 (~2.41%) and 15 (~5.07%) markers were identified. This SNP subsetting approach was beneficial for EG1 (Supplementary Fig. 19A), EG2-S1 (Supplementary Fig. 20) and EG2-S2 (Supplementary Fig. 21); however, there were less pronounced improvements for data from EG2 sites, which was expected because of the limited SSR marker dataset. We considered that, even with increased predictive accuracies, to achieve better results, a wider set of markers would be required. Then, we considered the best strategy for EG2-S1 to be the combination of the Inter2 FS approach with SVM and that for EG2-S2 to be the combination of Inter3 FS with the AdaBoost ML algorithm.

Even though FS approaches boosted prediction accuracies for EG1, when analyzing model performance by calculating the Pearson correlation between the real and predicted BLUPs for each family separately, this better performance was caused by the overall predictions. However, when analyzing predictive power within families (Supplementary Fig. 19B), such an approach was not sufficient for obtaining a reliable prediction with this evident data stratification. In this context, different from EG2, we developed an approach specific to datasets similar to EG1, i.e., a methodology with high capabilities to supply accurate predictions, even considering the subpopulation structure present in a dataset.

Considering a genomic prediction problem based on the creation of a regression model for a dataset containing genotypes that belong to different groups of genetically similar individuals, we modeled such a task by dividing the prediction into different stages (Fig. 1) and creating a divide-and-conquer approach for prediction. The basis of such an approach is that closely related genotypes will share QTLs that might not be the same in another group of genotypes. Therefore, we created a different neural network for each biparental population (divide part), coupled with an intrapopulation system of FS and with a different form of hyperparameter estimation. Following this division part, the separated systems were combined using an additional step (the conquer part). To do so, another neural network was created to infer which subpart of the system should be used for prediction.

### 3.3. Feature selection at the subpopulation level

The selection of subsets of markers was performed according to each EG1 network using the four different tasks: (i) subpopulation prediction, (ii) EG1-Pop1 BLUP prediction, (iii) EG1-Pop2 BLUP prediction, and (iv) EG1-Pop3 BLUP prediction. As expected, each FS strategy returned a different quantity of markers (Table 1). For each subset of markers selected considering Inter2 and Inter3, we evaluated their performance using the ML algorithms selected. Some of the models created for task (i) did not present any mistakes (Supplementary Fig. 22), which was expected due to the subpopulation structure present in the dataset and their evident linear separability. For this task, we considered the most suitable FS strategy to be the Inter2 approach.

**Table 1.** Feature selection strategies performed on the marker dataset considering the intersection among the three methods established (Inter3) and the intersection among at least two out of the three methods established (Inter2).

| Prediction Scenario | Inter2 | Inter3 |
| --- | --- | --- |
| Subpopulation Prediction | 224 | 17 |
| GT1 x PB235 | 345 | 20 |
| GT1 x RRIM701 | 454 | 62 |
| PR255 x PB217 | 591 | 119 |

For EG1-Pop1 (Supplementary Fig. 23), EG1-Pop2 (Supplementary Fig. 24) and EG1-Pop3 (Supplementary Fig. 25), the best accuracies were observed for the combination Inter2-SVM. However, considering the overall performance with the other algorithms, the best approach for SNP subsetting was Inter3. For this reason, we selected this strategy for the BLUP prediction task. Interestingly, there was no intersection between these three Inter3 datasets in the populations; the only case of overlap was a single SNP marker in Pop2 and Pop3.

From the genomic regions flanking these markers selected for BLUP prediction, we could retrieve several instances of correspondence between rubber tree sequences and CDSs from the *Malpighiales* clade in the Phytozome database. From the 20 markers used in Pop1 for prediction, 62 in Pop2, and 119 in Pop3, we found CDS correspondence for the genomic regions related to 8 (40%), 27 (~43.55%) and 48 (~40.32%) SNPs, respectively. Even though there was no obvious complementarity among these markers due to the absence of intersections, we found GO terms with similar biological processes (Supplementary Tables 1-3), indicating common molecular processes related to these genomic regions.

### 3.4. Neural network creation

With the marker dataset established through FS for EG1 subtasks, we estimated the best hyperparameter configuration for creating the networks proposed: (i) subpopulation prediction in EG1 (Supplementary Fig. 26), (ii) BLUP prediction in EG1-Pop1 (Supplementary Fig. 27), (iii) BLUP prediction in EG1-Pop2 (Supplementary Fig. 28), and (iv) BLUP prediction in EG1-Pop3 (Supplementary Fig. 29). With the exception of network (i), which is a classification task, for each hyperparameter combination, we evaluated the MAPE and R Pearson coefficient values using the development set to select the best configuration for prediction. For network (i), several hyperparameter combinations returned prediction capabilities without mistakes (Supplementary Fig. 26), which led us to select the configuration with the minimum value for the loss function (Table 2).

**Table 2.** Hyperparameter definition for each one of the created neural networks in experimental groups 1 (EG1) and 2 (EG2) considering (i) the number of neurons selected for the first hidden layer (N-1HL), (ii) the number of neurons selected for the second hidden layer (N-2HL), (iii) the learning rate (LR), and (iv) the activation function (AF).

| Neural Network | N-1HL | N-2HL | LR | AF |
| --- | --- | --- | --- | --- |
| EG1 (Subpopulation Prediction) | 45 | 25 | 0.005 | Rectified linear activation |
| EG1 (BLUP Prediction in GT1 x PB235) | 10 | 3 | 0.005 | Rectified linear activation |
| EG1 (BLUP Prediction in GT1 x RRIM701) | 30 | 7 | 0.005 | Rectified linear activation |
| EG1 (BLUP Prediction in PR255 x PB217) | 42 | 4 | 0.005 | Rectified linear activation |

For networks (ii), (iii) and (iv), we selected the best hyperparameter combination by evaluating the plot profiles. We selected the combinations closest to the right corner of the plots (Supplementary Figs. 27-29), ideally representing the best MAPE and R Pearson coefficient simultaneously. Interestingly, for the four networks, the best activation function was ReLU, and the learning rate was 0.005, only changing the quantity of neurons in the established HLs. An evaluation of the predictive performance of these networks compared to the traditional genomic prediction approaches with k-fold CV built in the development set revealed significant improvement and effective performance in each population, different from the FS performed using these datasets combined (Supplementary Fig. 19).

The network modeled for EG1-Pop1 showed the largest increases (Supplementary Fig. 30), with a mean improvement of 9 times the initial obtained accuracies. EG1-Pop2 (Supplementary Fig. 31) and EG1-Pop3 (Supplementary Fig. 32) showed increases of 7 and 3 times, respectively. In addition to such significant improvements, the models’ performance was also more stable, with the predictive accuracies having a narrow distribution, as observed in the boxplots’ conformations.

### 3.5. Divide-and-conquer approach

All of the individual networks were combined to create the proposed approach in EG1. Compared with the traditional approaches, this approach showed a mean improvement of 4 times the initial accuracies (Fig. 2A) in the k-fold evaluations. Moreover, BRR and SM-GK presented equivalent performance values. Additionally, when analyzing the performance of the development set for predicting the BLUP values of genotypes from the test set, we found Pearson R coefficients of 0.39, 0.42, and 0.81 for BRR, SM-GK, and the proposed approach, respectively, showing the methodology’s efficiency even for data not in the development set.

**Fig. 2.**
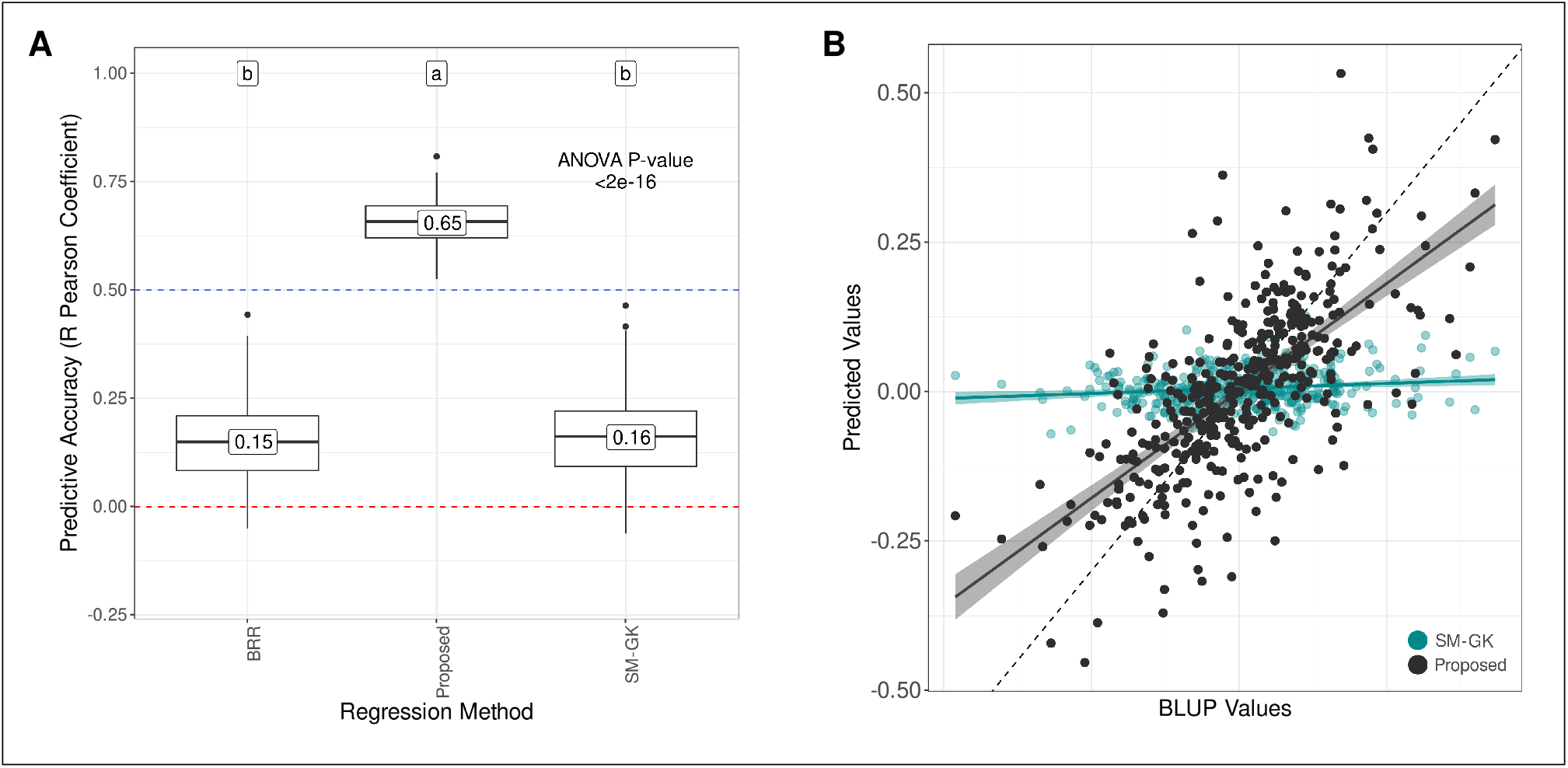
Predictive accuracies for stem circumference BLUP prediction in experimental group 1 (EG1) considering (A) a 4-fold cross validation (CV) scheme (50 times repeated) and (B) a leave-one-out CV strategy. The models used for prediction were a single-environment model with a nonlinear Gaussian kernel (SM-GK), Bayesian ridge regression (BRR), and the proposed strategy using the divide-and-conquer approach. The labels indicate the results from Tukey’s multiple comparison test.

As the final step in model evaluation, we performed a LOO CV split to check whether an increase in the training data improves prediction accuracy. By contrasting the real BLUP values with the predicted values, we found R Pearson coefficients of 0.14, 0.16 and 0.68 for BRR, SM-GK, and the proposed approach, respectively. The regression curve clearly indicates the proposed approach’s appropriateness for rubber tree data (Fig. 2B).

## 4. Discussion

GS has emerged as a potential tool for application in plant breeding programs (Cros et al., 2015; Crossa et al., 2016; O’Connor et al., 2018; Spindel et al., 2015; Wolfe et al., 2017; Xavier et al., 2016; Zhao et al., 2012). In rubber tree, previously obtained results (Cros et al., 2019; Souza et al., 2019) have demonstrated the potential of such a technique for reducing breeding cycles. Because of the strong commercial rubber demand, there have been many economic incentives for rubber tree production in more environments beyond its natural range (Ahrends et al., 2015; Warren-Thomas et al., 2015). Considering the difficulty of achieving ideal conditions for cultivating *H. brasiliensis* and the rubber demand, the development of more efficient varieties is needed. However, *Hevea*’s long life cycle considerably reduces breeding efficiency (An et al., 2019). Therefore, the application of GS in rubber tree represents an alternative for achieving the desired rubber production in less time by replacing clone trials and reducing the long period of phenotypic evaluation (Cros et al., 2019).

The main objective of rubber tree breeding programs is to increase latex production with rapid growth (Rosa et al., 2018). Increased SC development can be associated with several rubber tree characteristics, such as growth (Chandrashekar et al., 1998), latex production (Souza et al., 2019), and drought resistance (Zhang et al., 2019). Due to the high versatility of SC in evaluating rubber trees (Chanroj et al., 2017; Dijkman et al., 1951; Gonçalves et al., 1984; Khan et al., 2018), we proposed to develop more effective models for predicting this trait, providing a method to be incorporated into the estimation of tree performance. The lack of high genotype variability in the datasets used represents a real scenario for rubber tree breeding programs (Souza et al., 2019), which face the difficulty of generating a population (Cros et al., 2019). In addition to the within-family approach suggested for GS with full-sib families by Cros et al. (2019), the use of interconnected families is a common strategy for perennial species (Grattapaglia, 2017; Kumar et al., 2015; Muranty et al., 2015).

Using these dataset configurations, we evaluated ML algorithms as a more accurate methodology for predicting SC, a complex trait. Cros et al. (2019) obtained a mean accuracy for rubber production in a CV0 scenario of 0.53, which increased to 0.56 when selecting a set of markers based on heterozygosity values. In a CV1 scheme, the mean values ranged between 0.33 and 0.60. In the proposed work, we observed even lower accuracies when using SC instead of rubber production, which is in accordance with the findings of Souza et al. (2019). In (Souza et al., 2019), the authors achieved mean accuracies ranging between 0.19 and 0.28 in a CV1 scenario, contrasted with a CV2 scheme with values ranging between 0.84 and 0.86. For unknown tested genotypes, the predictive accuracies in rubber tree are low, and the inclusion of GS in *Hevea* breeding programs is therefore still not feasible.

Using the traditional approaches for prediction, we achieved LOO configurations of 0.14 and 0.16 for the BRR and SM-GK approaches, respectively, which is similar to what Souza et al. (2019) observed. The BRR and SM-GK methodologies were selected to represent a parametric and a semiparametric approach (Heslot et al., 2012). Different from BRR, which estimates marker effects, SM-GK estimates genotype effects through a relationship matrix obtained with a reproducing kernel (Granato et al., 2018a). Even though Souza et al. (2019) found similar results when using a linear and a nonlinear kernel for the estimation of the genomic relationship matrix, Gianola et al. (2014) considered GK to have a more flexible structure and a higher associated performance. Therefore, considering these findings together with the fact that no significant differences have been found among statistical models for GS (Ma et al., 2018; Roorkiwal et al., 2016; Varshney, 2016), we selected only these two statistical models for predictive evaluation.

Even though some previous attempts did not reveal significant differences in employing ML in GS compared with traditional linear regression methodologies (Crossa et al., 2019; Montesinos-López et al., 2019a, 2018, 2019b; Zingaretti et al., 2020), this is not what we observed in our study, which corroborates the findings of Bellot et al. (2018); Liu et al. (2019); Ma et al. (2018); Waldmann et al. (2020). This discrepancy may be explained by the different strategies used in the ML algorithms, especially distinct neural network architectures, training methodologies, and CV scenarios. The design of neural network architectures is an important step in using deep learning for prediction because differences in the definition of topologies can lead to decreased accuracies (Ma et al., 2018).

### 4.1. Divide-and-conquer strategy

Several factors are known to influence prediction accuracy in GS, such as the relationship between the individuals used to train models and those that will be predicted (Washburn et al., 2019), the size and structure of the populations used (Crossa et al., 2017), the trait heritability (Zhang et al., 2017), the marker density (Liu et al., 2018), and the linkage disequilibrium (LD) between the set of markers used and the associated QTLs (Raymond et al., 2018). This last aspect is especially critical in the datasets employed because of the limited set of markers obtained through GBS and SSR genotyping. Considering the reduced accuracies obtained with the CV1 technique already described in (Cros et al., 2019; Souza et al., 2019), it was expected that when using a K-fold strategy, the same observations would be found for the traditional regression models.

One of the main challenges in GS is the high dimensionality of the features in the datasets because the number of SNPs is much larger than the number of phenotypic observations (Long et al., 2007) (‘large *p*, small *n*’ problem). Although a greater saturation of markers enables an increase in the probability of finding LD, a larger number of markers in the same LD block does not contribute to better prediction performance (Liu et al., 2018). In this context, FS techniques may be an alternative strategy for building a predictive model, considering that not all markers are related to a specific phenotype (Yin et al., 2019) and that the quantity required for this task directly depends on the complexity and genetic architecture of the traits used (Liu et al., 2018). Therefore, like Bermingham et al. (2015), Bellot et al. (2018), Li et al. (2018), Inácio & Alves (2019), Aono et al. (2020), Ramzan et al. (2020), Luo et al. (2021), and Pimenta et al. (2021), we decided to test the prediction improvements by using an FS technique to enhance network performances.

Subset selection showed improvements for EG2 (Supplementary Figs. 20-21); however, there were no sizable improvements because of the genetic complexity of SC (Francisco et al., 2021) and the low density of SSR markers (Nadeem et al., 2018). In EG1, although an overall improvement in prediction accuracy was observed (Supplementary Fig. 19), when evaluating the intrapopulation predictive accuracy, we observed clear inefficiency of the approach, probably caused by the different allele substitution effects between the three subpopulations employed (Raymond et al., 2018). In such a scenario with unbalanced interconnected families, novel approaches are needed, and in this work, we have proposed the use of a divide-and-conquer strategy.

In computer science, the divide-and-conquer paradigm is based on the principle that if a problem is not simple enough to be solved directly, it can be divided into subproblems, and their results can be combined (Smith, 1985). In our prediction task, the BLUPs of the populations could not be properly predicted together; thus, we separated the problem into different networks for prediction, combining the strategy into a single network structure. Such an approach has already been applied to the development of neural network architectures (Feng et al., 2019; Frosyniotis et al., 2003; Mohamad, 2013; Sakhakarmi & Park, 2020); however, such a formulation has not been explored in genomic prediction. In addition to increasing prediction accuracies, such an approach can reduce the time required for network training and hyperparameter estimation (Mohamad, 2013), supply superior model interpretability without loss of performance (Fu et al., 2019), and be used in combination with other models (Intanagonwiwat, 1998), including traditional genomic prediction methods. Considering that in genomic prediction, most of the scenarios include different population structures, such a paradigm can benefit the application and development of GS strategies.

In our dataset, most of the observed variance within SNP markers was caused by population structure, which is clearly shown by the PCA results (Supplementary Fig. 14). As this strong variability can be associated with several genomic regions and influence various traits differently and simultaneously in the populations (Linhart & Grant, 1996), we hypothesize that traditional genomic prediction models are not capable of capturing these interpopulation differences related to SC QTLs. This is the main reason why performing FS on these unbalanced datasets together was not a promising strategy in our study. As intrapopulation QTLs are not transferable to other populations, the main effects on phenotypic variation are specific to the within-population genetic structure (Würschum, 2012). In this sense, the prediction task in single populations can be seen as simpler than that in multiple populations (Ogut et al., 2015), which was the basis for developing the divide-and-conquer strategy. Considering the specific effects of causal genetic variants within populations (Hirschhorn et al., 2001; Pressoir & Berthaud, 2004), we tried to incorporate such factors into separate networks with their specific hyperparameter optimization processes.

Interestingly, FS steps performed in the three different populations of EG1 returned different markers, but these markers were putatively associated with genes acting in similar biological processes. GO mRNA splicing was found in the intersection set of markers selected for the three populations. The occurrence of genetic variation related to such a regulatory process may influence the transcription of diverse mRNAs from the same gene in different ways. Such diversity of molecules may be related to differences in phenotypic performance, leading to increased plant capabilities (Mastrangelo et al., 2012; Szakonyi & Duque, 2018; Wei et al., 2017). Additionally, base-excision repair was found in both Pop1 and Pop3, which represents a very important defense pathway for maintaining genomic integrity (Roldán-Arjona et al., 2019) and is clearly essential for rubber tree growth and development (Murphy, 2005). Due to the increased quantity of individuals in Pop2 and Pop3, more GO categories were found, including important processes for plant growth, such as response to different types of stress and several metabolic processes (Francisco et al., 2021).

### 4.2. Deep learning architectures

Different studies have reported the use of deep learning for genomic prediction with various datasets, including for humans (Bellot et al., 2018; Yin et al., 2019), sows (Waldmann et al., 2020), and plant species such as soybean (Liu et al., 2019), wheat (Crossa et al., 2019; Ma et al., 2018; Montesinos-López et al., 2019a, 2018, 2019b), maize (Montesinos-López et al., 2018), and strawberry and blueberry (Zingaretti et al., 2020). Even though all of these studies used deep learning, the neural network creation approaches were not the same; some of them included architectures of convolutional neural networks (CNNs) (Waldmann et al., 2020; Yin et al., 2019; Zingaretti et al., 2020), while others included MLPs (Crossa et al., 2019; Montesinos-López et al., 2019a, 2018, 2019b) or both approaches (Bellot et al., 2018; Liu et al., 2019; Ma et al., 2018). There is no consensus on the efficiency of neural networks for genomic prediction; however, we decided to use such an architecture for combining multiple training processes into a single predictive structure.

For each of the neural network architectures, we employed an MLP structure. We did not include convolutional operations because of the reduced quantity of markers obtained through FS. Additionally, CNNs were developed for extracting unknown patterns from the dataset, and as we hypothesized that FS operations might work as indicators of QTL regions, such operations would not be necessary. To define the most promising network architecture, we used a grid search, testing different combinations of hyperparameters as already performed in relation to GS strategies (Crossa et al., 2019; Montesinos-López et al., 2019a, 2018, 2019b). Although other researchers have used the ‘trial and error’ approach to define the network topology (Sheela & Deepa, 2013), we preferred to develop a strategy that could be replicated in other predictive scenarios, especially with other traits and crops.

The approximation of functions through neural networks was supported first based on Kolmogorov (1957) and later on Hecht-Nielsen (1987), which extended the theorem of Kolmogorov (1957), proving that any continuous function can be represented by a neural network with one HL containing 2*n* + 1 nodes (*n* features) and a more complex activation function than that usually employed by current researchers (Stathakis, 2009). It has already been proven that one HL is capable of universal approximation by using a complex activation function (Hornik, 1993; Hornik et al., 1989; Huang, 2003; Thomas et al., 2017; Wang, 2003); however, when using regular functions, such as sigmoid and ReLU functions, there is reduced efficiency of such networks. In this context, Kurková (1992) suggested that two HLs could be a solution for this reduced efficiency. In addition, the usage of an additional HL can substantially reduce the total number of required nodes for a satisfactory predictive capability (Stathakis, 2009), and it has already been shown that some problems can be solved only by the use of two HLs (Chester, 1990; Sontag, 1991; Thomas et al., 2017). In practical situations, a neural network architecture with two HLs generalizes better than that with one and has been considered a superior approach (Islam & Murase, 2001; Thomas et al., 2017). Therefore, in our study, we decided to include two HLs in our proposed architecture, representing a network with more complex training complexity (Kurková & Sanguineti, 2013).

Concerning the quantity of hidden neurons in a neural network, many researchers have developed different strategies, aiming at increasing accuracy and prediction while decreasing errors (Sheela & Deepa, 2013). Huang (2003) has already proven that in a network architecture with two HLs, the number of nodes required to achieve a reasonable predictive accuracy with *m* samples and *q* output neurons is 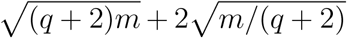 in the first HL and 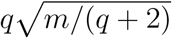 in the second HL. However, the quantity of suggested nodes tends to lead to overfitting of the training data with any arbitrary small error (Sheela & Deepa, 2013), and considering the capability of predicting unknown data, these values can be considered the maximum number of nodes in an artificial neural network structure (Stathakis, 2009). The lower bound for hidden neurons was already proposed by Jiang et al. (2008), which can be useful for accelerating the learning speed, but there was no evidence on separating this quantity across HLs, and the study was based on an MLP with 3 HLs (Sheela & Deepa, 2013). Thus, in our architecture definition, we decided to test a large quantity of neurons, considering the findings of Huang (2003), as our upper bound.

The created network coupling the population-specific architectures could increase the initial prediction capabilities by more than four times. Such an improvement represents the first attempt to develop a ML strategy for genomic prediction in rubber tree, with a high potential to be adapted to other species with the same data configuration. Considering a broader scenario with distantly related genotypes belonging to a population with undefined structure, this same approach could be applied. Instead of relying on the predefined stratification, clustering analyses could be performed and used for the divide part. Such a practice is already common in breeding, i.e., taking advantage of population structure for model prediction through multivariate techniques (Berro et al., 2019; Guo et al., 2014; Stewart-Brown et al., 2019; Wang et al., 2017). Taking into account the importance of such group configuration in the differentiation of multiple traits (Bolnick et al., 2011; Goodnight, 1989; Merilä & Crnokrak, 2001), the strategy developed represents a promising approach for several plant species with a difficult prediction scenario.

The use of GS in rubber tree can optimize breeding programs, and the incorporation of ML techniques can be seen as a new possibility for building more robust models with higher associated prediction capabilities. By using data from rubber tree breeding programs, we were able to generate promising predictive results for a highly complex trait and a novel strategy for prediction, which has significant potential to enhance selection efficiency, reduce the length of the selection cycle, and supply a means of developing low-density markers to be employed in MAS because of the FS steps. Although our results confirmed the efficiency of the methodology proposed for rubber tree data, to properly evaluate the full potential of the method in other species and broader scenarios, our approach should be investigated in further studies with more genetically diverse populations in contrasting environments.

## Author contributions

AA and FF performed all the analyses and wrote the manuscript; PG, EJ and VG conducted the field experiments; LS, RF, MQ, GG and AS conceived the project. All authors reviewed, read and approved the manuscript.

## Acknowledgements

The authors gratefully acknowledge the Fundação de Amparo à Pesquisa do Estado de São Paulo (FAPESP) for Ph.D. fellowships to FF (2018/18985-7) and AA (2019/03232-6) and for a research internship abroad (BEPE) scholarship to AA (2019/26858-8), the Coordenação de Aperfeiçoamento do Pessoal de Nível Superior (CAPES) for financial support (Computational Biology Program and CAPES-Agropolis Program), and the Conselho Nacional de Desenvolvimento Científico e Tecnológico (CNPq) for research fellowships to AS and PG.

## Data availability statement

The genotypic data from EG1 are available under NCBI accessions PRJNA540286 (ID: 5440286) (GT1 × PB235 and GT1 × RRIM701) and PRJNA541308 (ID: 541308) (PR255 × PB217). The datasets from EG2 were made available by Cros et al. (2019).

## Notes

### Competing Interest Statement

The authors have declared no competing interest.

## References

Ahrends, A., Hollingsworth, P. M., Ziegler, A. D., Fox, J. M., Chen, H., Su, Y., & Xu, J., 2015. Current trends of rubber plantation expansion may threaten biodiversity and livelihoods. Global Environ. Change, 34, 48–58.

Albrecht, T., Wimmer, V., Auinger, H.-J., Erbe, M., Knaak, C., Ouzunova, M., Simianer, H., & Schön, C.-C. (2011. Genome-based prediction of testcross values in maize. Theor. Appl. Genet., 123, 339.

Altschul, S. F., Gish, W., Miller, W., Myers, E. W., & Lipman, D. J., 1990. Basic local alignment search tool. J. Mol. Biol., 215, 403–410.

An, Z., Zhao, Y., Zhang, X., Huang, X., Hu, Y., Cheng, H., Li, X., & Huang, H., 2019. A high-density genetic map and qtl mapping on growth and latex yield-related traits in hevea brasiliensis müll. arg. Ind. Crops Prod., 132, 440–448.

Aono, A. H., Costa, E. A., Rody, H. V. S., Nagai, J. S., Pimenta, R. J. G., Mancini, M. C., Dos Santos, F. R. C., Pinto, L. R., de Andrade Landell, M. G., de Souza, A. P. et al., 2020. Machine learning approaches reveal genomic regions associated with sugarcane brown rust resistance. Sci. Rep., 10, 1–16.

Baudouin, L., Baril, C., Clément-Demange, A., Leroy, T., & Paulin, D., 1997. Recurrent selection of tropical tree crops. Euphytica, 96, 101–114.

Bellot, P., de los Campos, G., & Pérez-Enciso, M., 2018. Can deep learning improve genomic prediction of complex human traits? Genetics, 210, 809–819.

Bengio, Y., 2015. Rmsprop and equilibrated adaptive learning rates for nonconvex optimization. corr abs/1502.04390.

Bermingham, M. L., Pong-Wong, R., Spiliopoulou, A., Hayward, C., Rudan, I., Campbell, H., Wright, A. F., Wilson, J. F., Agakov, F., Navarro, P. et al., 2015. Application of high-dimensional feature selection: evaluation for genomic prediction in man. Sci. Rep., 5, 1–12.

Bernardo, R., & Yu, J., 2007. Prospects for genomewide selection for quantitative traits in maize. Crop Sci., 47, 1082–1090.

Berro, I., Lado, B., Nalin, R. S., Quincke, M., & Gutiérrez, L., 2019. Training population optimization for genomic selection. Plant Genome, 12, 190028.

Bolnick, D. I., Amarasekare, P., Araújo, M. S., Bürger, R., Levine, J. M., Novak, M., Rudolf, V. H., Schreiber, S. J., Urban, M. C., & Vasseur, D. A., 2011. Why intraspecific trait variation matters in community ecology. Trends Ecol. Evol., 26, 183–192.

Botstein, D., Cherry, J. M., Ashburner, M., Ball, C. A., Blake, J. A., Butler, H., Davis, A. P., Dolinski, K., Dwight, S. S., Eppig, J. T. et al., 2000. Gene ontology: tool for the unification of biology. Nat. Genet., 25, 25–9.

Breiman, L., 2001. Random forests. Mach. Learn., 45, 5–32.

Browning, S. R., & Browning, B. L., 2007. Rapid and accurate haplotype phasing and missing-data inference for whole-genome association studies by use of localized haplotype clustering. Am. J. Hum. Genet., 81, 1084–1097.

Chandrashekar, T., Nazeer, M., Marattukalam, J., Prakash, G., Annamalainathan, K., & Thomas, J., 1998. An analysis of growth and drought tolerance in rubber during the immature phase in a dry subhumid climate. Exp. Agric., 34, 287–300.

Chanroj, V., Rattanawong, R., Phumichai, T., Tangphatsornruang, S., & Ukoskit, K., 2017. Genome-wide association mapping of latex yield and girth in amazonian accessions of hevea brasiliensis grown in a suboptimal climate zone. Genomics, 109, 475–484.

Charrad, M., Ghazzali, N., Boiteau, V., & Niknafs, A., 2014. Nbclust: an r package for determining the relevant number of clusters in a data set. J. Stat. Softw., 61, 1–36.

Chen, T., & Guestrin, C., 2016. Xgboost: A scalable tree boosting system. In Proceedings of the 22nd ACM sigkdd international conference on knowledge discovery and data mining (pp. 785–794).

Chester, D. L., 1990. Why two hidden layers are better than one. In Proc. IJCNN, Washington, DC (pp. 265–268). volume 1.

Chollet, F. et al., 2015. Keras. https://keras.io.

Conson, A. R., Taniguti, C. H., Amadeu, R. R., Andreotti, I. A., de Souza, L. M., dos Santos, L. H., Rosa, J. R., Mantello, C. C., da Silva, C. C., José Scaloppi Junior, E. et al., 2018. High-resolution genetic map and qtl analysis of growth-related traits of hevea brasiliensis cultivated under suboptimal temperature and humidity conditions. Front. Plant Sci., 9, 1255.

Cover, T., & Hart, P., 1967. Nearest neighbor pattern classification. IEEE Trans. Inf. Theory, 13, 21–27.

Cros, D., Denis, M., Sánchez, L., Cochard, B., Flori, A., Durand-Gasselin, T., Nouy, B., Omoré, A., Pomiès, V., Riou, V. et al., 2015. Genomic selection prediction accuracy in a perennial crop: case study of oil palm (elaeis guineensis jacq.). Theor. Appl. Genet., 128, 397–410.

Cros, D., Mbo-Nkoulou, L., Bell, J. M., Oum, J., Masson, A., Soumahoro, M., Tran, D. M., Achour, Z., Le Guen, V., & Clement-Demange, A., 2019. Within-family genomic selection in rubber tree (hevea brasiliensis) increases genetic gain for rubber production. Ind. Crops Prod., 138, 111464.

Crossa, J., Jarquín, D., Franco, J., Pérez-Rodríguez, P., Burgueño, J., Saint-Pierre, C., Vikram, P., Sansaloni, C., Petroli, C., Akdemir, D. et al., 2016. Genomic prediction of gene bank wheat landraces. G3: Genes Genom. Genet., 6, 1819–1834.

Crossa, J., Martini, J. W., Gianola, D., Pérez-Rodríguez, P., Jarquin, D., Juliana, P., Montesinos-López, O., & Cuevas, J., 2019. Deep kernel and deep learning for genome-based prediction of single traits in multienvironment breeding trials. Front. Genet., 10.

Crossa, J., Perez, P., Hickey, J., Burgueno, J., Ornella, L., Cerón-Rojas, J., Zhang, X., Dreisigacker, S., Babu, R., Li, Y. et al., 2014. Genomic prediction in CIMMYT maize and wheat breeding programs. Heredity, 112, 48–60.

Crossa, J., Pérez-Rodríguez, P., Cuevas, J., Montesinos-López, O., Jarquín, D., de los Campos, G., Burgueño, J., González-Camacho, J. M., Pérez-Elizalde, S., Beyene, Y. et al., 2017. Genomic selection in plant breeding: methods, models, and perspectives. Trends Plant Sci., 22, 961–975.

Cuevas, J., Crossa, J., Soberanis, V., Pérez-Elizalde, S., Pérez-Rodríguez, P., Campos, G. d. l., Montesinos-López, O., & Burgueño, J., 2016. Genomic prediction of genotype× environment interaction kernel regression models. Plant Genome, 9.

Da Silva, I. N., Spatti, D. H., Flauzino, R. A., Liboni, L. H. B., & dos Reis Alves, S. F., 2017. Artificial Neural Networks. Cham: Springer International Publishing, (p. 39).

Danecek, P., Auton, A., Abecasis, G., Albers, C. A., Banks, E., DePristo, M. A., Handsaker, R. E., Lunter, G., Marth, G. T., Sherry, S. T. et al., 2011. The variant call format and VCFtools. Bioinformatics, 27, 2156–2158.

De Los Campos, G., Naya, H., Gianola, D., Crossa, J., Legarra, A., Manfredi, E., Weigel, K., & Cotes, J. M., 2009. Predicting quantitative traits with regression models for dense molecular markers and pedigree. Genetics, 182, 375–385.

Dijkman, M. J. et al., 1951. Hevea, thirty years of research in the far east. Hevea, Thirty years of research in the Far East..

Elshire, R. J., Glaubitz, J. C., Sun, Q., Poland, J. A., Kawamoto, K., Buckler, E. S., & Mitchell, S. E., 2011. A robust, simple genotyping-by-sequencing (GBS) approach for high diversity species. PLoS One, 6.

Endelman, J. B., 2011. Ridge regression and other kernels for genomic selection with R package rrBLUP. Plant Genome, 4, 250–255.

Feng, J., Wang, L., Yu, H., Jiao, L., & Zhang, X., 2019. Divide-and-conquer dual-architecture convolutional neural network for classification of hyperspectral images. Remote Sens., 11, 484.

Francisco, F. R., Aono, A. H., da Silva, C. C., Gonçalves, P. d. S., Scaloppi Junior, E. J., Le Guen, V., Neto, R. F., Souza, L. M. D., & de Souza, A. P., 2021. Unravelling rubber tree growth by integrating GWAS and biological network-based approaches. Front. Plant Sci., (p. 2719).

Freund, Y., & Schapire, R. E., 1997. A decision-theoretic generalization of on-line learning and an application to boosting. J. Comput. Syst. Sci., 55, 119–139.

Frosyniotis, D., Stafylopatis, A., & Likas, A., 2003. A divide-and-conquer method for multi-net classifiers. Pattern Anal. Appl., 6, 32–40.

Fu, W., Breininger, K., Schaffert, R., Ravikumar, N., & Maier, A., 2019. A divide-and-conquer approach towards understanding deep networks. In International Conference on Medical Image Computing and Computer-Assisted Intervention (pp. 183–191). Springer.

Gianola, D., 2013. Priors in whole-genome regression: The bayesian alphabet returns. Genetics, 194, 573–596.

Gianola, D., Weigel, K. A., Krämer, N., Stella, A., & Schön, C.-C., 2014. Enhancing genome-enabled prediction by bagging genomic blup. PLoS One, 9.

Glaubitz, J. C., Casstevens, T. M., Lu, F., Harriman, J., Elshire, R. J., Sun, Q., & Buckler, E. S., 2014. Tassel-GBS: A high capacity genotyping by sequencing analysis pipeline. PLoS One, 9, e90346.

Gonçalves, P. d. S., Rossetti, A. G., Valois, A. C. C., & Viegas, I., 1984. Estimativas de correlações genéticas e fenotípicas de alguns caracteres quantitativos em clones jovens de seringueira (hevea spp). Embrapa Amazônia Ocidental-Artigo em periódico indexado (AL-ICE).

González-Camacho, J., de Los Campos, G., Pérez, P., Gianola, D., Cairns, J., Mahuku, G., Babu, R., & Crossa, J., 2012. Genome-enabled prediction of genetic values using radial basis function neural networks. Theor. Appl. Genet., 125, 759–771.

González-Camacho, J. M., Ornella, L., Pérez-Rodríguez, P., Gianola, D., Dreisigacker, S., & Crossa, J. 2018. Applications of machine learning methods to genomic selection in breeding wheat for rust resistance. Plant Genome, 11.

Goodnight, C. J., 1989. Population differentiation and the correlation among traits at the population level. Am. Nat., 133, 888–900.

Goodstein, D. M., Shu, S., Howson, R., Neupane, R., Hayes, R. D., Fazo, J., Mitros, T., Dirks, W., Hellsten, U., Putnam, N. et al., 2012. Phytozome: a comparative platform for green plant genomics. Nucleic Acids Res., 40, D1178–D1186.

Granato, I., Cuevas, J., Luna-Vázquez, F., Crossa, J., Montesinos-López, O., Burgueño, J., & Fritsche-Neto, R., 2018a. BGGE: a new package for genomic-enabled prediction incorporating genotype× environment interaction models. G3: Genes Genom. Genet., 8, 3039–3047.

Granato, I. S., Galli, G., de Oliveira Couto, E. G., e Souza, M. B., Mendonça, L. F., & Fritsche-Neto, R., 2018b. snpready: a tool to assist breeders in genomic analysis. Mol. Breed., 38, 102.

Grattapaglia, D., 2017. Status and perspectives of genomic selection in forest tree breeding. In Genomic Selection for Crop Improvement (pp. 199–249). Springer, New York.

Guo, Z., Tucker, D. M., Basten, C. J., Gandhi, H., Ersoz, E., Guo, B., Xu, Z., Wang, D., & Gay, G., 2014. The impact of population structure on genomic prediction in stratified populations. Theor. Appl. Genet., 127, 749–762.

Harfouche, A. L., Jacobson, D. A., Kainer, D., Romero, J. C., Harfouche, A. H., Mugnozza, G. S., Moshelion, M., Tuskan, G. A., Keurentjes, J. J., & Altman, A., 2019. Accelerating climate resilient plant breeding by applying next-generation artificial intelligence. Trends Biotechnol..

Hayes, B., Goddard, M. et al., 2001. Prediction of total genetic value using genome-wide dense marker maps. Genetics, 157, 1819–1829.

Hayes, B. J., Lewin, H. A., & Goddard, M. E., 2013. The future of livestock breeding: genomic selection for efficiency, reduced emissions intensity, and adaptation. Trends Genet., 29, 206–214.

Hecht-Nielsen, R., 1987. Kolmogorov’s mapping neural network existence theorem. In Proceedings of the International Conference on Neural Networks (pp. 11–14). IEEE Press, New York volume 3.

Hecht-Nielsen, R., 1992. Theory of the backpropagation neural network. In Neural Networks for Perception (pp. 65–93). Elsevier.

Heffner, E. L., Lorenz, A. J., Jannink, J.-L., & Sorrells, M. E., 2010. Plant breeding with genomic selection: gain per unit time and cost. Crop Sci., 50, 1681–1690.

Heslot, N., Yang, H.-P., Sorrells, M. E., & Jannink, J.-L., 2012. Genomic selection in plant breeding: a comparison of models. Crop Sci., 52, 146–160.

Hirschhorn, J. N., Lindgren, C. M., Daly, M. J., Kirby, A., Schaffner, S. F., Burtt, N. P., Altshuler, D., Parker, A., Rioux, J. D., Platko, J. et al., 2001. Genomewide linkage analysis of stature in multiple populations reveals several regions with evidence of linkage to adult height. Am. J. Hum. Genet., 69, 106–116.

Hoffer, E., Ben-Nun, T., Hubara, I., Giladi, N., Hoefler, T., & Soudry, D., 2019. Augment your batch: better training with larger batches. arXiv preprint arXiv:1901.09335.

Hoffer, E., Hubara, I., & Soudry, D., 2017. Train longer, generalize better: closing the generalization gap in large batch training of neural networks. In Advances in Neural Information Processing Systems (pp. 1731–1741).

Hornik, K., 1993. Some new results on neural network approximation. Neural Netw., 6, 1069–1072.

Hornik, K., Stinchcombe, M., White, H. et al., 1989. Multilayer feedforward networks are universal approximators. Neural Netw., 2, 359–366.

Huang, G.-B., 2003. Learning capability and storage capacity of two-hidden-layer feedforward networks. IEEE Trans. Neural Netw., 14, 274–281.

Inácio, Í. S. C. G. F., & Alves, M. F. C., 2019. Increasing accuracy and reducing costs of genomic prediction by marker selection. Euphytica, 215, 18.

Intanagonwiwat, C., 1998. The divide-and-conquer neural network: its architecture and training. In 1998 IEEE International Joint Conference on Neural Networks Proceedings. IEEE World Congress on Computational Intelligence (Cat. No. 98CH36227) (pp. 462–467). IEEE volume 1.

Islam, M. M., & Murase, K., 2001. A new algorithm to design compact two-hidden-layer artificial neural networks. Neural Netw., 14, 1265–1278.

Jannink, J.-L., Lorenz, A. J., & Iwata, H., 2010. Genomic selection in plant breeding: from theory to practice. Brief. Funct. Genom., 9, 166–177.

Jarquín, D., Lemes da Silva, C., Gaynor, R. C., Poland, J., Fritz, A., Howard, R., Batten-field, S., & Crossa, J., 2017. Increasing genomic-enabled prediction accuracy by modeling genotype× environment interactions in kansas wheat. Plant Genome, 10.

Jiang, N., Zhang, Z., Ma, X., & Wang, J., 2008. The lower bound on the number of hidden neurons in multi-valued multi-threshold neural networks. In 2008 Second International Symposium on Intelligent Information Technology Application (pp. 103–107). IEEE volume 1.

Khan, M. A., Tong, F., Wang, W., He, J., Zhao, T., & Gai, J., 2018. Analysis of QTL– allele system conferring drought tolerance at seedling stage in a nested association mapping population of soybean [G lycine max (L.) Merr.] using a novel GWAS procedure. Planta, 248, 947–962.

Kingma, D. P., & Ba, J., 2014. Adam: A method for stochastic optimization. arXiv preprint arXiv:1412.6980.

Kolmogorov, A. N., 1957. On the representation of continuous functions of many variables by superposition of continuous functions of one variable and addition. In Doklady Akademii Nauk (pp. 953–956). Russian Academy of Sciences volume 114.

Kumar, S., Molloy, C., Muñoz, P., Daetwyler, H., Chagné, D., & Volz, R., 2015. Genome-enabled estimates of additive and nonadditive genetic variances and prediction of apple phenotypes across environments. G3: Genes Genom. Genet., 5, 2711–2718.

Kurková, V., 1992. Kolmogorov’s theorem and multilayer neural networks. Neural Netw., 5, 501–506.

Kurková, V., & Sanguineti, M., 2013. Can two hidden layers make a difference? In International Conference on Adaptive and Natural Computing Algorithms (pp. 30–39). Springer, New York.

Lau, N.-S., Makita, Y., Kawashima, M., Taylor, T. D., Kondo, S., Othman, A. S., Shu-Chien, C., & Matsui, M., 2016. The rubber tree genome shows expansion of gene family associated with rubber biosynthesis. Sci. Rep., 6, 28594.

Le Guen, V., Doaré, F., Weber, C., & Seguin, M., 2009. Genetic structure of amazonian populations of hevea brasiliensis is shaped by hydrographical network and isolation by distance. Tree Genet. Genom., 5, 673–683.

Le Guen, V., Garcia, D., Doaré, F., Mattos, C. R., Condina, V., Couturier, C., Chambon, A., Weber, C., Espéout, S., & Seguin, M., 2011. A rubber tree’s durable resistance to microcyclus ulei is conferred by a qualitative gene and a major quantitative resistance factor. Tree Genet. Genom., 7, 877–889.

Le Guen, V., Garcia, D., Mattos, C. R. R., Doaré, F., Lespinasse, D., & Seguin, M., 2007. Bypassing of a polygenic microcyclus ulei resistance in rubber tree, analyzed by qtl detection. New Phytolog., 173, 335–345.

Lespinasse, D., Grivet, L., Troispoux, V., Rodier-Goud, M., Pinard, F., & Seguin, M., 2000a. Identification of QTLs involved in the resistance to South American leaf blight (Microcyclus ulei) in the rubber tree. Theor. Appl. Genet., 100, 975–984.

Lespinasse, D., Rodier-Goud, M., Grivet, L., Leconte, A., Legnaté, H., & Seguin, M., 2000b. A saturated genetic linkage map of rubber tree (Hevea spp.) based on RFLP, AFLP, microsatellite, and isozyme markers. Theor. Appl. Genet., 100, 127–138.

Li, B., Zhang, N., Wang, Y.-G., George, A. W., Reverter, A., & Li, Y., 2018. Genomic prediction of breeding values using a subset of SNPs identified by three machine learning methods. Front. Genet., 9, 237.

Li, H., & Durbin, R., 2009. Fast and accurate short read alignment with burrows–wheeler transform. Bioinformatics, 25, 1754–1760.

Linhart, Y. B., & Grant, M. C., 1996. Evolutionary significance of local genetic differentiation in plants. Ann. Rev. Ecol. Syst., 27, 237–277.

Liu, J., Shi, C., Shi, C.-C., Li, W., Zhang, Q.-J., Zhang, Y., Li, K., Lu, H.-F., Shi, C., Zhu, S.-T. et al., 2020. The chromosome-based rubber tree genome provides new insights into spurge genome evolution and rubber biosynthesis. Mol. Plant, 13, 336–350.

Liu, X., Wang, H., Wang, H., Guo, Z., Xu, X., Liu, J., Wang, S., Li, W.-X., Zou, C., Prasanna, M. et al., 2018. Factors affecting genomic selection revealed by empirical evidence in maize. Crop J., 6, 341–352.

Liu, Y., Wang, D., He, F., Wang, J., Joshi, T., & Xu, D., 2019. Phenotype prediction and genome-wide association study using deep convolutional neural network of soybean. Front. Genet., 10, 1091.

Long, N., Gianola, D., Rosa, G. J., Weigel, K. A., & Avendaño, S., 2007. Machine learning classification procedure for selecting SNPs in genomic selection: application to early mortality in broilers. J. Anim. Breed. Genet., 124, 377–389.

Luo, Z., Yu, Y., Xiang, J., & Li, F., 2021. Genomic selection using a subset of SNPs identified by genome-wide association analysis for disease resistance traits in aquaculture species. Aquaculture, 539, 736620.

Ma, W., Qiu, Z., Song, J., Li, J., Cheng, Q., Zhai, J., & Ma, C., 2018. A deep convolutional neural network approach for predicting phenotypes from genotypes. Planta, 248, 1307–1318.

Mastrangelo, A. M., Marone, D., Laid`o, G., De Leonardis, A. M., & De Vita, P., 2012. Alternative splicing: enhancing ability to cope with stress via transcriptome plasticity. Plant Sci., 185, 40–49.

de Mendiburu, F., & de Mendiburu, M. F., 2019. Package ‘agricolae’. R Package Version, (pp. 1–2).

Merilá, J., & Crnokrak, P., 2001. Comparison of genetic differentiation at marker loci and quantitative traits. J. Evol. Biol., 14, 892–903.

Mohamad, M., 2013. Divide and conquer approach in reducing ANN training time for small and large data. J. Appl. Sci., 13, 133–139.

Montesinos-Lopez, O. A., Martín-Vallejo, J., Crossa, J., Gianola, D., Hernández-Suárez, C. M., Montesinos-López, A., Juliana, P., & Singh, R., 2019a. A benchmarking between deep learning, support vector machine and bayesian threshold best linear unbiased prediction for predicting ordinal traits in plant breeding. G3: Genes Genom. Genet., 9, 601–618.

Montesinos-Lopez, O. A., Montesinos-López, A., Crossa, J., Gianola, D., Hernández-Suárez, M., & Martín-Vallejo, J., 2018. Multi-trait, multi-environment deep learning modeling for genomic-enabled prediction of plant traits. G3: Genes Genom. Genet., 8, 3829–3840.

Montesinos-Lopez, O. A., Montesinos-López, A., Tuberosa, R., Maccaferri, M., Sciara, G., Ammar, K., & Crossa, J., 2019b. Multi-trait, multi-environment genomic prediction of durum wheat with genomic best linear unbiased predictor and deep learning methods. Front. Plant Sci., 10.

Muñoz, F., & Sanchez, L., 2019. breedR: Statistical Methods for Forest Genetic Resources Analysts. URL: https://github.com/famuvie/breedR R Package Version 0.12-4.

Muranty, H., Troggio, M., Sadok, I. B., Al Rifaï, M., Auwerkerken, A., Banchi, E., Velasco, R., Stevanato, P., Van De Weg, W. E., Di Guardo, M. et al., 2015. Accuracy and responses of genomic selection on key traits in apple breeding. Hortic. Res., 2, 1–12.

Murphy, T. M., 2005. What is base excision repair good for? Knockout mutants for FPG and OGG glycosylase genes in Arabidopsis. Physiol. Plant., 123, 227–232.

Nadeem, M. A., Nawaz, M. A., Shahid, M. Q., Doğan, Y., Comertpay, G., Yıldız, M., Hatipoğlu, R., Ahmad, F., Alsaleh, A., Labhane, N. et al., 2018. Dna molecular markers in plant breeding: current status and recent advancements in genomic selection and genome editing. Biotechnol. Biotechnol. Equip., 32, 261–285.

Nakkanong, K., Nualsri, C., & Sdoodee, S., 2008. Analysis of genetic diversity in early introduced clones of rubber tree (hevea brasiliensis) using rapd and microsatellite markers. Songklanakarin J. Sci. Technol., 30.

O’Connor, K., Hayes, B., & Topp, B., 2018. Prospects for increasing yield in macadamia using component traits and genomics. Tree Genet. Genom., 14, 7.

Ogut, F., Bian, Y., Bradbury, P. J., & Holland, J. B., 2015. Joint-multiple family linkage analysis predicts within-family variation better than single-family analysis of the maize nested association mapping population. Heredity, 114, 552–563.

O’Shea, K., & Nash, R., 2015. An introduction to convolutional neural networks. arXiv preprint arXiv:1511.08458.

Pedregosa, F., Varoquaux, G., Gramfort, A., Michel, V., Thirion, B., Grisel, O., Blondel, M., Prettenhofer, P., Weiss, R., Dubourg, V. et al., 2011. Scikit-learn: Machine learning in python. J. Mach. Learn. Res., 12, 2825–2830.

Pérez, P., & de Los Campos, G., 2014. Genome-wide regression and prediction with the BGLR statistical package. Genetics, 198, 483–495.

Pérez-Rodríguez, P., Gianola, D., González-Camacho, J. M., Crossa, J., Manès, Y., & Dreisigacker, S., 2012. Comparison between linear and non-parametric regression models for genome-enabled prediction in wheat. G3: Genes Genom. Genet., 2, 1595–1605.

Peterson, R., 2017. Estimating normalization transformations with bestnormalize. URL Https-github CompetersonRbestNormalize.

Pimenta, R. J. G., Aono, A. H., Burbano, R. C. V., Coutinho, A. E., da Silva, C. C., Dos Anjos, I. A., Perecin, D., Landell, M. G. d. A., Gonçalves, M. C., Pinto, L. R. et al., 2021. Genomewide approaches for the identification of markers and genes associated with sugarcane yellow leaf virus resistance. Sci. Rep., 11, 1–18.

Popescu, M.-C., Balas, V. E., Perescu-Popescu, L., & Mastorakis, N., 2009. Multilayer perceptron and neural networks. WSEAS Transactions on Circuits and Systems, 8, 579–588.

Pressoir, G., & Berthaud, J., 2004. Patterns of population structure in maize landraces from the central valleys of Oaxaca in Mexico. Heredity, 92, 88–94.

Ramzan, F., Gültas, M., Bertram, H., Cavero, D., & Schmitt, A. O., 2020. Combining random forests and a signal detection method leads to the robust detection of genotype-phenotype associations. Genes, 11, 892.

Rao, G. P., & Kole, P., 2016. Evaluation of brazilian wild hevea germplasm for cold tolerance: genetic variability in the early mature growth. J. For. Res., 27, 755–765.

Raymond, B., Bouwman, A. C., Schrooten, C., Houwing-Duistermaat, J., & Veerkamp, R. F., 2018. Utility of whole-genome sequence data for across-breed genomic prediction. Genet. Sel. Evol., 50, 1–12.

Roldán-Arjona, T., Ariza, R. R., & Córdoba-Cañero, D., 2019. Dna base excision repair in plants: An unfolding story with familiar and novel characters. Front. Plant Sci., 10, 1055.

Romain, B., & Thierry, C., 2011. Rubberclones (hevea clonal descriptions).

Roorkiwal, M., Jarquin, D., Singh, M. K., Gaur, P. M., Bharadwaj, C., Rathore, A., Howard, R., Srinivasan, S., Jain, A., Garg, V. et al., 2018. Genomic-enabled prediction models using multi-environment trials to estimate the effect of genotype× environment interaction on prediction accuracy in chickpea. Sci. Rep., 8, 1–11.

Roorkiwal, M., Rathore, A., Das, R. R., Singh, M. K., Jain, A., Srinivasan, S., Gaur, P. M., Chellapilla, B., Tripathi, S., Li, Y. et al., 2016. Genome-enabled prediction models for yield related traits in chickpea. Front. Plant Sci., 7, 1666.

Rosa, J. R. B. F., Mantello, C. C., Garcia, D., de Souza, L. M., Da Silva, C. C., Gazaffi, R., da Silva, C. C., Toledo-Silva, G., Cubry, P., Garcia, A. A. F. et al., 2018. QTL detection for growth and latex production in a full-sib rubber tree population cultivated under suboptimal climate conditions. BMC Plant Biol., 18, 223.

Rumelhart, D. E., 1986. Learning representations by error propagation, in de rumelhart, jl mcclelland & pdp research group. Parallel Distrib. Proc., 1.

Sakhakarmi, S., & Park, J. W., 2020. Multi-level-phase deep learning using divide-and-conquer for scaffolding safety. Int. J. Environ. Res. Public Health, 17, 2391.

Shawe-Taylor, J., & Cristianini, N., 2000. An introduction to support vector machines and other kernel-based learning methods volume 204.

Sheela, K. G., & Deepa, S. N., 2013. Review on methods to fix number of hidden neurons in neural networks. Math. Probl. Eng., 2013.

Sivakumaran, S., Haridas, G., & Abraham, P., 1988. Problem of tree dryness with high yielding precocious clones and methods to exploit such clones. Proc. Coll. Hevea, 88, 253–267.

Smith, D. R., 1985. The design of divide and conquer algorithms. Sci. Comput. Program., 5, 37–58.

Sontag, E. D., 1991. Feedback stabilization using two-hidden-layer nets. In 1991 American Control Conference (pp. 815–820). IEEE.

Souza, L. M., Gazaffi, R., Mantello, C. C., Silva, C. C., Garcia, D., Le Guen, V., Cardoso, S. E. A., Garcia, A. A. F., & Souza, A. P., 2013. Qtl mapping of growth-related traits in a full-sib family of rubber tree (hevea brasiliensis) evaluated in a sub-tropical climate. PLoS One, 8.

de Souza, L. M., Toledo-Silva, G., Cardoso-Silva, C. B., Da Silva, C. C., de Araujo Andreotti, I. A., Conson, A. R. O., Mantello, C. C., Le Guen, V., & de Souza, A. P., 2016. Development of single nucleotide polymorphism markers in the large and complex rubber tree genome using next-generation sequence data. Mol. Breed., 36, 115.

Souza, L. M. d., Francisco, F. R., Gonçalves, P. d. S., Scaloppi-Junior, E. J. J., Le Guen, V., Fritsche-Neto, R., & Souza, A. P. d., 2019. Genomic selection in rubber tree breeding: A comparison of models and methods for managing g× e interactions. Front. Plant Sci., 10, 1353.

Spindel, J., Begum, H., Akdemir, D., Virk, P., Collard, B., Redona, E., Atlin, G., Jannink, J.-L., & McCouch, S. R., 2015. Genomic selection and association mapping in rice (oryza sativa): effect of trait genetic architecture, training population composition, marker number and statistical model on accuracy of rice genomic selection in elite, tropical rice breeding lines. PLoS Genet., 11, e1004982.

Stathakis, D., 2009. How many hidden layers and nodes? Int. J. Remote Sens., 30, 2133–2147.

Stewart-Brown, B. B., Song, Q., Vaughn, J. N., & Li, Z., 2019. Genomic selection for yield and seed composition traits within an applied soybean breeding program. G3: Genes Genom. Genet., 9, 2253–2265.

Szakonyi, D., & Duque, P., 2018. Alternative splicing as a regulator of early plant development. Front. Plant Sci., 9, 1174.

Tang, C., Yang, M., Fang, Y., Luo, Y., Gao, S., Xiao, X., An, Z., Zhou, B., Zhang, B., Tan, X. et al., 2016. The rubber tree genome reveals new insights into rubber production and species adaptation. Nat. Plants, 2, 1–10.

Team, R. C. et al., 2013. R: A language and environment for statistical computing.

Thomas, A. J., Petridis, M., Walters, S. D., Gheytassi, S. M., & Morgan, R. E., 2017. Two hidden layers are usually better than one. In International Conference on Engineering Applications of Neural Networks (pp. 279–290). Springer, New York.

Tran, D. M., Clément-Demange, A., Deon, M., Garcia, D., Le Guen, V., Clément-Vidal, A., Soumahoro, M., Masson, A., Label, P., Le, M. T. et al., 2016. Genetic determinism of sensitivity to corynespora cassiicola exudates in rubber tree (hevea brasiliensis). PLoS One, 11.

VanRaden, P., 2007. Genomic measures of relationship and inbreeding. INTERBULL bulletin, (pp. 33–33).

VanRaden, P. M., 2008. Efficient methods to compute genomic predictions. J. Dairy Sci., 91, 4414–4423.

Varshney, R. K., 2016. Exciting journey of 10 years from genomes to fields and markets: some success stories of genomics-assisted breeding in chickpea, pigeonpea and groundnut. Plant Sci., 242, 98–107.

Venkatachalam, P., Priya, P., Gireesh, T., Amma, C. S., & Thulaseedharan, A., 2006. Molecular cloning and sequencing of a polymorphic band from rubber tree [hevea brasiliensis (muell.) arg.]: the nucleotide sequence revealed partial homology with proline-specific permease gene sequence. Current Sci., (pp. 1510–1515).

Waldmann, P., Pfeiffer, C., & Mészáros, G., 2020. Sparse convolutional neural networks for genome-wide prediction. Front. Genet., 11.

Wang, Q., Yu, Y., Yuan, J., Zhang, X., Huang, H., Li, F., & Xiang, J., 2017. Effects of marker density and population structure on the genomic prediction accuracy for growth trait in pacific white shrimp litopenaeus vannamei. BMC Genet., 18, 1–9.

Wang, S.-C., 2003. Artificial neural network. In Interdisciplinary Computing in Java Programming (pp. 81–100). Springer, New York.

Wang, X., Xu, Y., Hu, Z., & Xu, C., 2018. Genomic selection methods for crop improvement: Current status and prospects. Crop J., 6, 330–340.

Warren-Thomas, E., Dolman, P. M., & Edwards, D. P., 2015. Increasing demand for natural rubber necessitates a robust sustainability initiative to mitigate impacts on tropical biodiversity. Conserv. Lett., 8, 230–241.

Washburn, J. D., Burch, M. B., Franco, V., & José, A., 2019. Predictive breeding for maize: Making use of molecular phenotypes, machine learning, and physiological crop models. Crop Sci..

Wei, H., Lou, Q., Xu, K., Yan, M., Xia, H., Ma, X., Yu, X., & Luo, L., 2017. Alternative splicing complexity contributes to genetic improvement of drought resistance in the rice maintainer huhan2b. Sci. Rep., 7, 1–13.

Wickham, H., 2016. ggplot2: elegant graphics for data analysis. Springer, New York.

Wolfe, M. D., Del Carpio, D. P., Alabi, O., Ezenwaka, L. C., Ikeogu, U. N., Kayondo, I. S., Lozano, R., Okeke, U. G., Ozimati, A. A., Williams, E. et al., 2017. Prospects for genomic selection in cassava breeding. Plant Genome, 10.

Würschum, T., 2012. Mapping QTL for agronomic traits in breeding populations. Theor. Appl. Genet., 125, 201–210.

Xavier, A., Muir, W. M., & Rainey, K. M., 2016. Assessing predictive properties of genome-wide selection in soybeans. G3: Genes Genom. Genet., 6, 2611–2616.

Yin, B., Balvert, M., van der Spek, R. A., Dutilh, B. E., Bohte, S., Veldink, J., & Schönhuth, A., 2019. Using the structure of genome data in the design of deep neural networks for predicting amyotrophic lateral sclerosis from genotype. Bioinformatics, 35, i538–i547.

Yu, G., Smith, D. K., Zhu, H., Guan, Y., & Lam, T. T.-Y., 2017. ggtree: an r package for visualization and annotation of phylogenetic trees with their covariates and other associated data. Methods Ecol. Evol., 8, 28–36.

Zhang, A., Wang, H., Beyene, Y., Semagn, K., Liu, Y., Cao, S., Cui, Z., Ruan, Y., Burgueño, J., San Vicente, F. et al., 2017. Effect of trait heritability, training population size and marker density on genomic prediction accuracy estimation in 22 bi-parental tropical maize populations. Front. Plant Sci., 8, 1916.

Zhang, C., Stratopoulos, L. M. F., Pretzsch, H., & Rötzer, T., 2019. How do tilia cordata greenspire trees cope with drought stress regarding their biomass allocation and ecosystem services? Forests, 10, 676.

Zhao, Y., Gowda, M., Liu, W., Würschum, T., Maurer, H. P., Longin, F. H., Ranc, N., & Reif, J. C., 2012. Accuracy of genomic selection in european maize elite breeding populations. Theor. Appl. Genet., 124, 769–776.

Zingaretti, L. M., Gezan, S. A., Ferrão, L. F. V., Osorio, L. F., Monfort, A., Muñoz, P. R., Whitaker, V. M., & Pérez-Enciso, M., 2020. Exploring deep learning for complex trait genomic prediction in polyploid outcrossing species. Front. Plant Sci., 11, 25.

