## Supplementary Material for "A divide-and-conquer approach for genomic prediction in rubber tree using machine learning"

### 1 Supplementary Tables

**Supplementary Table 1**. Gene Ontology (GO) terms summarized through the Revigo tool for the 20 SNPs selected by the feature selection techniques employed on the individuals of the GT1 x PB235 population.

| **GO Term** | **Name** | **Frequency (%)** |
|---|---|---|
| GO:0006284 | base-excision repair | 0.298165436998951 |
| GO:0000398 | mRNA splicing, via spliceosome | 0.436433865037325 |

**Supplementary Table 2**. Gene Ontology (GO) terms summarized through the Revigo tool for the 62 SNPs selected by the feature selection techniques employed on the individuals of the GT1 x RRIM701 population.

| GO Term | Name | Frequency (%) |
|---|---|---|
| GO:0006355 | regulation of transcription, DNA-templated | 10.0330757015203 |
| GO:0006468 | protein phosphorylation | 4.23350556337001 |
| GO:0006950 | response to stress | 4.70454267943585 |
| GO:0007018 | microtubule-based movement | 0.349985633268822 |
| GO:0006886 | intracellular protein transport | 1.23288694299588 |
| GO:0000103 | sulfate assimilation | 0.084209367967923 |
| GO:0000398 | mRNA splicing, via spliceosome | 0.436433865037325 |
| GO:0016192 | vesicle-mediated transport | 1.40900125690309 |
| GO:0006810 | transport | 18.3767799586826 |
| GO:0000160 | phosphorelay signal transduction system | 2.76884251732889 |
| GO:0006396 | RNA processing | 3.83107599133337 |
| GO:0006412 | translation | 5.34694375263645 |
| GO:0006351 | transcription, DNA-templated | 2.0006354738138 |

**Supplementary Table 3**. Gene Ontology (GO) terms summarized through the Revigo tool for the 119 SNPs selected by the feature selection techniques employed on the individuals of the PR255 x PB217 population.

| **GO Term** | **Name** | **Frequency (%)** |
|---|---|---|
| GO:0006468 | protein phosphorylation | 4.23350556337001 |
| GO:0008152 | metabolic process | 62.7747296551307 |
| GO:0009611 | response to wounding | 0.102041794217188 |
| GO:0010223 | secondary shoot formation | 0.002263648270697 |
| GO:0006886 | intracellular protein transport | 1.23288694299588 |
| GO:0009694 | jasmonic acid metabolic process | 0.00696443610291 |
| GO:0022900 | electron transport chain | 1.71627498667009 |
| GO:0071704 | organic substance metabolic process | 55.4341066032933 |
| GO:0005975 | carbohydrate metabolic process | 5.96269325897111 |
| GO:0000398 | mRNA splicing, via spliceosome | 0.436433865037325 |
| GO:0006788 | heme oxidation | 0.016089251851028 |
| GO:0009116 | nucleoside metabolic process | 0.779120471856656 |
| GO:0006260 | DNA replication | 1.52888373885959 |
| GO:0006465 | signal peptide processing | 0.134129421528815 |
| GO:0000160 | phosphorelay signal transduction system | 2.76884251732889 |
| GO:0007165 | signal transduction | 7.31364928685992 |
| GO:0034968 | histone lysine methylation | 0.101822864731154 |
| GO:0016571 | histone methylation | 0.115326270199963 |
| GO:0006355 | regulation of transcription, DNA-templated | 10.0330757015203 |
| GO:0009408 | response to heat | 0.187738230392138 |

| | | |
|---|---|---|
| GO:0043039 | tRNA aminoacylation | 1.06491019677713 |
| GO:0006419 | alanyl-tRNA aminoacylation | 0.058288942970439 |
| GO:0006633 | fatty acid biosynthetic process | 0.793193920138159 |
| GO:0006979 | response to oxidative stress | 0.582480485947453 |
| GO:0006486 | protein glycosylation | 0.522109647487177 |
| GO:0006487 | protein N-linked glycosylation | 0.113839201992936 |
| GO:0006284 | base-excision repair | 0.298165436998951 |
| GO:0009873 | ethylene-activated signaling pathway | 0.037081698040226 |
| GO:0006505 | GPI anchor metabolic process | 0.153101933403468 |
| GO:0009864 | induced systemic resistance, jasmonic acid mediated signaling pathway | 0.001012032529782 |

### 2 Supplementary Figures

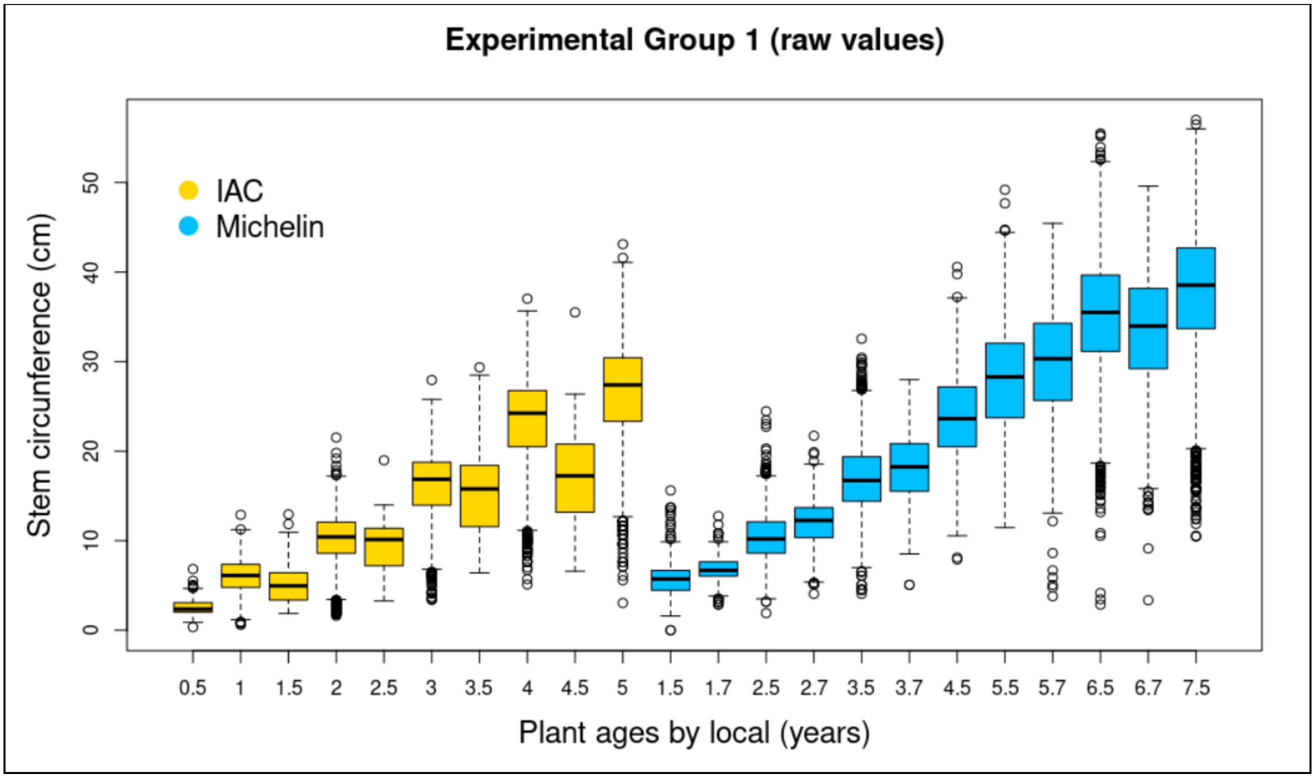


**Supplementary Fig. 1.** Distribution of raw phenotypic data in Experimental Group 1 (EG1) for stem circumference (SC) separated according to the location and the year of measurement.

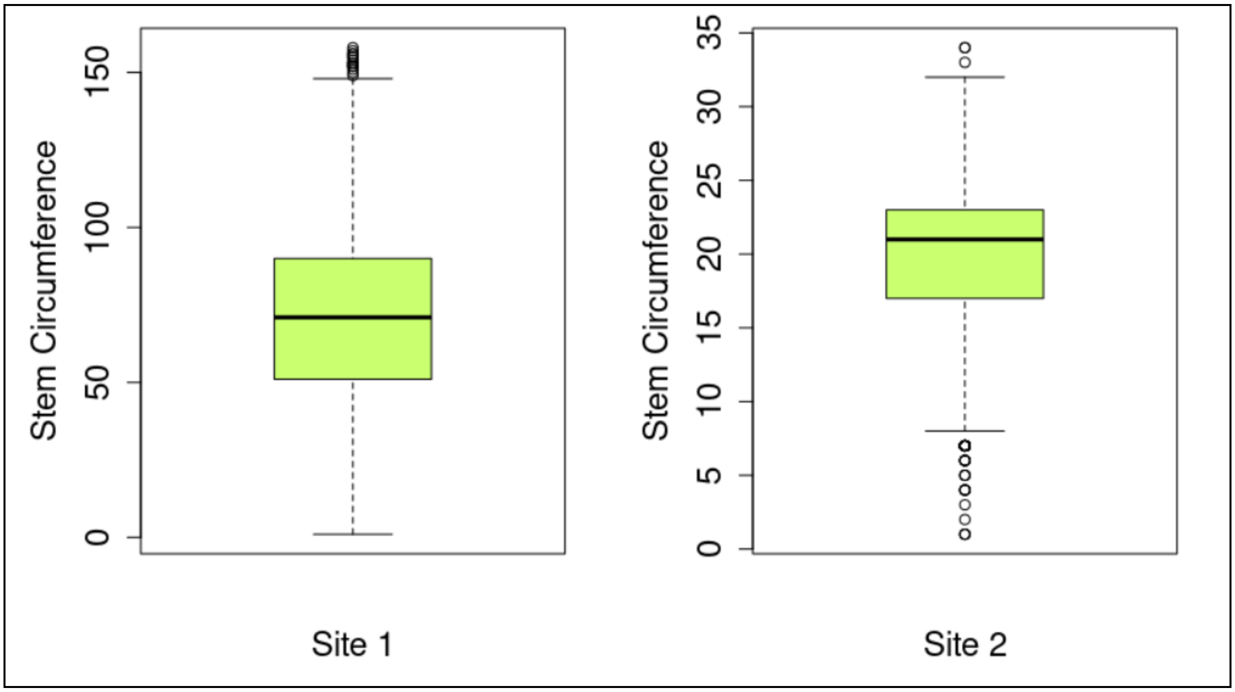


**Supplementary Fig. 2.** Distribution of raw phenotypic data in Experimental Group 2 (EG2) for stem circumference (SC) at sites 1 and 2.

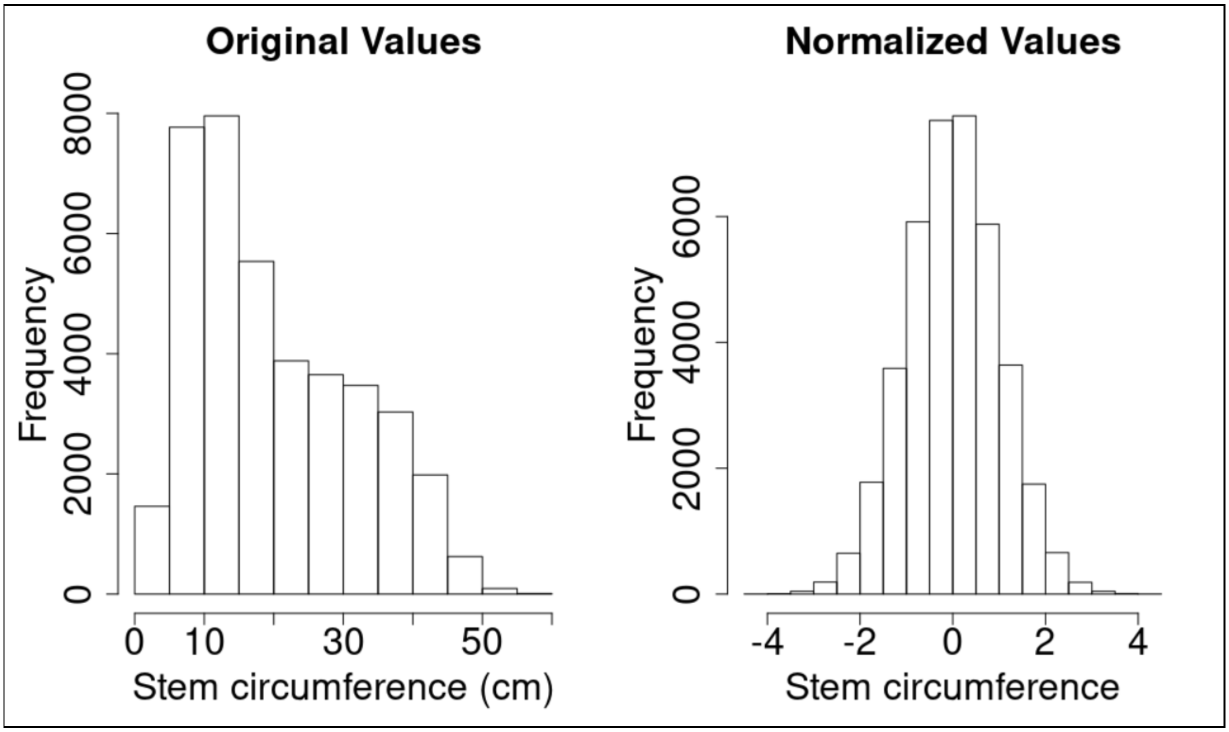


**Supplementary Fig. 3.** Distribution of trait data before and after normalization in Experimental Group 1 (EG1) for stem circumference (SC).

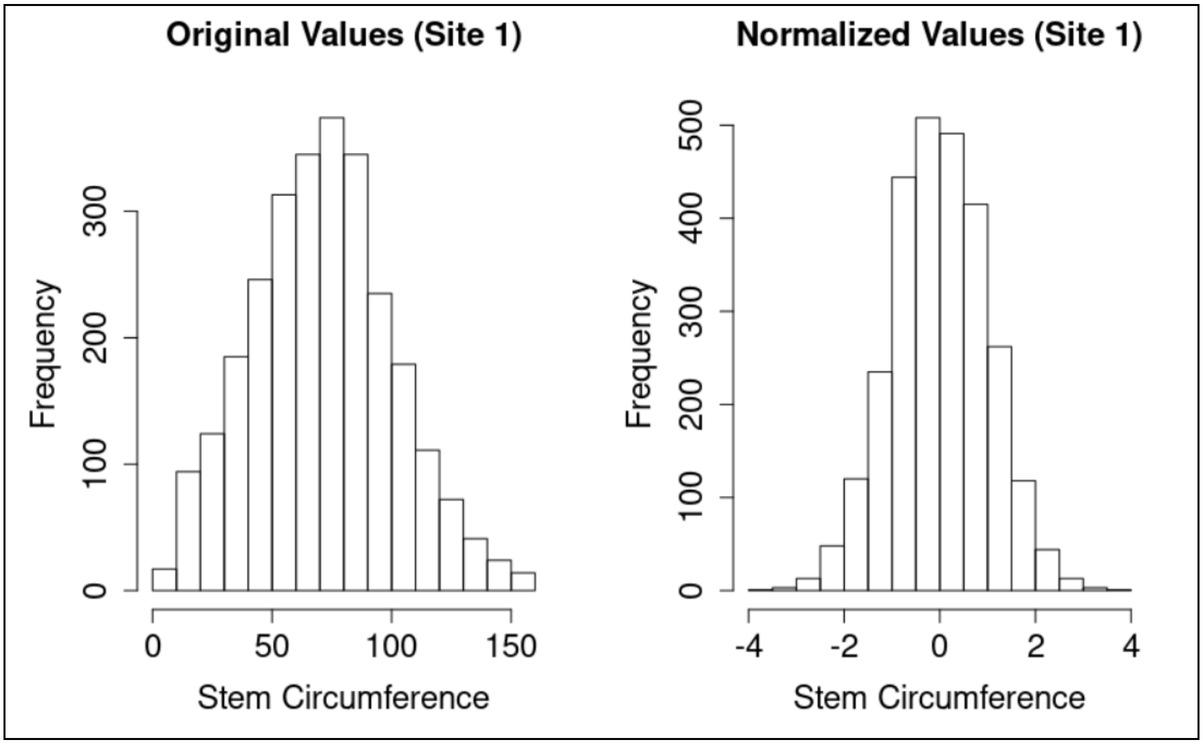


**Supplementary Fig. 4.** Distribution of trait data before and after normalization in Experimental Group 2 (EG2) for stem circumference (SC) at site 1 (S1).

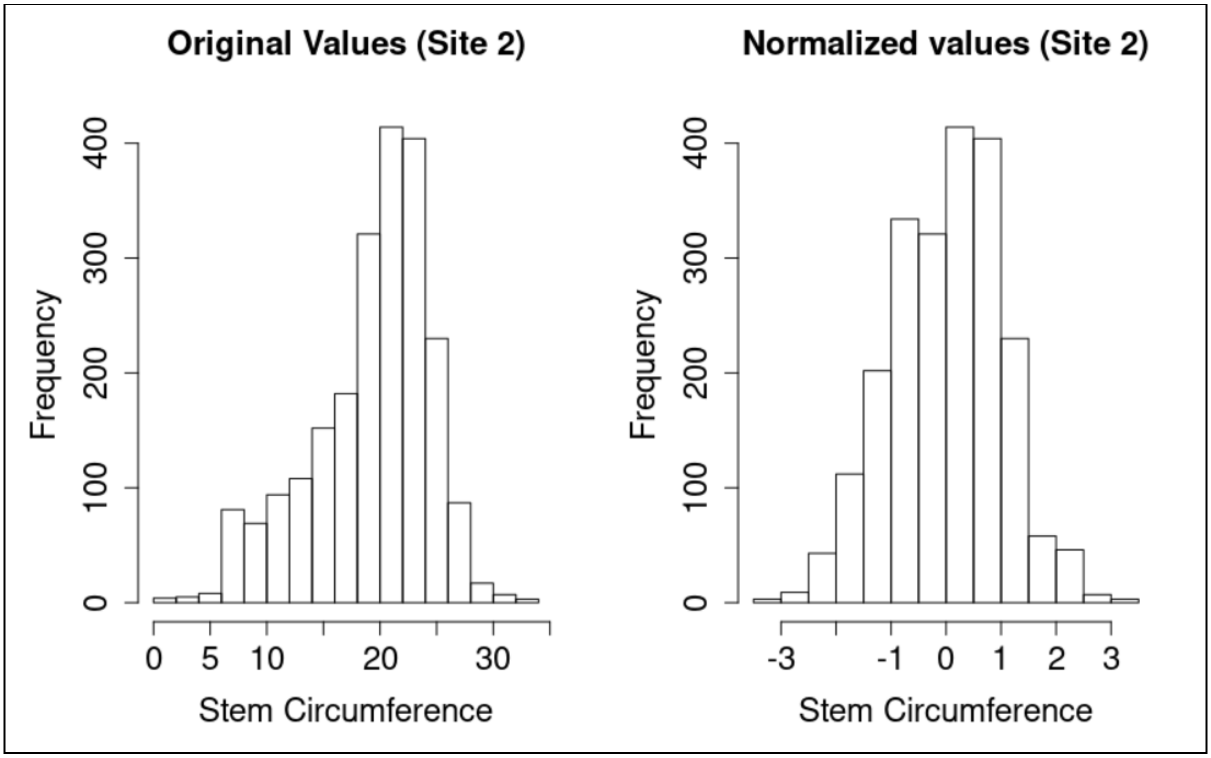


**Supplementary Fig. 5.** Distribution of trait data before and after normalization in Experimental Group 2 (EG2) for stem circumference (SC) at site 2 (S2).

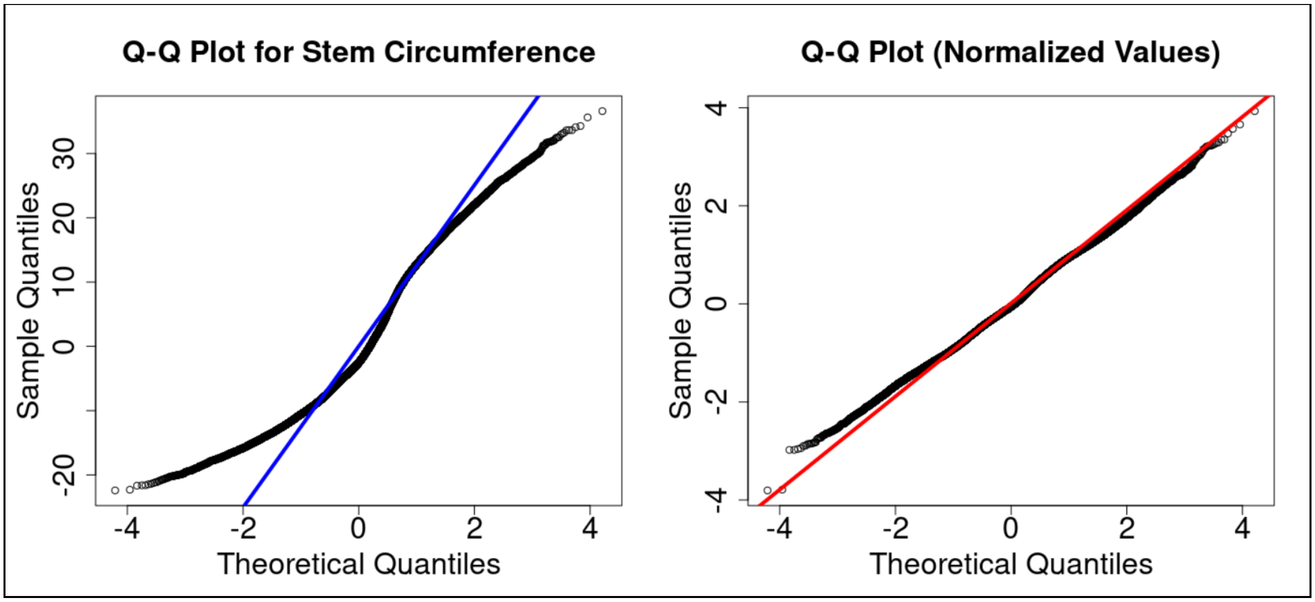


**Supplementary Fig. 6.** Quantile-quantile (Q-Q) plots for the residual distributions of the mixed models created for stem circumference (SC) in Experimental Group 1 (EG1).

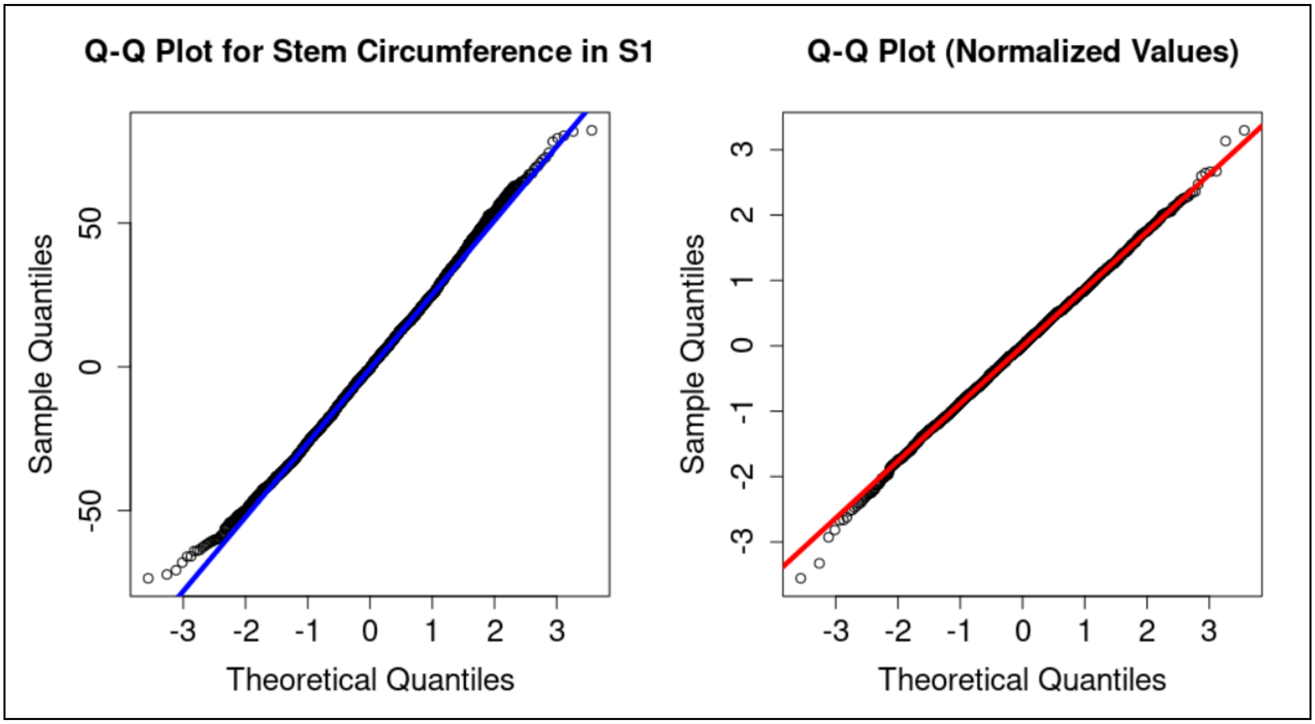


**Supplementary Fig. 7.** Quantile-quantile (Q-Q) plots for the residual distributions of the mixed models created for stem circumference (SC) in Experimental Group 2 (EG2) at site 1 (S1).

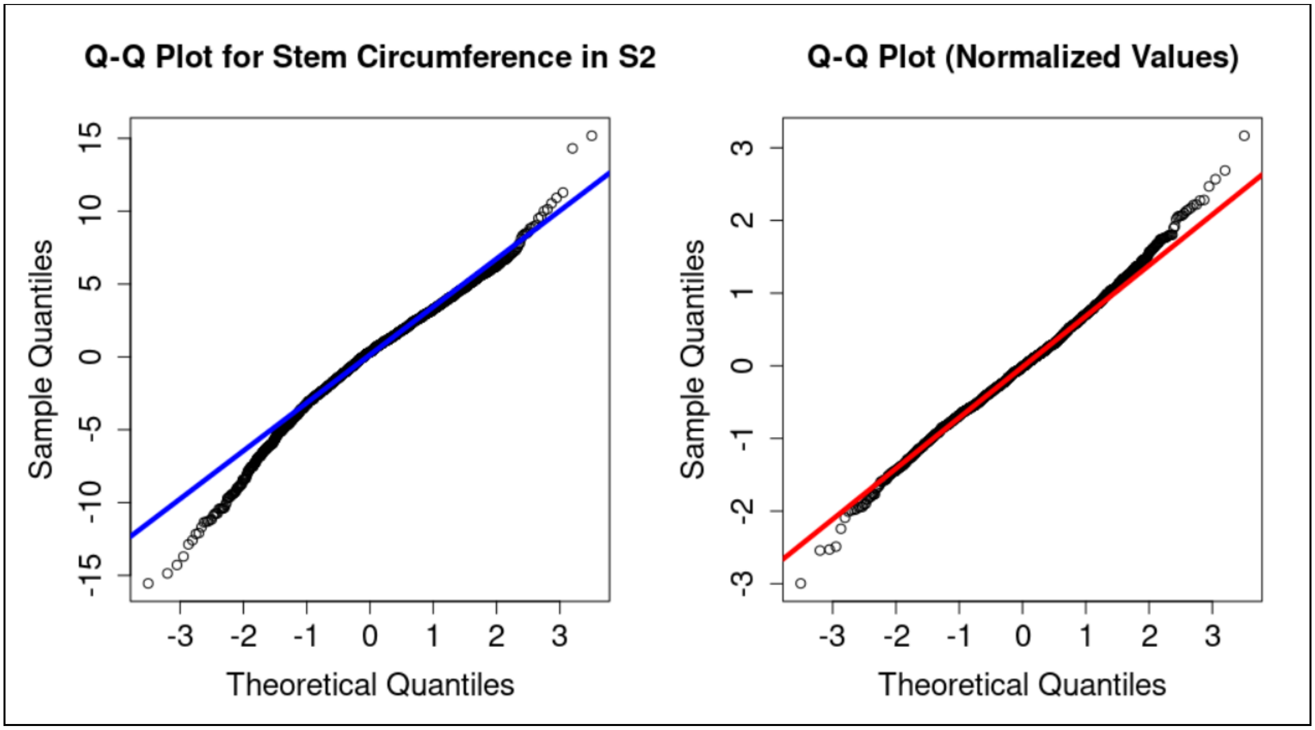


**Supplementary Fig. 8.** Quantile-quantile (Q-Q) plots for the residual distributions of the mixed models created for stem circumference (SC) in Experimental Group 2 (EG2) at site 2 (S2).

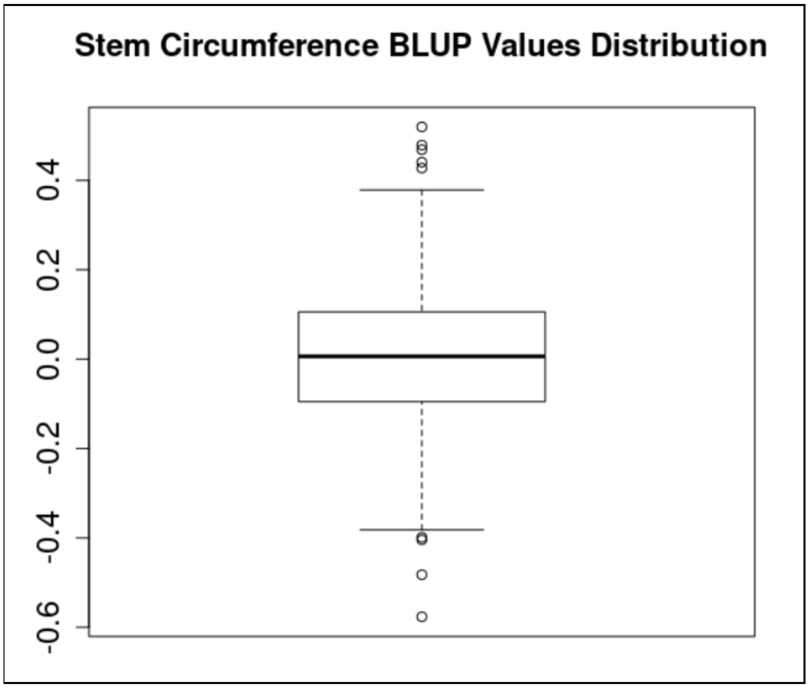


**Supplementary Fig. 9.** Distribution of estimated BLUP (B) values in Experimental Group 1 (EG1) for stem circumference (SC).

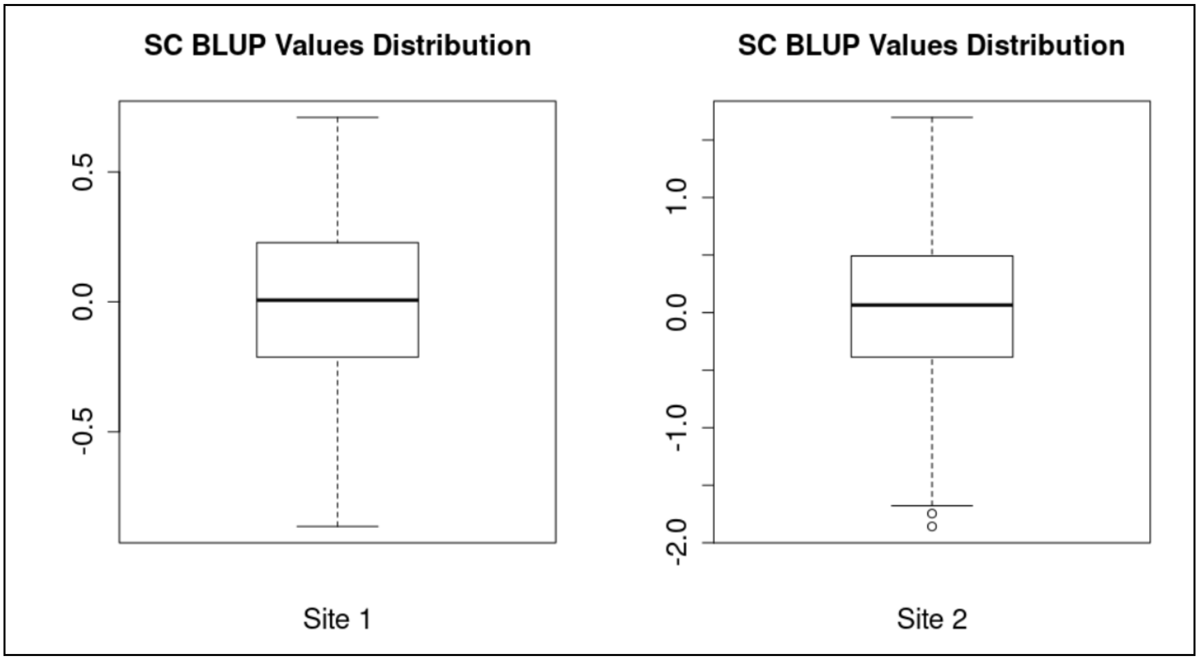


**Supplementary Fig. 10.** Distribution of estimated BLUP (B) values in Experimental Group 2 (EG2) for stem circumference (SC) at sites 1 and 2.

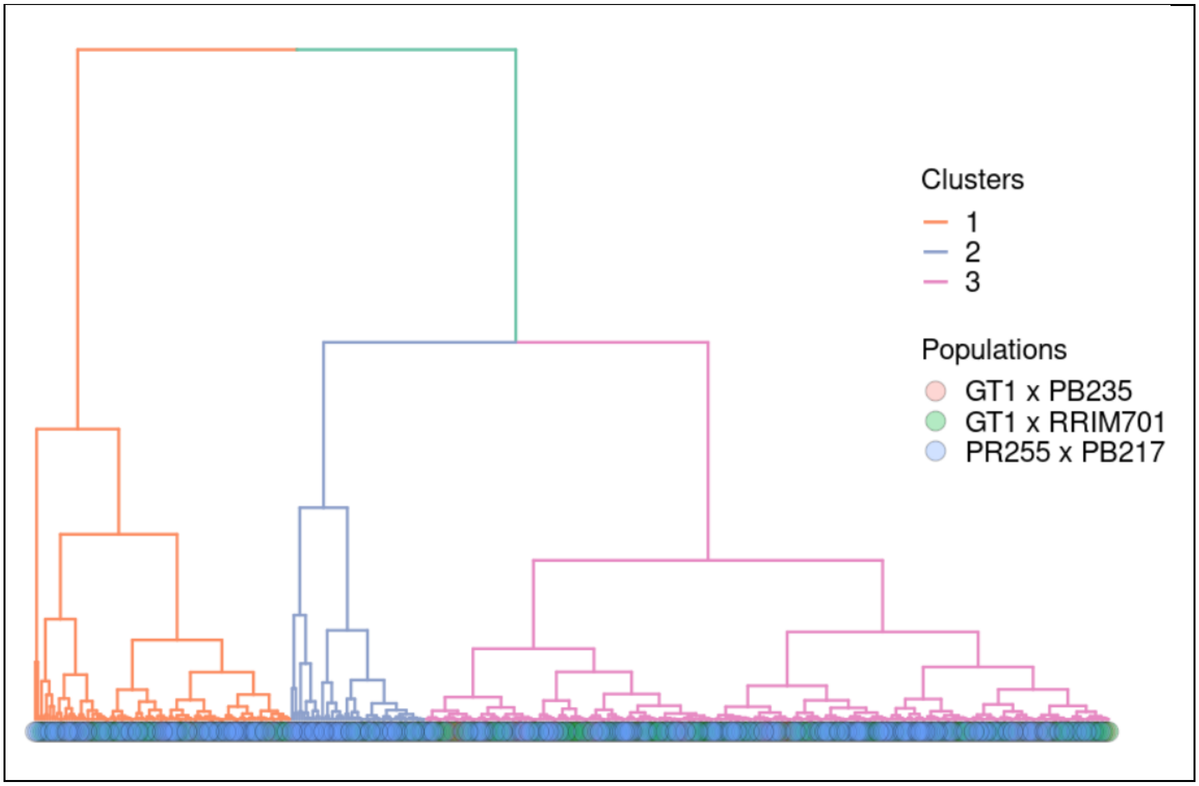


**Supplementary Fig. 11.** Constructed dendrograms colored based on the identified phenotypic groups for the BLUP (B) values in Experimental Group 1 (EG1) for stem circumference (SC).

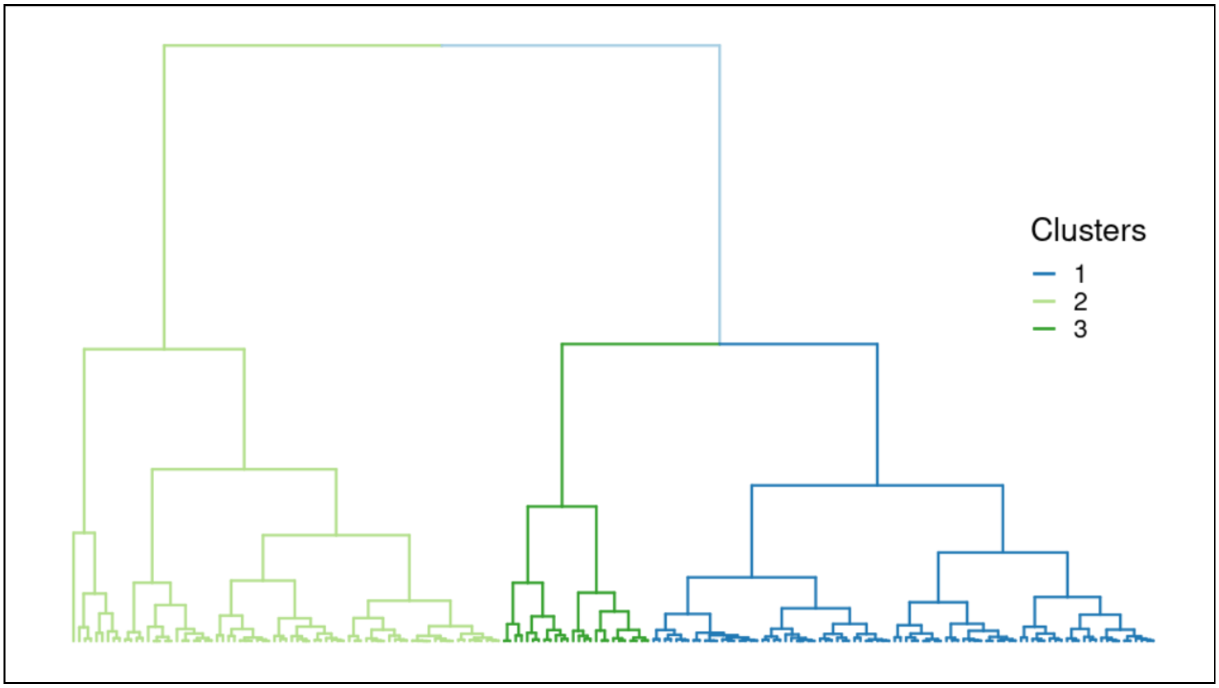


**Supplementary Fig. 12.** Constructed dendrograms colored based on the identified phenotypic groups for the BLUP (B) values in Experimental Group 2 (EG2) for stem circumference (SC) at site 1 (S1).

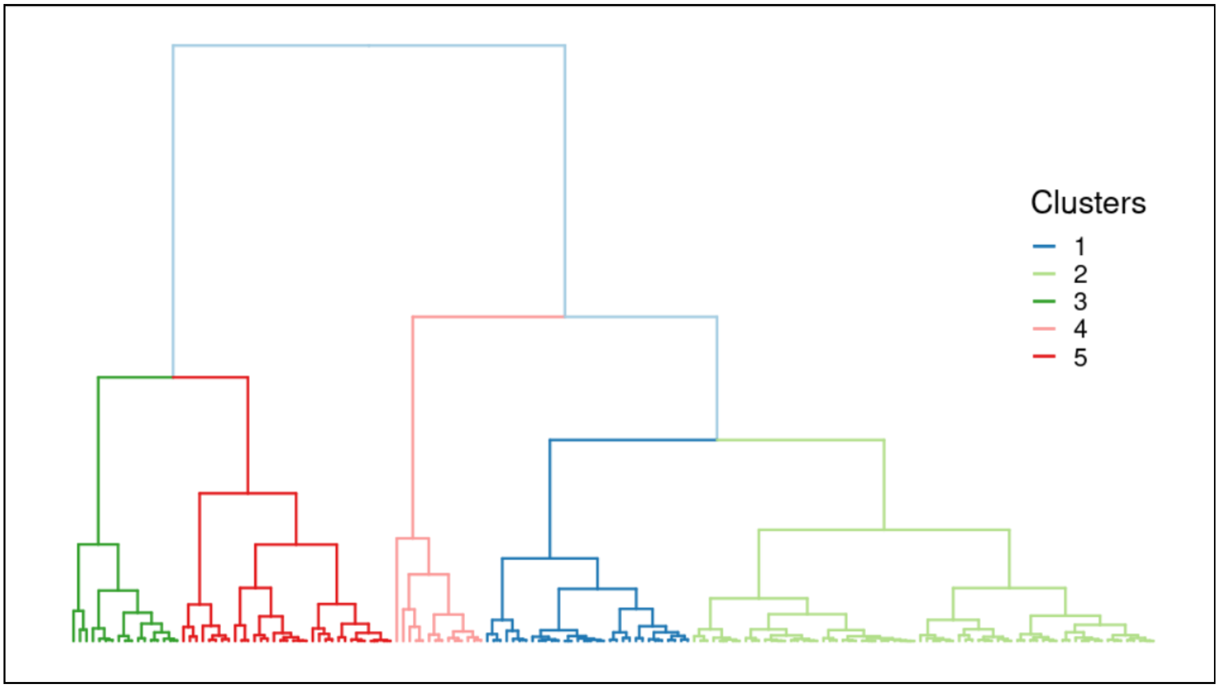


**Supplementary Fig. 13.** Constructed dendrograms colored based on the identified phenotypic groups for the BLUP (B) values in Experimental Group 2 (EG2) for stem circumference (SC) at site 2 (S2).

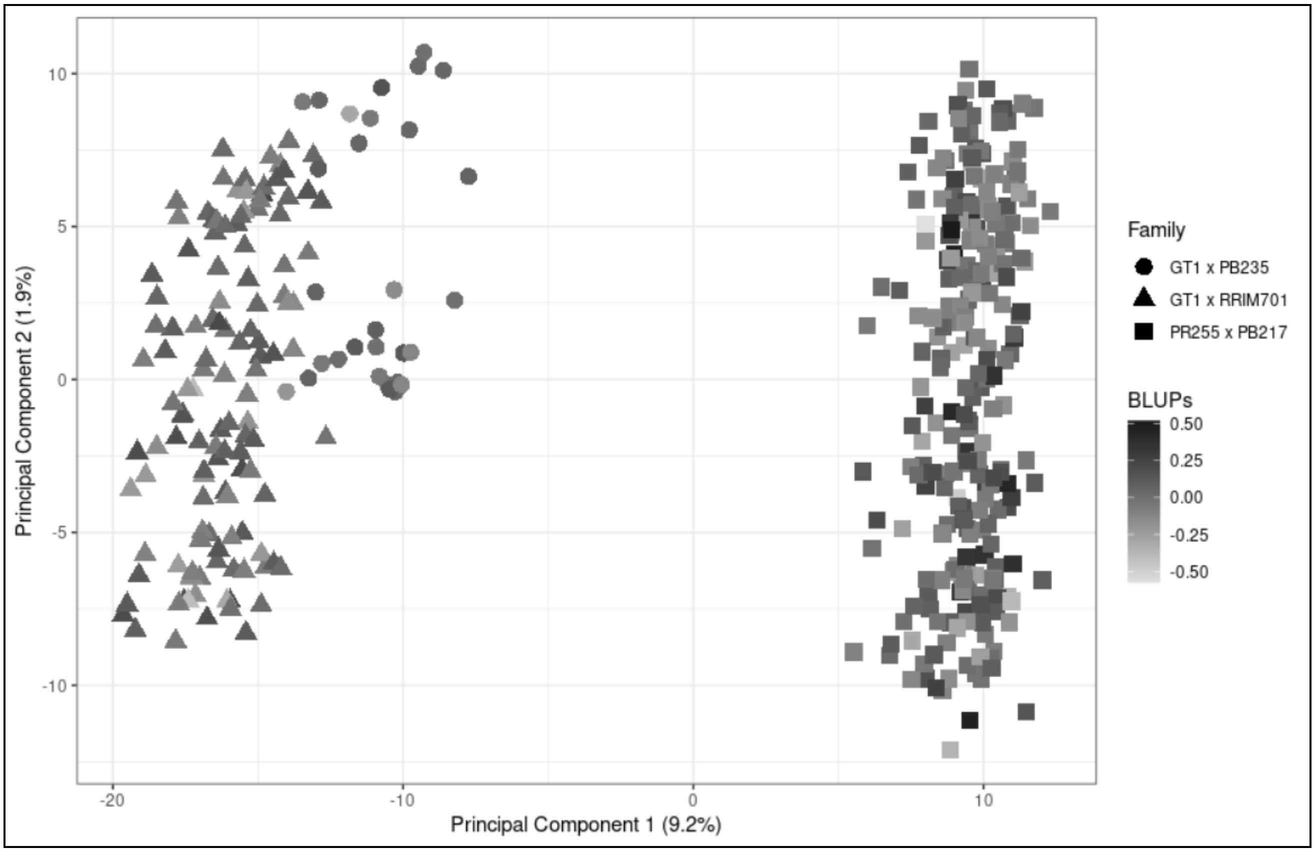


**Supplementary Fig. 14.** Principal component analysis (PCA) performed on SNP markers and colored based on stem circumference (SC) BLUPs. Each point represents an individual of experimental group 1 (EG1), belonging to a specific family (point shapes) and with a different BLUP value (intensity of gray).

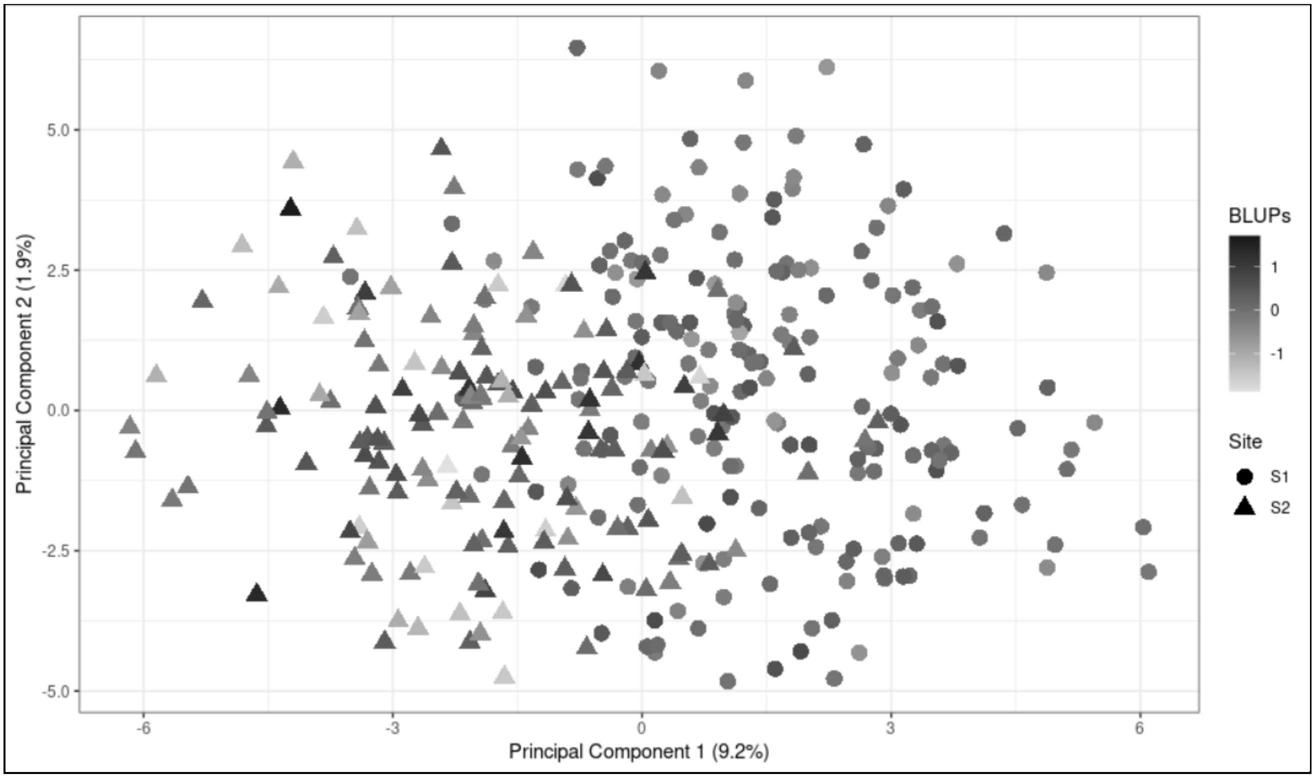


**Supplementary Fig. 15.** Principal component analysis (PCA) performed on SSR markers and colored based on stem circumference (SC) BLUPs. Each point represents an individual of experimental group 2 (EG2) at sites 1 (S1) and 2 (S2) with a different BLUP value (intensity of gray).

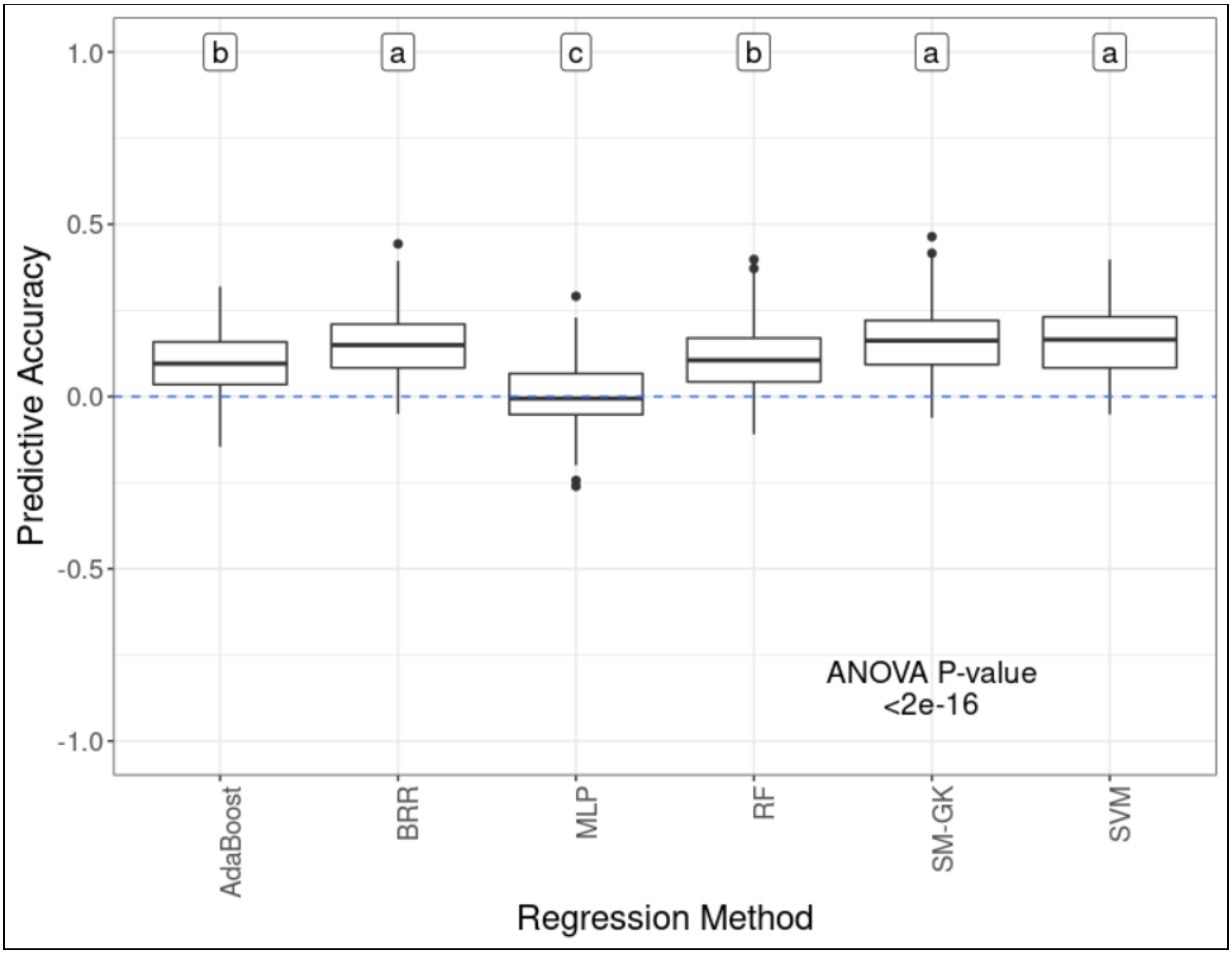


**Supplementary Fig. 16**. Predictive accuracies for stem circumference BLUP prediction in experimental group 1 (EG1) considering a 4-fold cross validation scheme (repeated 50 times), a single-environment model with a nonlinear Gaussian kernel (SM-GK), Bayesian ridge regression (BRR), and several machine learning algorithms for regression (AdaBoost, multilayer perceptron (MLP), random forest (RF), and support vector machine (SVM)). The letters at the top indicate the results from Tukey's multiple comparison test.

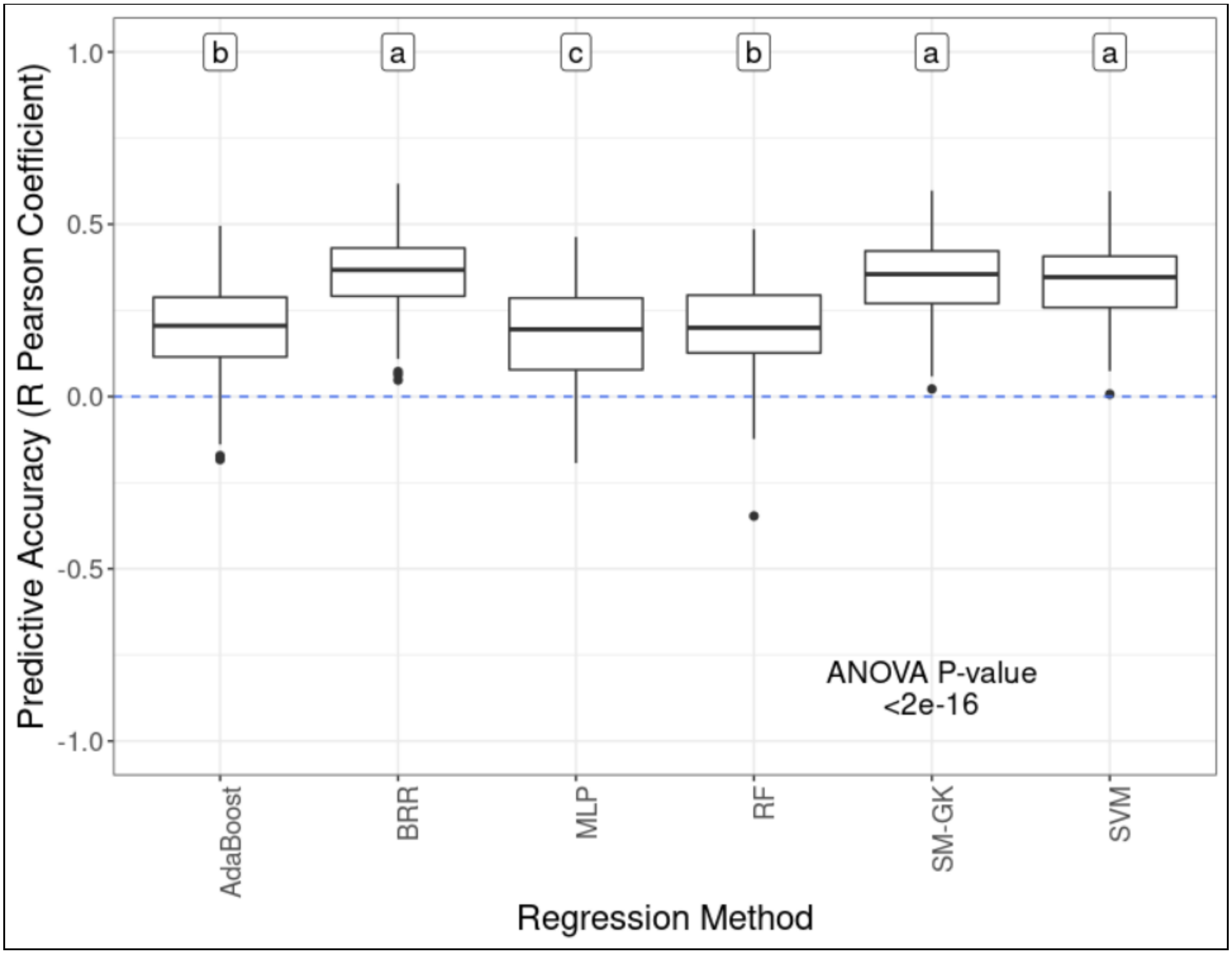


**Supplementary Fig. 17**. Predictive accuracies for stem circumference BLUP prediction in experimental group 2 (EG2) at site 1 (S1) considering a 4-fold cross validation scheme (repeated 50 times), a single-environment model with a nonlinear Gaussian kernel (SM-GK), Bayesian ridge regression (BRR), and several machine learning algorithms for regression (AdaBoost, multilayer perceptron (MLP), random forest (RF), and support vector machine (SVM)). The letters at the top indicate the results from Tukey's multiple comparison test.

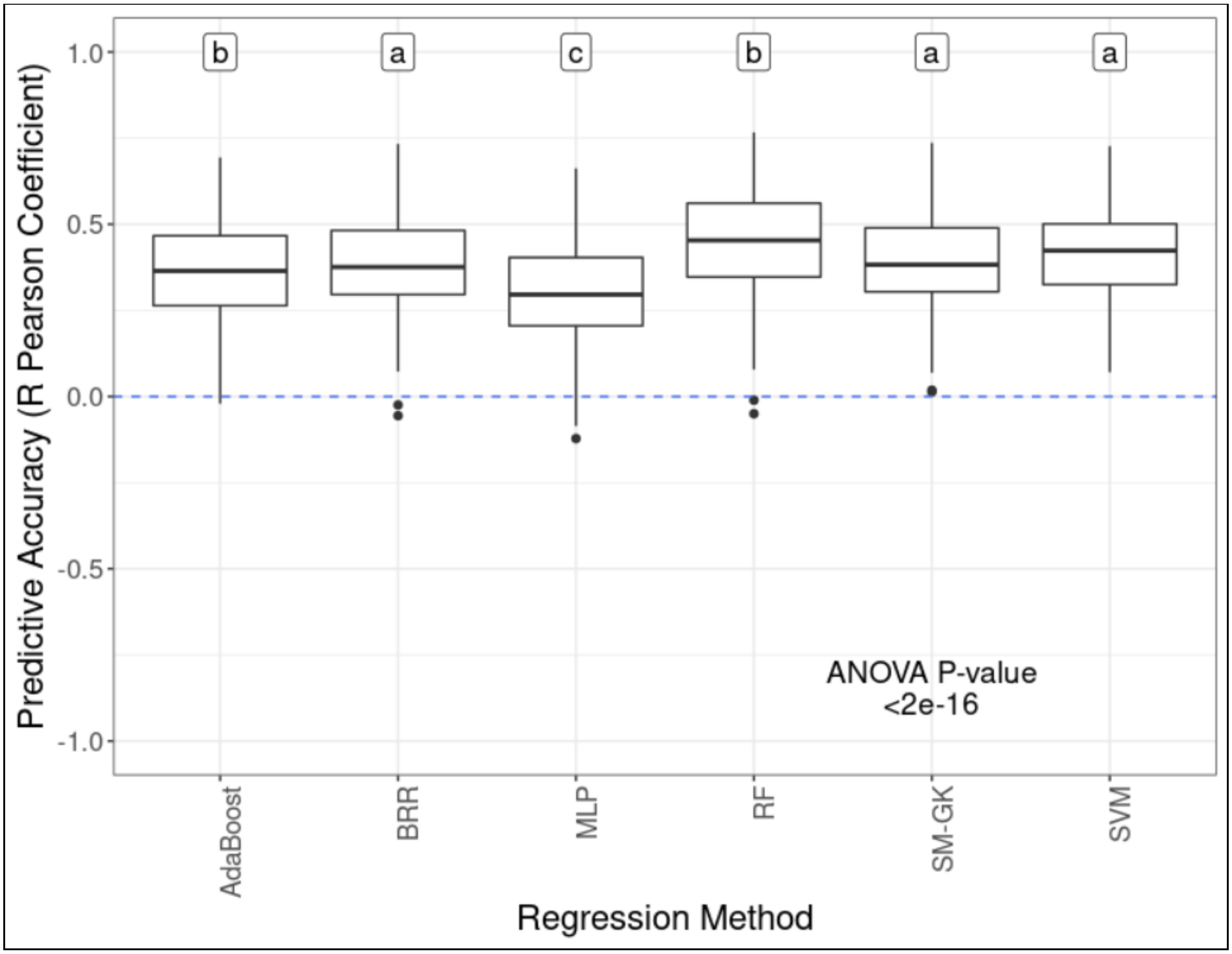


**Supplementary Fig. 18**. Predictive accuracies for stem circumference BLUP prediction in experimental group 2 (EG2) at site 2 (S2) considering a 4-fold cross validation scheme (50 times repeated), a single-environment model with a nonlinear Gaussian kernel (SM-GK), Bayesian ridge regression (BRR), and several machine learning algorithms for regression (AdaBoost, multilayer perceptron (MLP), random forest (RF), and support vector machine (SVM)). The letters at the top indicate the results from Tukey's multiple comparison test.

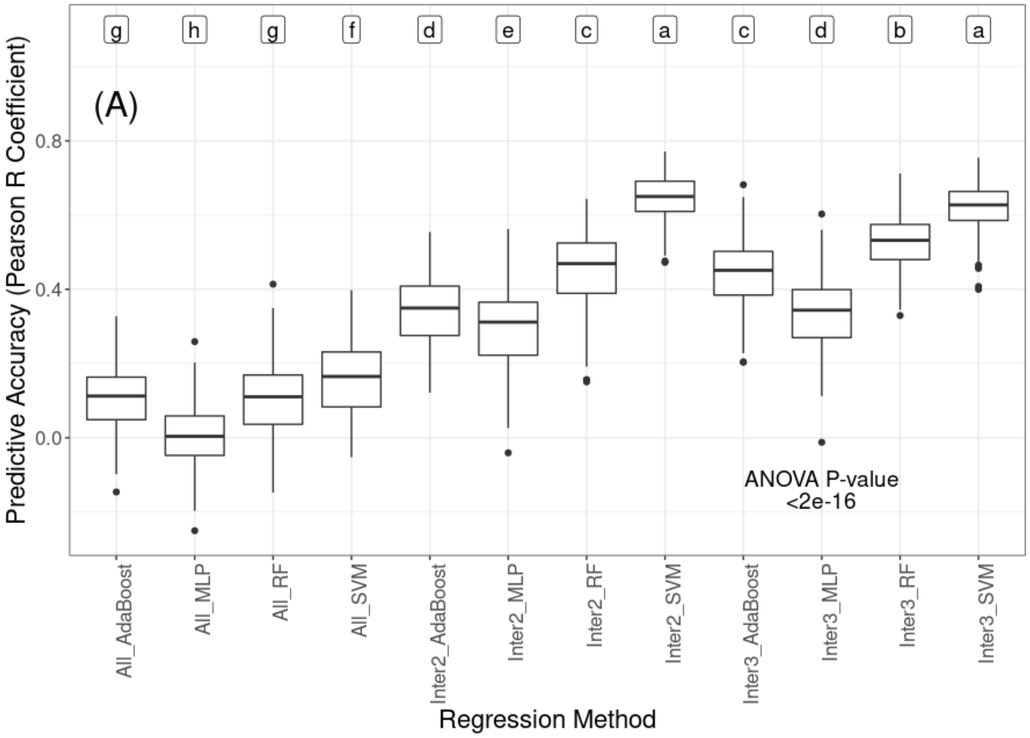
(A)
g h g f d e c a c d b a
Predictive Accuracy (Pearson R Coefficient)
0.8
0.4
0.0
ANOVA P-value
<2e-16
All_AdaBoost
All_MLP
All_RF
All_SVM
Inter2_AdaBoost
Inter2_MLP
Inter2_RF
Inter2_SVM
Inter3_AdaBoost
Inter3_MLP
Inter3_RF
Inter3_SVM
Regression Method

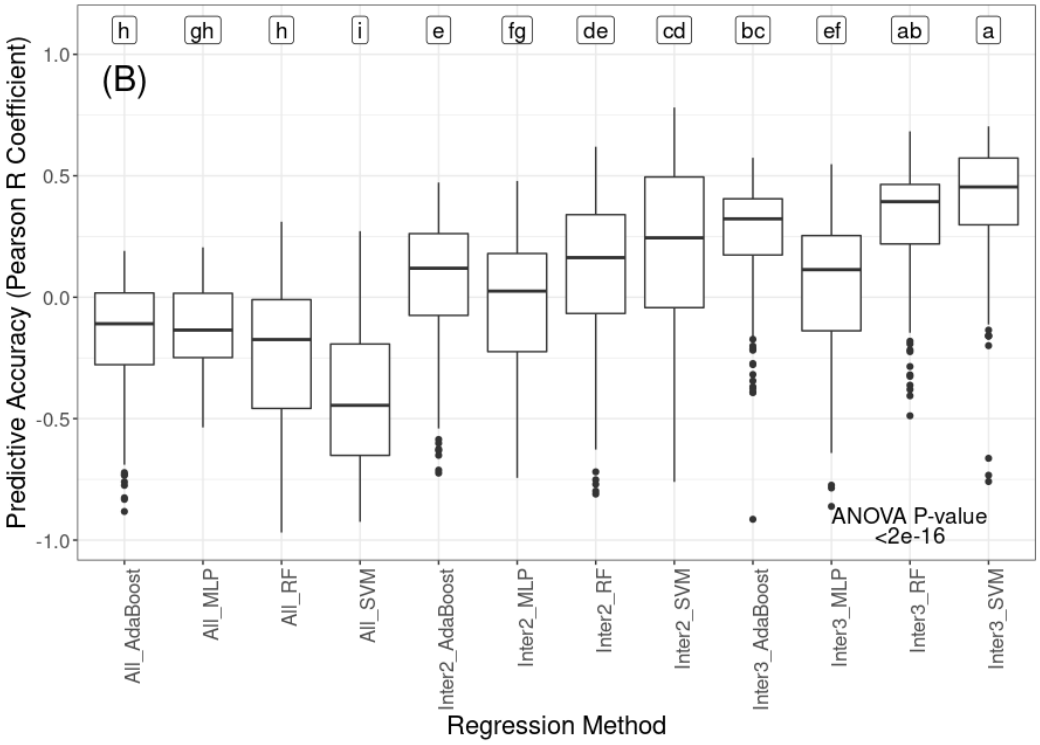
(B)
h gh h i e fg de cd bc ef ab a
Predictive Accuracy (Pearson R Coefficient)
1.0
0.5
0.0
-0.5
-1.0
ANOVA P-value
<2e-16
All_AdaBoost
All_MLP
All_RF
All_SVM
Inter2_AdaBoost
Inter2_MLP
Inter2_RF
Inter2_SVM
Inter3_AdaBoost
Inter3_MLP
Inter3_RF
Inter3_SVM
Regression Method

**Supplementary Fig. 19**. Predictive accuracies for stem circumference BLUP prediction in experimental group 1 (EG1) considering a 4-fold cross validation scheme (repeated 50 times), different machine learning algorithms for regression (AdaBoost, multilayer perceptron (MLP), random forest (RF), and support vector machine (SVM)), and feature selection (nonincorporation (All), intersection among the three methods established (Inter3), and intersection between at least two out of the three methods established (Inter2)). The letters at the top indicate the results from Tukey's multiple comparison test. In (A), the overall correlation between the predicted and real BLUPs is evaluated, and in (B), the worst correlation is evaluated considering the Pearson correlation analysis performed according to subpopulation structure.

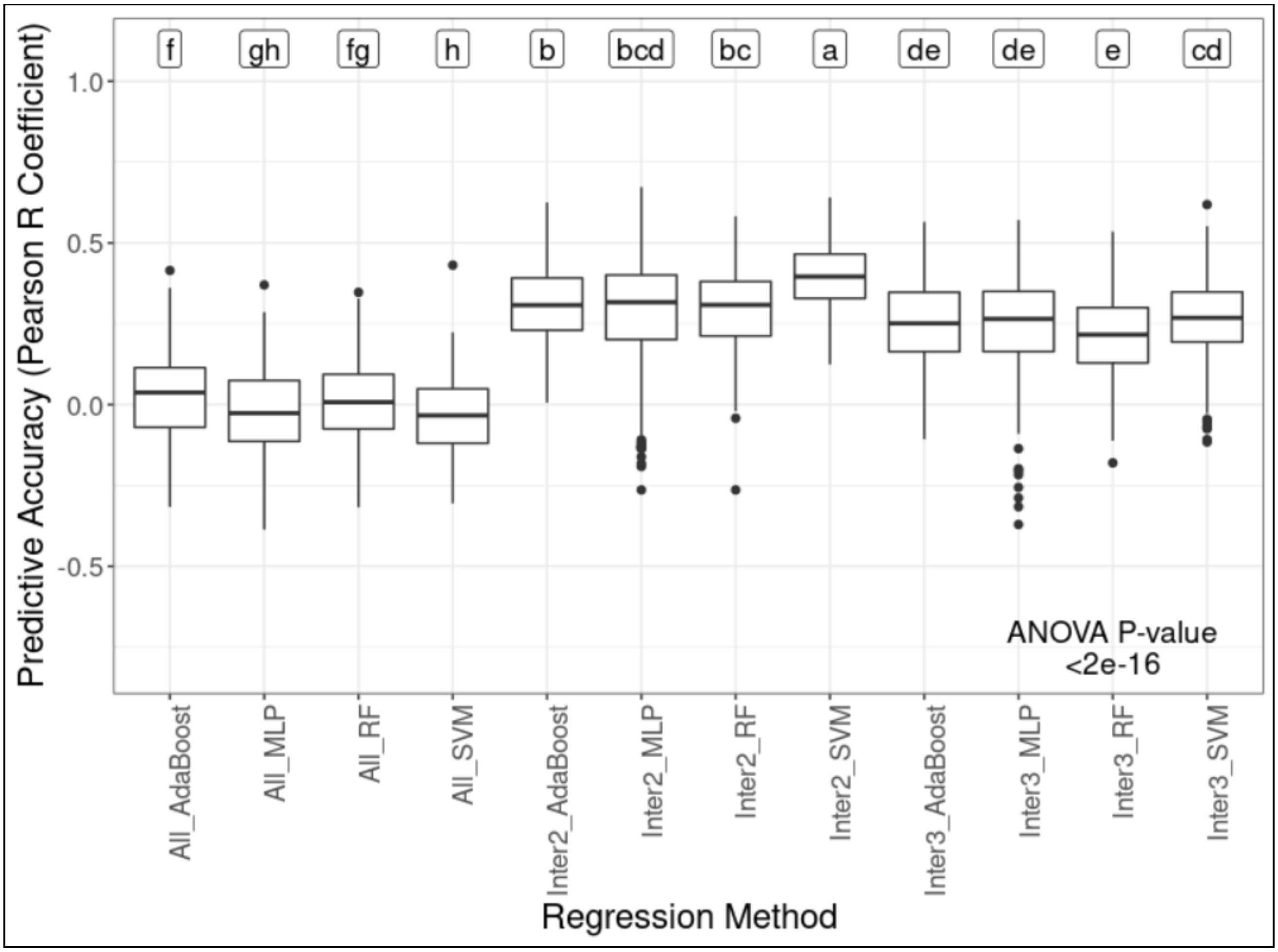


**Supplementary Fig. 20**. Predictive accuracies for stem circumference BLUP prediction in experimental group 2 (EG2) and the first site (S1) considering a 4-fold cross validation scheme (repeated 50 times), different machine learning algorithms for regression (AdaBoost, multilayer perceptron (MLP), random forest (RF), and support vector machine (SVM)), and feature selection (nonincorporation (All), intersection among the three methods established (Inter3), and intersection between at least two out of the three methods established (Inter2)). The letters at the top indicate the results from Tukey’s multiple comparison test.

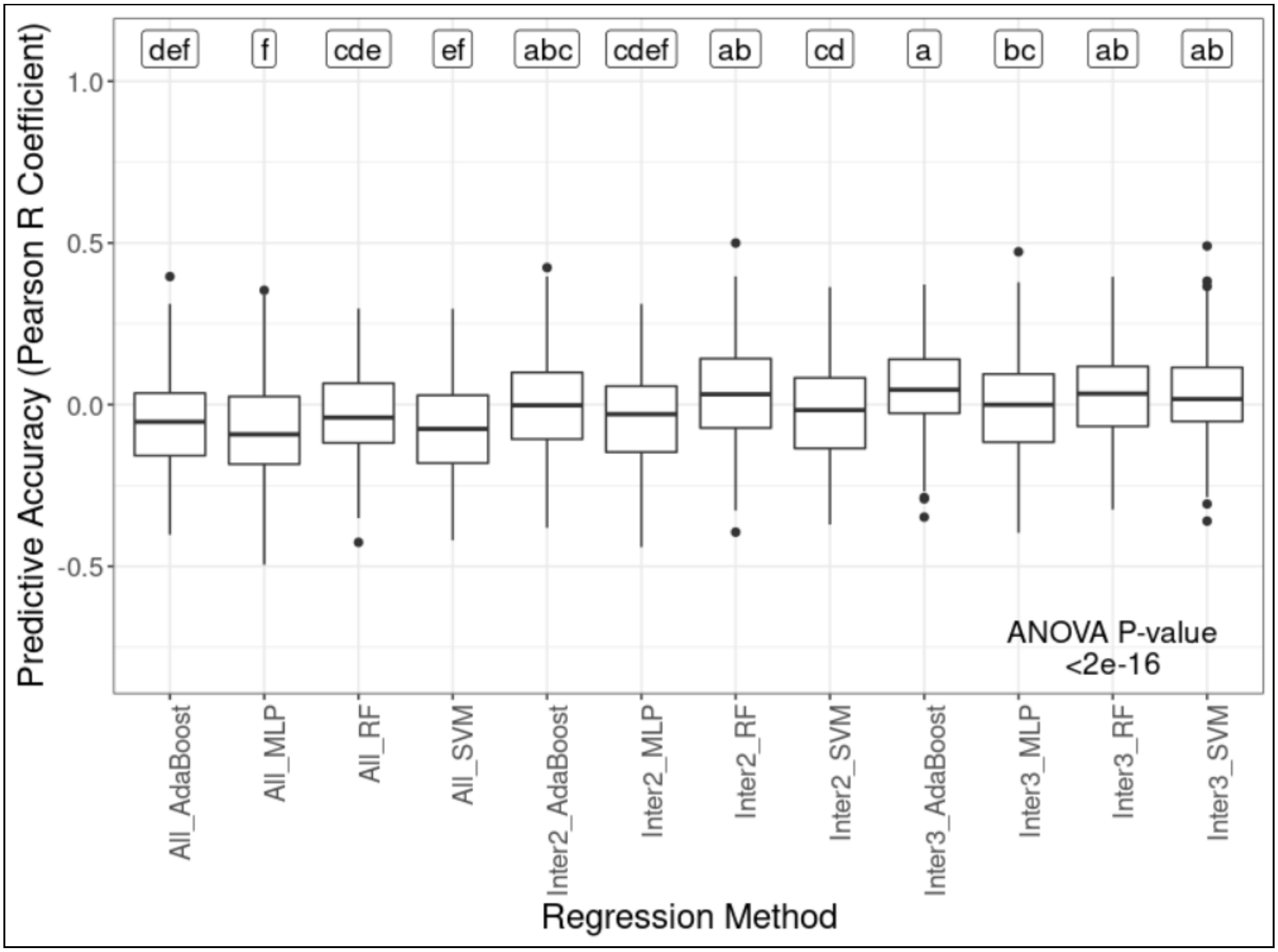


**Supplementary Fig. 21**. Predictive accuracies for stem circumference BLUP prediction in experimental group 2 (EG2) at the second site (S2) considering a 4-fold cross validation scheme (repeated 50 times), different machine learning algorithms for regression (AdaBoost, multilayer perceptron (MLP), random forest (RF), and support vector machine (SVM)), and feature selection (nonincorporation (All), intersection among the three methods established (Inter3), and intersection among at least two out of the three methods established (Inter2)). The letters at the top indicate the results from Tukey's multiple comparison test.

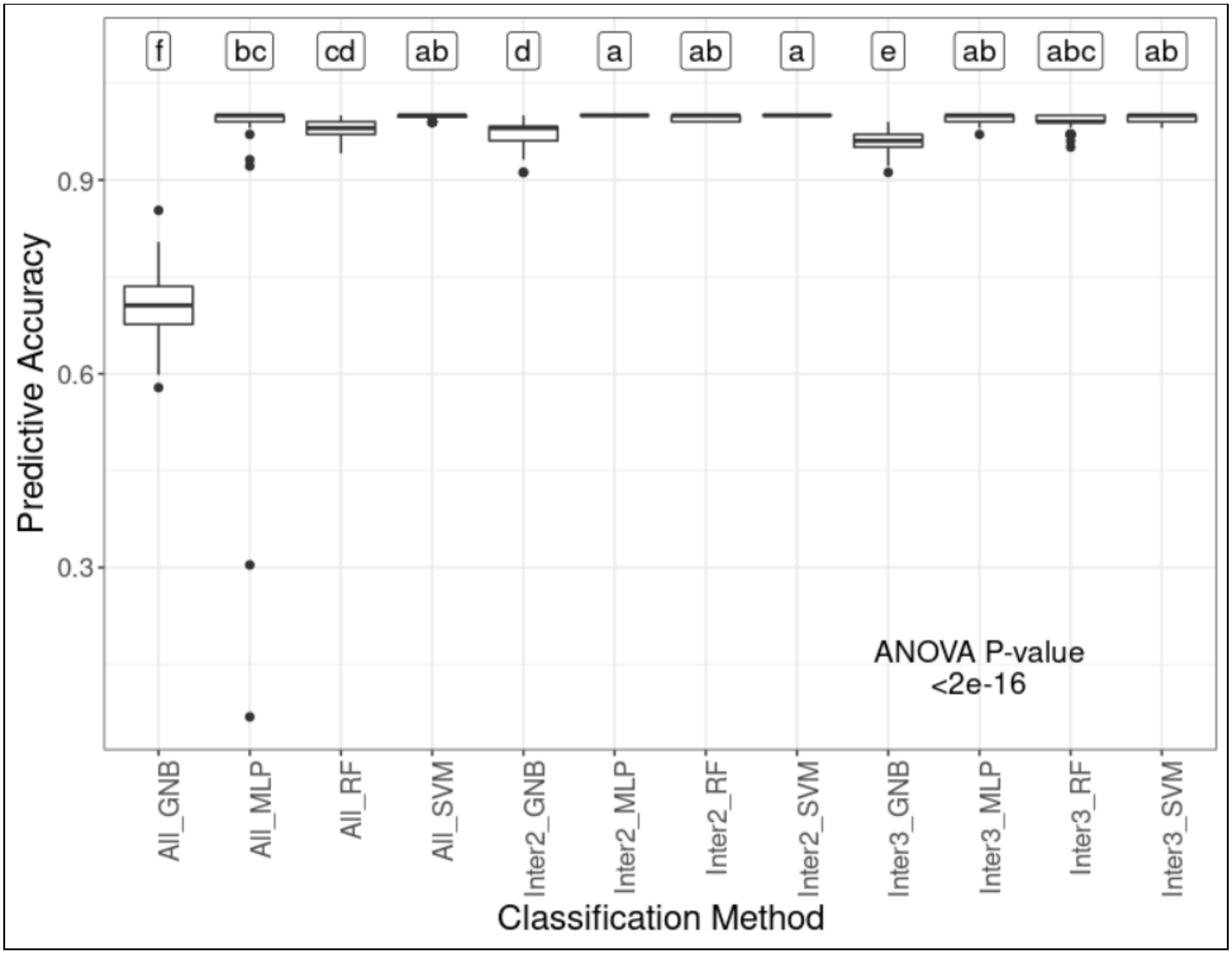


**Supplementary Fig. 22**. Predictive accuracies for population classification in experimental group 1 (EG1) considering a 4-fold cross validation scheme (repeated 50 times), different classification algorithms for prediction (Gaussian naive Bayes (GNB), multilayer perceptron (MLP), random forest (RF), and support vector machine (SVM)), and feature selection (nonincorporation (All), intersection among the three methods established (Inter3), and intersection between at least two out of the three methods established (Inter2)). The letters at the top indicate the results from Tukey's multiple comparison test.

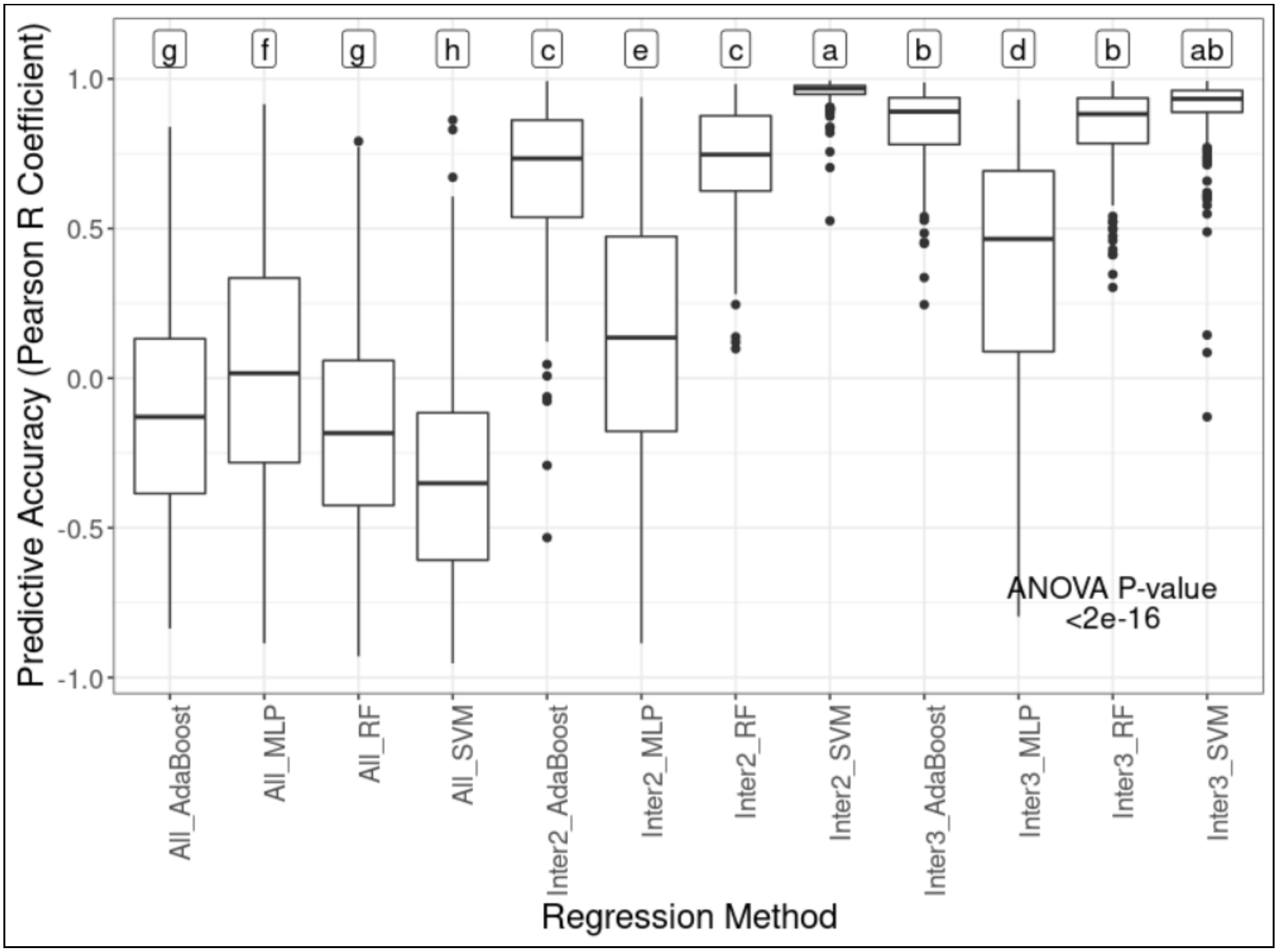


**Supplementary Fig. 23**. Predictive accuracies for stem circumference BLUP prediction in experimental group 1 (EG1) and the population GT1 x PB235 considering a 4-fold cross validation scheme (repeated 50 times), different machine learning algorithms for regression (AdaBoost, multilayer perceptron (MLP), random forest (RF), and support vector machine (SVM)), and feature selection (nonincorporation (All), intersection among the three methods established (Inter3), and intersection between at least two out of the three methods established (Inter2)). The letters at the top indicate the results from Tukey's multiple comparison test.

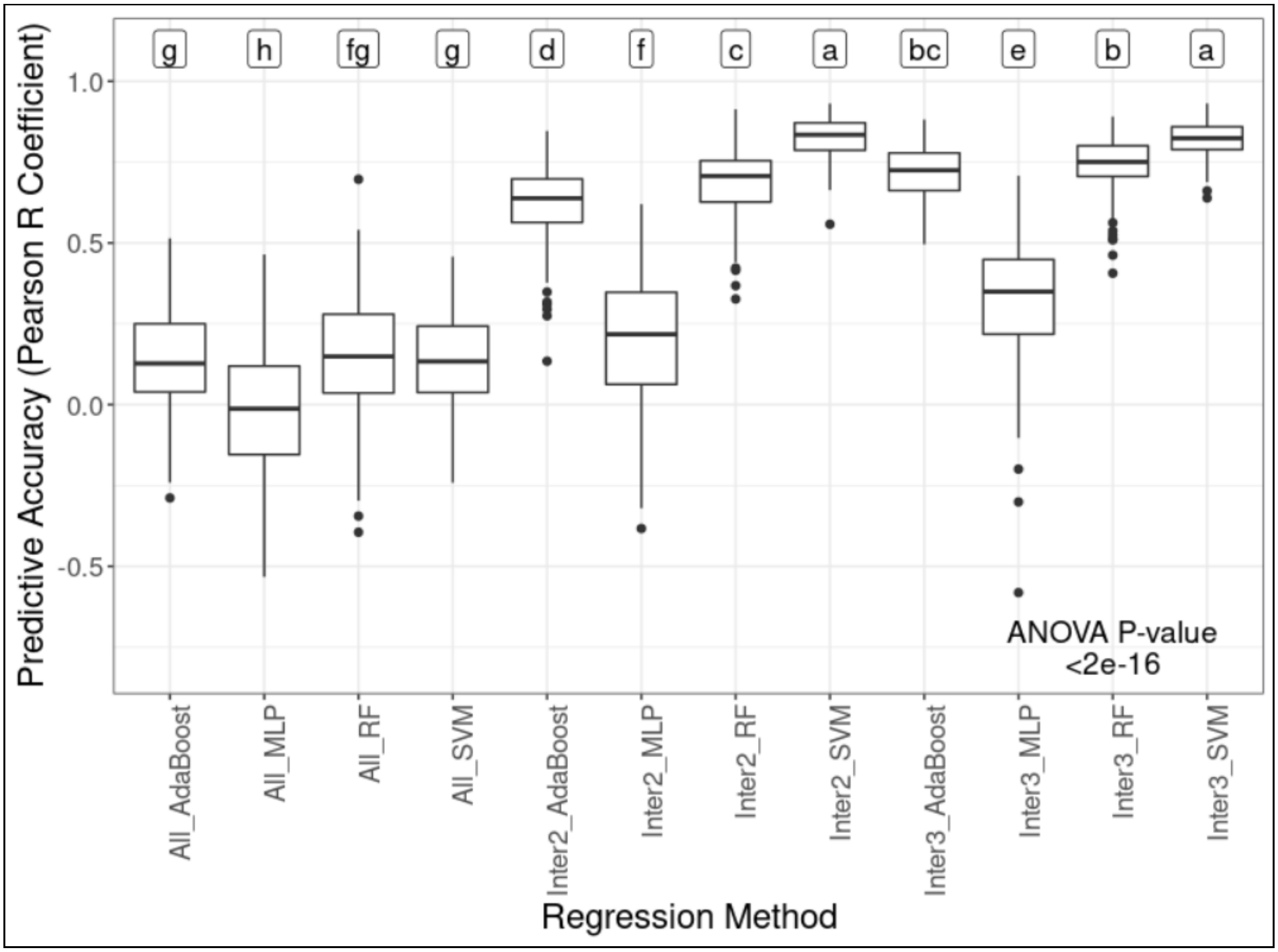


**Supplementary Fig. 24**. Predictive accuracies for stem circumference BLUP prediction in experimental group 1 (EG1) and the population GT1 x RRIM701 considering a 4-fold cross validation scheme (repeated 50 times), different machine learning algorithms for regression (AdaBoost, multilayer perceptron (MLP), random forest (RF), and support vector machine (SVM)), and feature selection (nonincorporation (All), intersection among the three methods established (Inter3), and intersection between at least two out of the three methods established (Inter2)). The letters at the top indicate the results from Tukey's multiple comparison test.

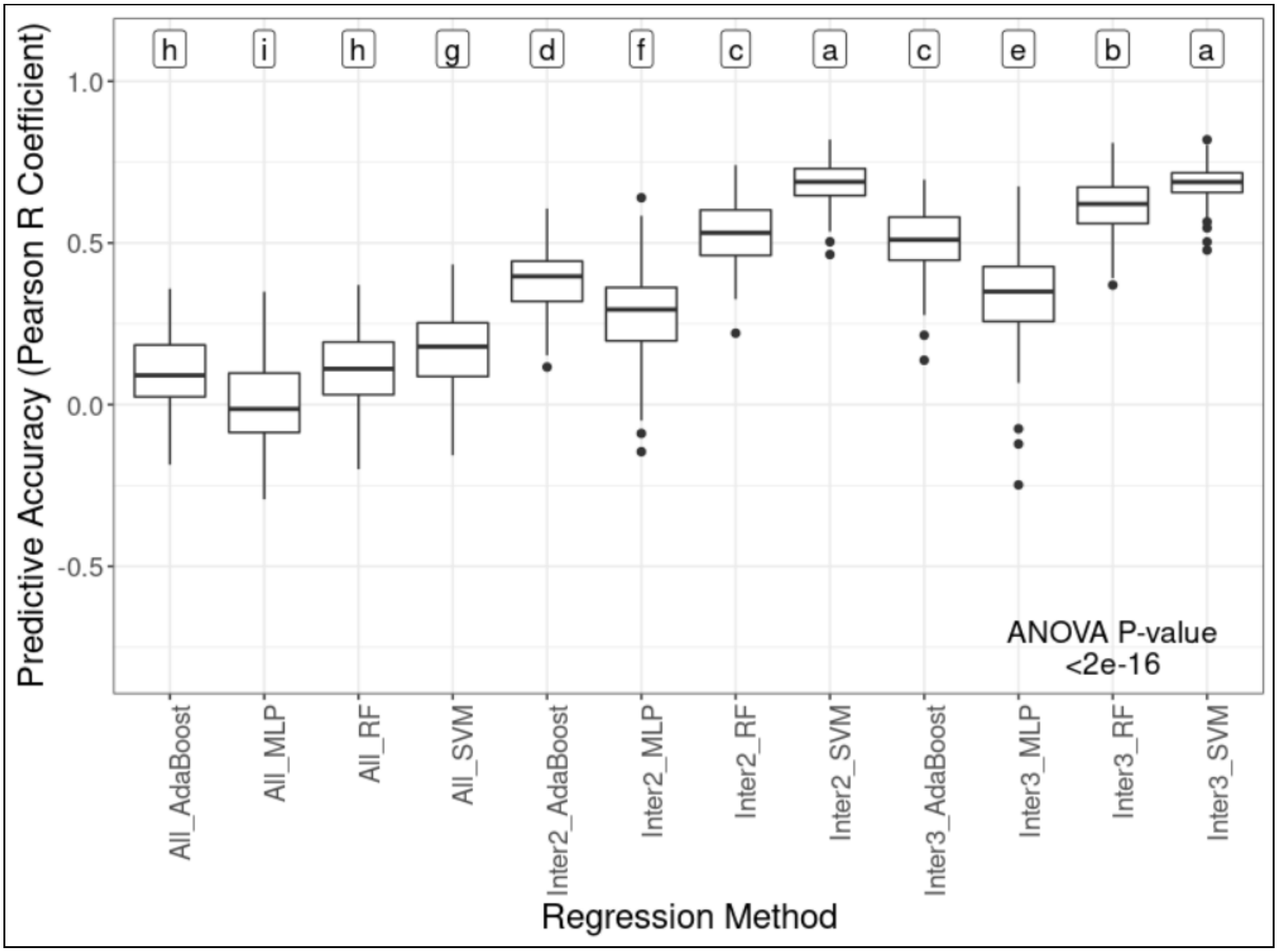


**Supplementary Fig. 25**. Predictive accuracies for stem circumference BLUP prediction in experimental group 1 (EG1) and the population PR255 x PB217 considering a 4-fold cross validation scheme (repeated 50 times), different machine learning algorithms for regression (AdaBoost, multilayer perceptron (MLP), random forest (RF), and support vector machine (SVM)), and feature selection (nonincorporation (All), intersection among the three methods established (Inter3), and intersection between at least two out of the three methods established (Inter2)). The letters at the top indicate the results from Tukey's multiple comparison test.

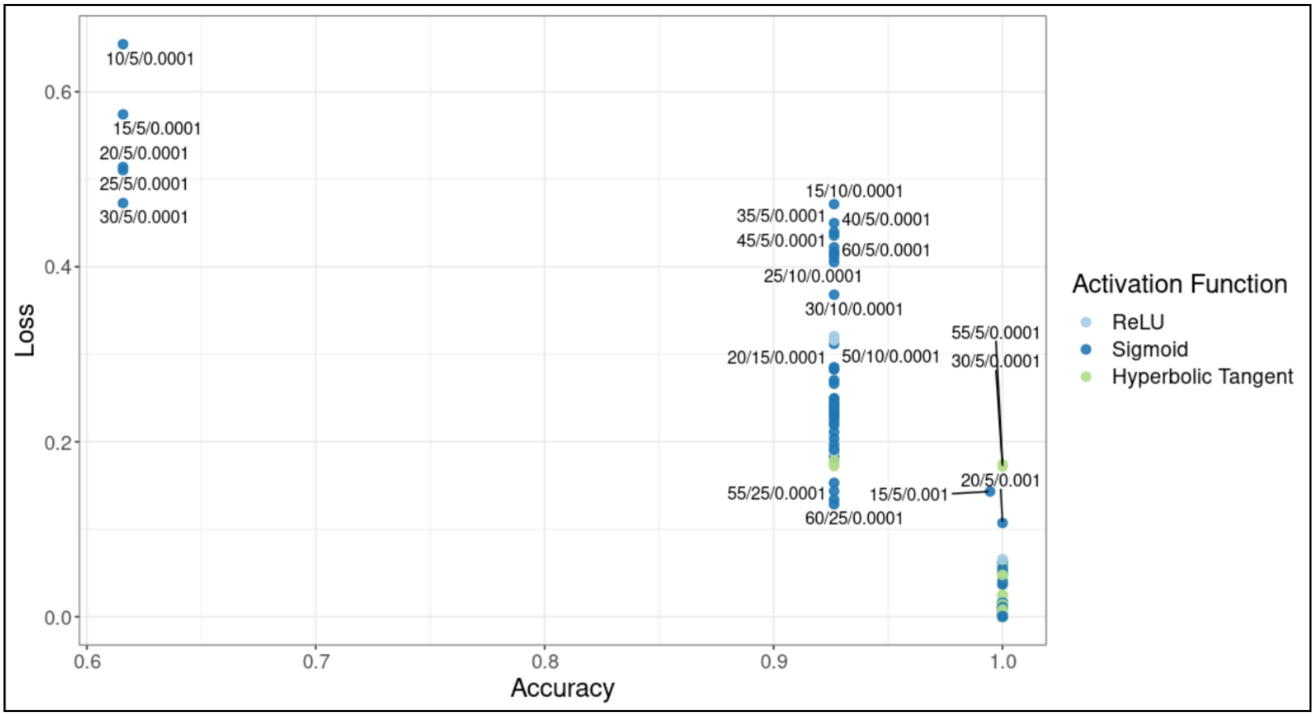


**Supplementary Fig. 26**. Hyperparameter search for subpopulation prediction in experimental group 1 (EG1). Each point represents a different combination of hyperparameters, considering (a) neurons in the first hidden layer, (b) neurons in the second hidden layer, (c) the learning rate, and (d) the activation function. These combinations were evaluated in the development set considering accuracy and the loss function evaluations.

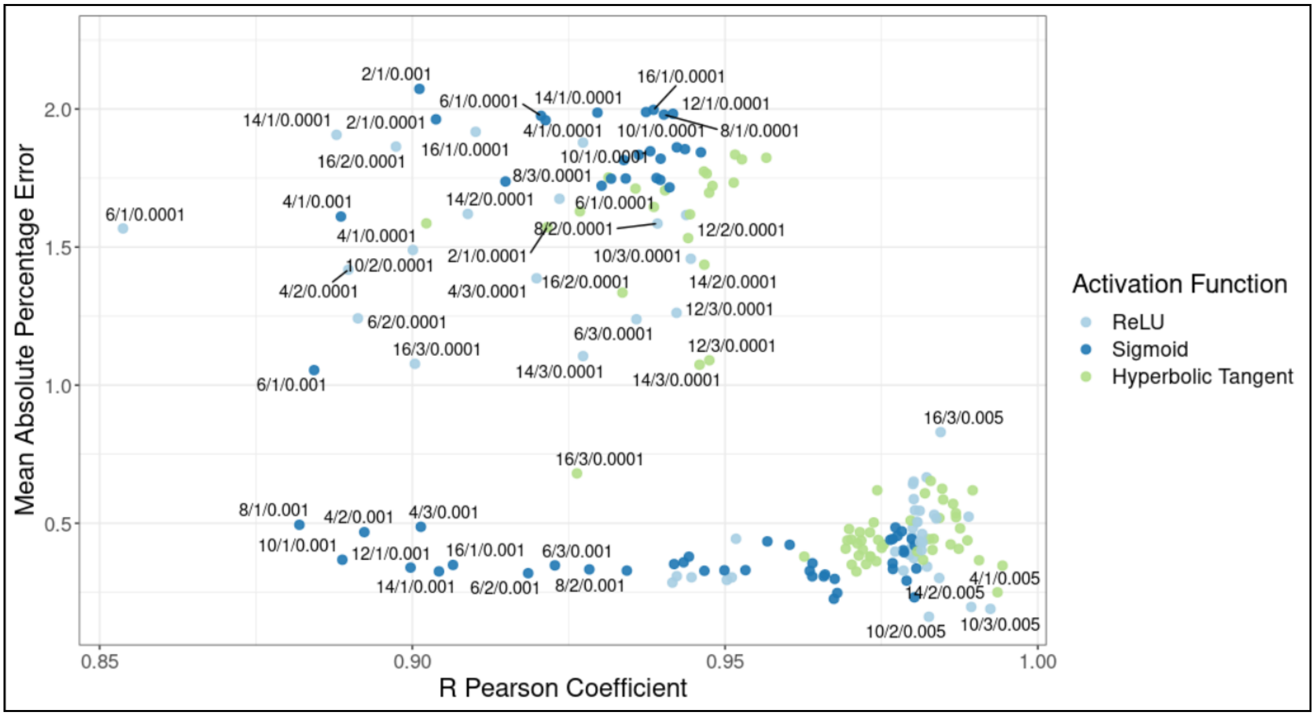


**Supplementary Fig. 27**. Hyperparameter search for stem circumference BLUP prediction in experimental group 1 (EG1) for the population GT1 x PB235. Each point represents a different combination of hyperparameters, considering (a) neurons in the first hidden layer, (b) neurons in the second hidden layer, (c) the learning rate, and (d) the activation function. These combinations were evaluated in the development set considering the mean absolute percentage error (MAPE) and the R Pearson coefficient.

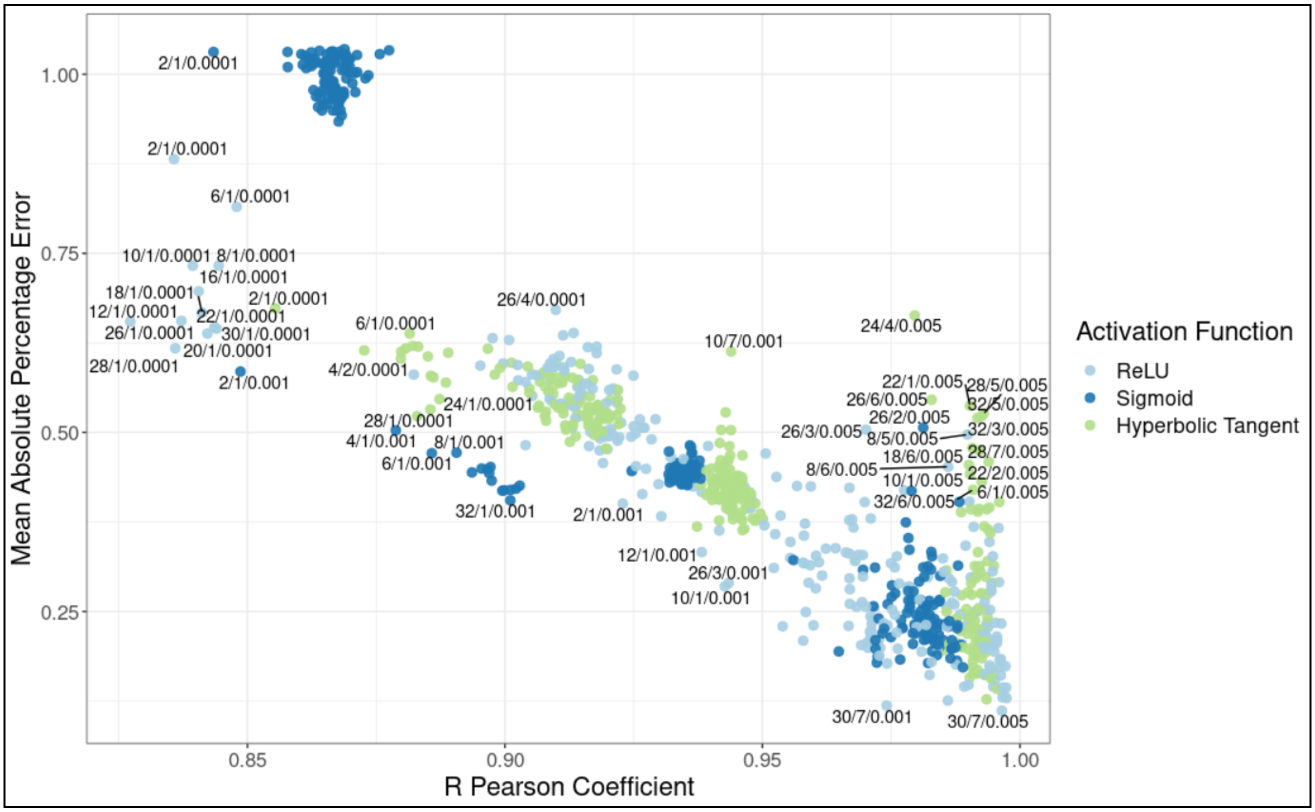


**Supplementary Fig. 28**. Hyperparameter search for stem circumference BLUP prediction in experimental group 1 (EG1) for the population GT1 x RRIM701. Each point represents a different combination of hyperparameters, considering (a) neurons in the first hidden layer, (b) neurons in the second hidden layer, (c) the learning rate, and (d) the activation function. These combinations were evaluated in the development set considering the mean absolute percentage error (MAPE) and the R Pearson coefficient.

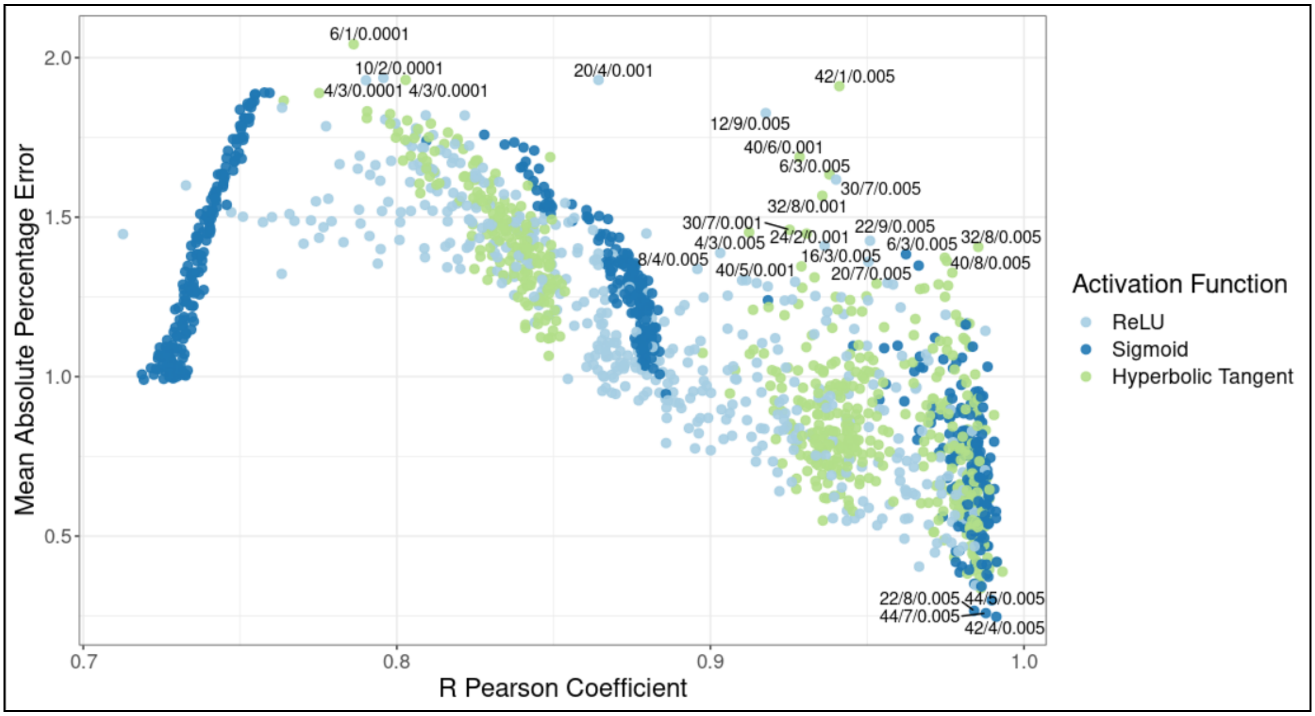


**Supplementary Fig. 29**. Hyperparameter search for stem circumference BLUP prediction in experimental group 1 (EG1) for the population PR255 x PB217. Each point represents a different combination of hyperparameters, considering (a) neurons in the first hidden layer, (b) neurons in the second hidden layer, (c) the learning rate, and (d) the activation function. These combinations were evaluated in the development set considering the mean absolute percentage error (MAPE) and the R Pearson coefficient.

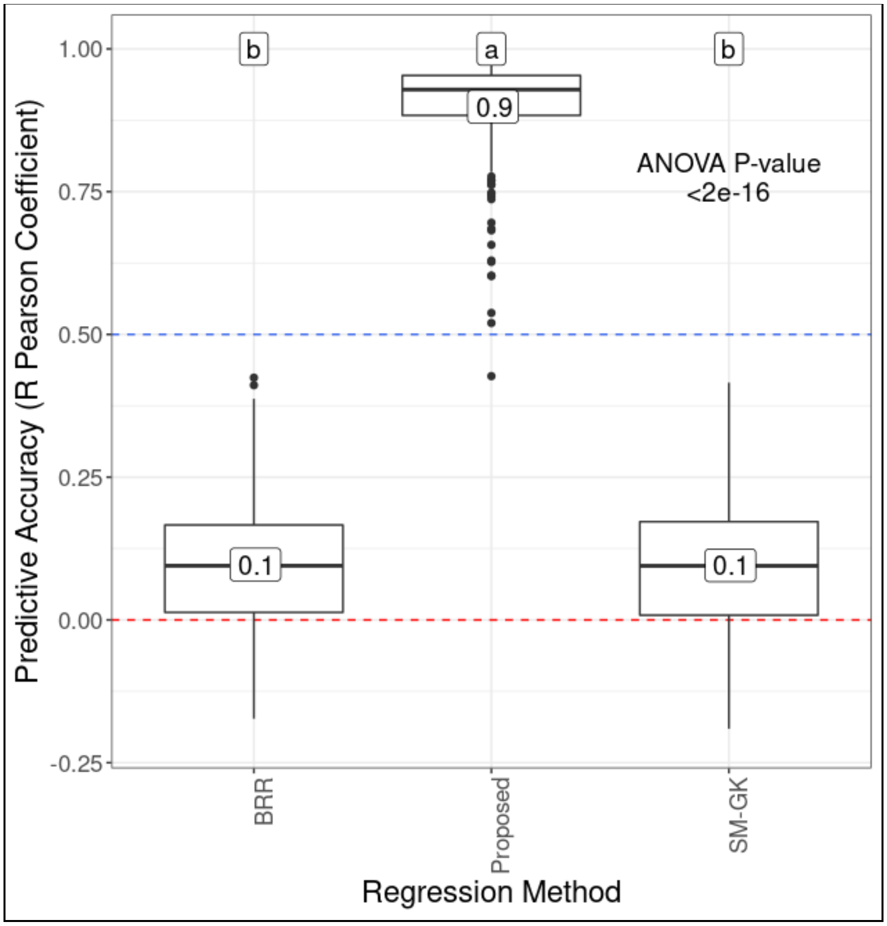


**Supplementary Fig. 30**. Predictive accuracies for stem circumference BLUP prediction in experimental group 1 (EG1) and the population GT1 x PB235 considering a 4-fold cross validation scheme (repeated 50 times), the proposed approach and traditional genomic prediction methods (a single-environment model with a nonlinear Gaussian kernel (SM-GK) and Bayesian ridge regression (BRR)). The letters at the top indicate the results from Tukey's multiple comparison test.

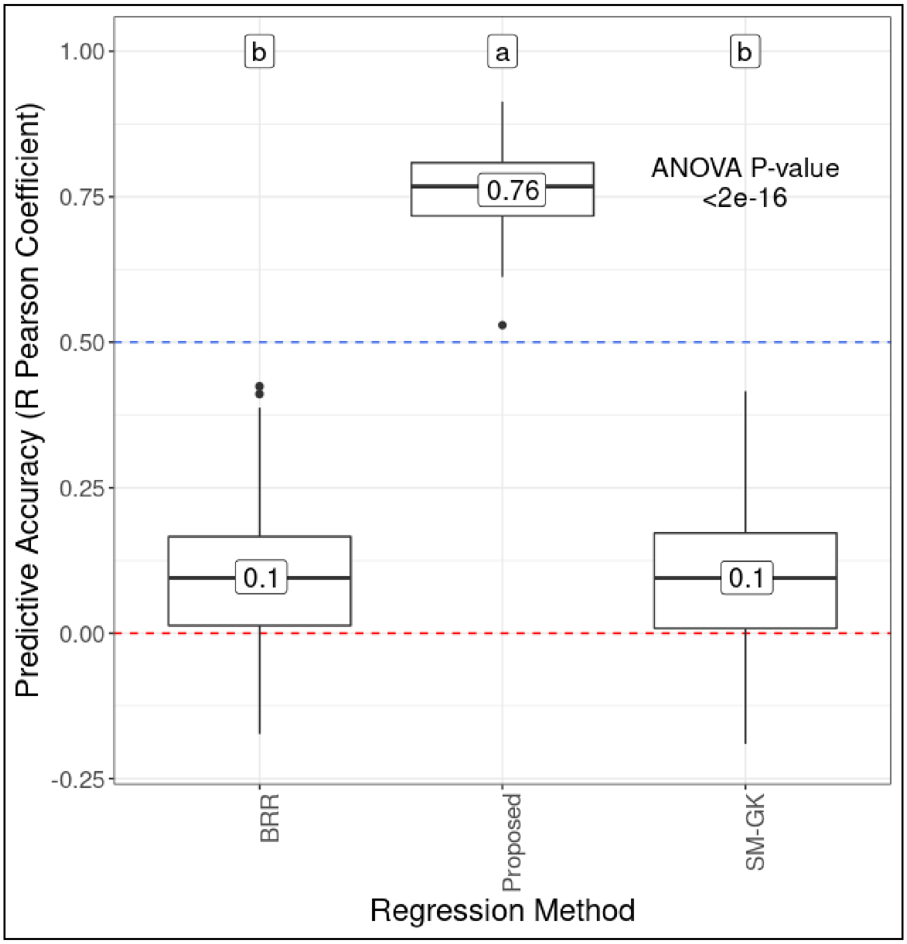


**Supplementary Fig. 31**. Predictive accuracies for stem circumference BLUP prediction in experimental group 1 (EG1) and the population GT1 x RRIM701 considering a 4-fold cross validation scheme (repeated 50 times), the proposed approach and traditional genomic prediction methods (a single-environment model with a nonlinear Gaussian kernel (SM-GK) and Bayesian ridge regression (BRR)). The letters at the top indicate the results from Tukey's multiple comparison test.

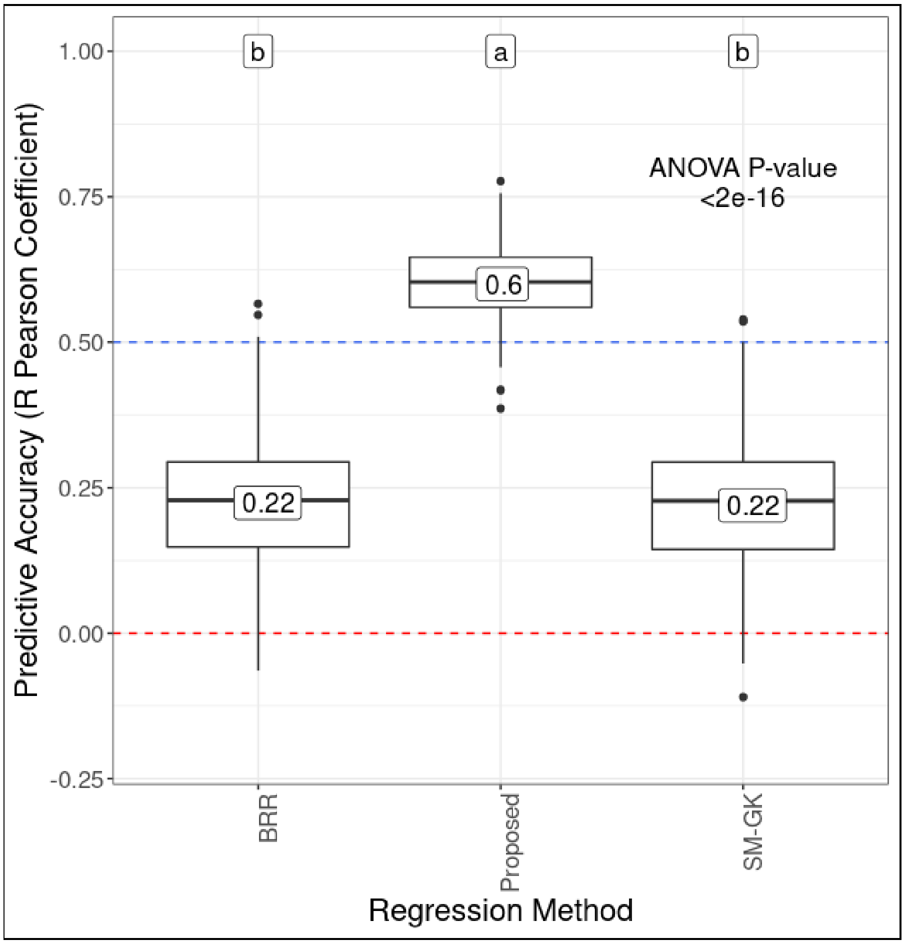


**Supplementary Fig. 32**. Predictive accuracies for stem circumference BLUP prediction in experimental group 1 (EG1) and the population PR255 x PB217 considering a 4-fold cross validation scheme (repeated 50 times), the proposed approach and traditional genomic prediction methods (a single-environment model with a nonlinear Gaussian kernel (SM-GK) and Bayesian ridge regression (BRR)). The letters at the top indicate the results from Tukey's multiple comparison test.
